# Mechanistic dissection of *Candidatus* Liberibacter Triggered Chronic Immune Disease

**DOI:** 10.1101/2025.05.21.654963

**Authors:** Xiaoen Huang, Wenxiu Ma, Wenting Wang, Yu Feng, Tirtha Lamichhane, Jin Xu, Sheo Shankar Pandey, Diann S Achor, Jinyun Li, Yuanchun Wang, Javier L. Dalmendray, Zhuyuan Hu, Camila Ribeiro, Ning Zhang, Madhurababu Kunta, Allison K. Hansen, Nian Wang

**Author notes:** WW, WM, XH, YF, TL, and JX contributed equally to this study.

## Abstract

Immunity is generally considered critical for plant health against pathogen infection. Citrus Huanglongbing (HLB) caused by the phloem colonizing bacterial pathogen *Candidatus* Liberibacter asiaticus (CLas) was suggested to be a pathogen triggered chronic immune disease. However, the genetic evidence and mechanistic understanding for such a disease model is lacking. Here, we show CLas triggers phloem cell death, reactive oxygen species (ROS) production and callose deposition in photosynthesis tissues, but little or none in non-photosynthesis tissues of citrus. We further demonstrate that CLas triggers ROS production in chloroplasts. Overexpression of *flavodoxin* (*Fld*), an electron shuttle, in chloroplasts reduced ROS production, cell death and HLB symptoms, but not phloem callose deposition induced by CLas. Knockout, silencing, and overexpression of phloem callose synthase genes *CsCalS7a*, *CsCalS7b*, or *CsCalS7c* demonstrated that phloem callose deposition also caused phloem cell death. Trunk injection with 2-deoxy-d-glucose, a callose deposition inhibitor, reduced fruit drop and increased fruit yield of HLB symptomatic trees in field trials. Using tomato-*Ca*. L. psyllaurous (Lpsy) as a model, knockout of *Eds1* and *Pad4* but not *RbohB*, *Bik1* and *Sobir1* abolished ROS production, phloem callose deposition, cell death and disease damages caused by Lpsy. This study provides genetic evidence for *Ca*. Liberibacter-triggered immune disease and reveals that tuning plant immune responses converts *Ca*. Liberibacter into a benign endophyte, providing a promising strategy for precision breeding to enhance resistance/tolerance against HLB.

---

Immunity not only plays critical roles in protecting animals and humans from infection by pathogens, but also is causal in the development of many diseases including inflammatory bowel disease and sepsis ^1^. In contrast, the possible involvement of immunity in the development of plant diseases is largely ignored. A previous study suggested that citrus Huanglongbing (HLB), caused by the phloem colonizing bacterial pathogen *Candidatus* Liberibacter asiaticus (CLas), is a pathogen triggered chronic immune disease involving reactive oxygen species (ROS) production, phloem callose deposition, and phloem cell death ^2^. However, the genetic evidence and mechanistic understanding for such a disease model is lacking. *Candidatus* Liberibacter causes severe diseases on many crops including citrus, solanaceous (e.g., tomato, and potato) and apiaceous crops. Specifically, HLB is the most devastating citrus disease worldwide. Typical HLB symptoms include blotchy mottles in leaves, leaf yellowing, leaf hardening, corky veins, upright and small leaves, root decay, leaf nutrient deficiency, branch die back, leaf and fruit drop, tree decline and tree death ^3^. CLas has not been cultured in artificial media in the absence of other bacteria ^4^ and genetic manipulation of CLas remains impossible. Multiple studies have found that phloem callose deposition around sieve pores, phloem cell death and starch accumulation occur in HLB diseased plants even though their involvement in HLB disease development remains largely unknown ^5–7^. A large amount of work has been conducted to identify the virulence factors of Clas such as Sec-dependent effectors (SDEs). So far, no such virulence factors of CLas have been shown to cause HLB symptoms and the identified CLas effectors are involved in either inducing plant immune responses such as callose deposition and reactive oxygen species (ROS) production or suppressing plant immune responses such as the suppression of programmed cell death that is induced by elicitors ^8–12^. One recent breakthrough study has revealed that several citrus relatives resist CLas via the PUB21-MYC2 circuit ^13^. In this study, we demonstrated that *Ca.* Liberibacter triggers a chronic immune disease through the Eds1 and Pad4 mediated induction of both chloroplastic ROS production and phloem callose deposition. This provides genetic evidence for HLB as a pathogen-triggered immune disease and elucidates targets for genetic manipulation for disease resistance breeding against HLB.

## CLas triggers phloem cell death, ROS production and callose deposition in photosynthesis tissues but not or significantly less in non-photosynthesis tissues of citrus

Phloem cell death in leaves that is triggered by CLas positively correlates with CLas titers and HLB symptoms ^2^. Here, we investigated whether CLas triggers phloem cell death in other tissues of *Citrus sinensis* as well using transmission electron microscopy (TEM), which enables visualization of both phloem cell death and callose deposition around sieve pores. In addition to leaves, phloem cell death and callose deposition were also observed in CLas-infected fruit albedo, columella, and bark tissues of small branches and the trunks of trees, but not in those of healthy plants (Fig. 1a). Phloem cell death and callose deposition were not observed in either heathy or CLas-infected seed coats (Fig. 1a), consistent with a previous study ^14^. Similar callose deposition was observed between healthy and Clas colonized roots (Fig. 1a). Dead cells were observed in CLas-infected roots, but much less abundantly than in leaves. To get a better picture of the cell death in root tissues that was triggered by CLas, we conducted TEM analysis of newly generated roots of HLB symptomatic and healthy *C. sinensis* cv. Valencia on Cleopatra mandarin rootstock. Phloem callose deposition was similar in both CLas infected and uninfected roots (Fig. S1a). No obvious cell death was observed in newly generated roots of 1–4-week-old. However, cell death was observed in 5-week-old roots. Unlike leaves which exhibited only phloem cell death, ubiquitous death of all cell types was seen in most root samples, suggesting that it might result from carbohydrate starvation ^15^ owing to Clas-triggered phloem malfunction. In support of this model, starch accumulation was found to be significantly reduced in the CLas-infected samples compared to the healthy plants in all time points (Fig. S1b). Furthermore, ion leakage assays of root samples of CLas-infected Valencia scions grafted on Kuharske rootstocks and Sugar Belle, Valencia, and Tango scions grafted on US942 rootstocks, exhibited root cell death regardless of their infection status. This supports the hypothesis that increased root cell death in HLB symptomatic trees is driven by carbon starvation instead of phloem cell death as in leaves (Fig. S2).

**Fig. 1:**
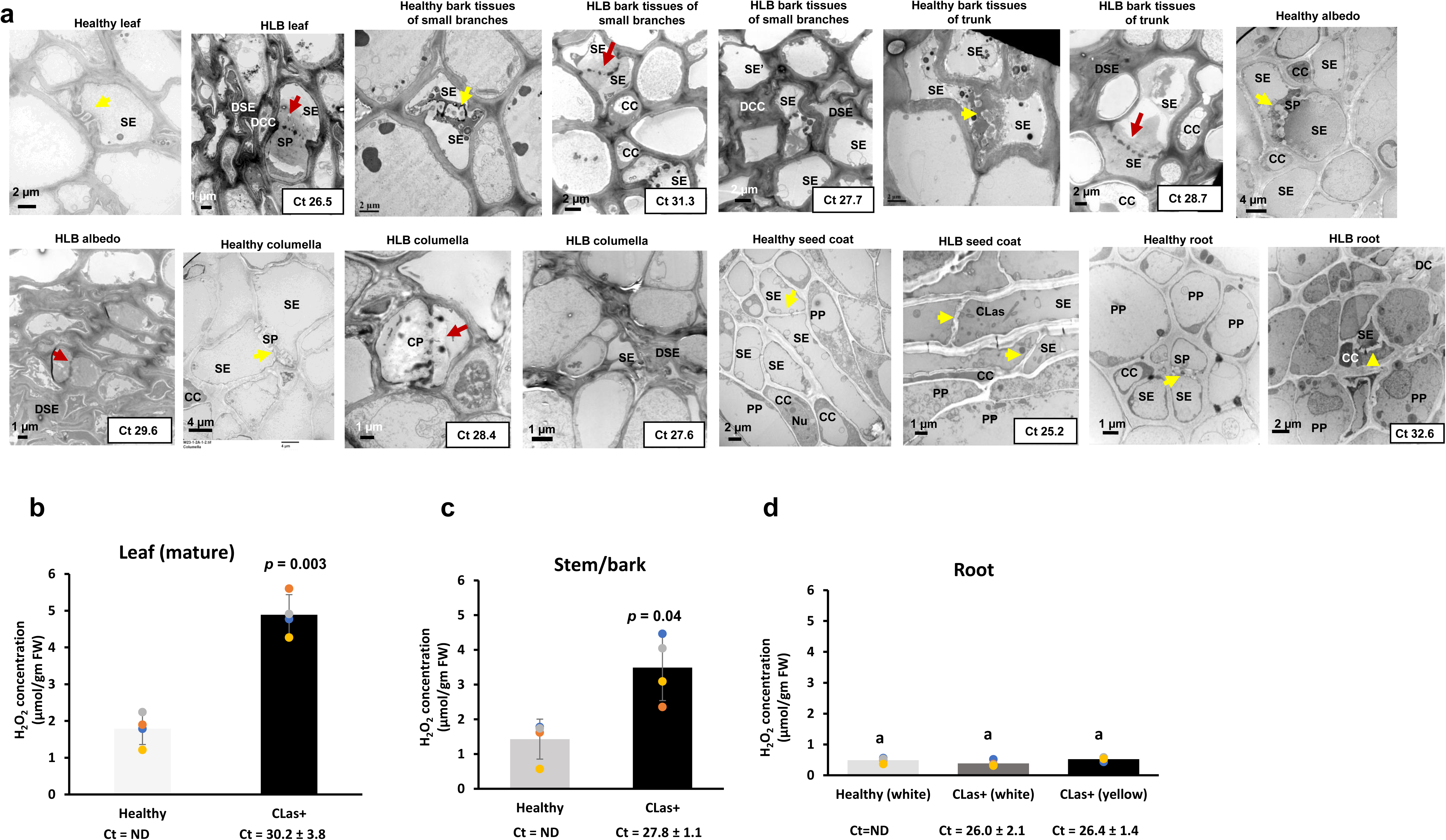
Phloem cell death, callose deposition and ROS production in different citrus tissues. (**a**) Transmission electron microscopy (TEM) analysis of *Citrus sinensis* on the Swingle citrumelo rootstock with or without CLas infection. PC: parenchyma cell; SP: sieve plate; CC: companion cell; SE: sieve element; PP: phloem protein; DCC: dead CC; DSE: dead SE, CLas: live CLas cells; DC: dead cells; Nu: nucleus. Yellow arrows indicate sieve pores without excessive callose deposition. Red arrows indicate sieve pores with excessive callose deposition. (**b-d**) H_2_O_2_ concentrations in different tissues of CLas infected sweet orange (*Citrus sinensis* cv Valencia) on the Kuharske rootstock. *P* value represents the significant difference (n = 4, student’s t-test, *P* < 0.05) compared to the healthy control. Different letters indicate a significant difference. The corresponding Ct value represents CLas concentrations in the infected citrus tissues with low Ct values indicating high CLas titers. ND = Not detected.

ROS are known to contribute to phloem cell death triggered by CLas ^2^. Given this, we investigated ROS production in leaf, stem and root tissues in infected plants. The first two are photosynthetic tissues while the latter is non-photosynthetic tissues. CLas infection caused significantly higher ROS levels to develop in leaves and bark tissues of stems compared to healthy controls, but not in roots (Fig. 1b-d). Additionally, it was reported that CLas does not increase ROS production in seed coats ^14^. Together, these results demonstrate that CLas triggers phloem cell death, ROS production and callose deposition in tissues capable of photosynthesis such as leaves and the bark tissues of branches and trunks, but much less in tissues that do not contain chloroplasts such as roots and seed coats ^2,14^. Thus, chloroplasts and photosynthesis appear to play critical roles in the immune-mediated development of HLB disease.

## CLas triggers ROS production in chloroplasts

Chloroplasts are known to be involved in ROS production in response to pathogen infection ^16–18^. In addition, chloroplasts play a critical role in regulating programmed cell death (PCD) in response to pathogen infection ^19^. We therefore tested whether CLas triggers ROS production in chloroplasts using CLas-infected and healthy leaves of *C. sinensis* cv. Valencia. Total ROS were detected using 2’,7’-dichlorodihydrofluorescein diacetate (DCFH-DA) staining ^2^. Hydrogen peroxide (H_2_O_2_) was detected via DAB staining. Superoxide (O_2_^·^−) was detected by staining with nitro blue tetrazolium chloride (NBT) ^20^. ROS were found in the chloroplasts of CLas infected citrus leaves but were absent in healthy leaves (Fig. S3). Both DAB and NBT staining showed induced production of H_2_O_2_ and O_2_^·−^ in the chloroplasts located in vein tissue, respectively (Fig. 2 a, b). We further investigated singlet oxygen (^1^O_2_) in the vein tissues of CLas-infected and healthy leaves by using singlet oxygen sensor green reagent (SOSG), which is highly responsive only to ^1^O_2_. We clearly observed the induction and localization of ^1^O_2_ only in the chloroplasts of CLas-infected veins, while it was absent in healthy tissues (Fig. 2c).

**Figure 2:**
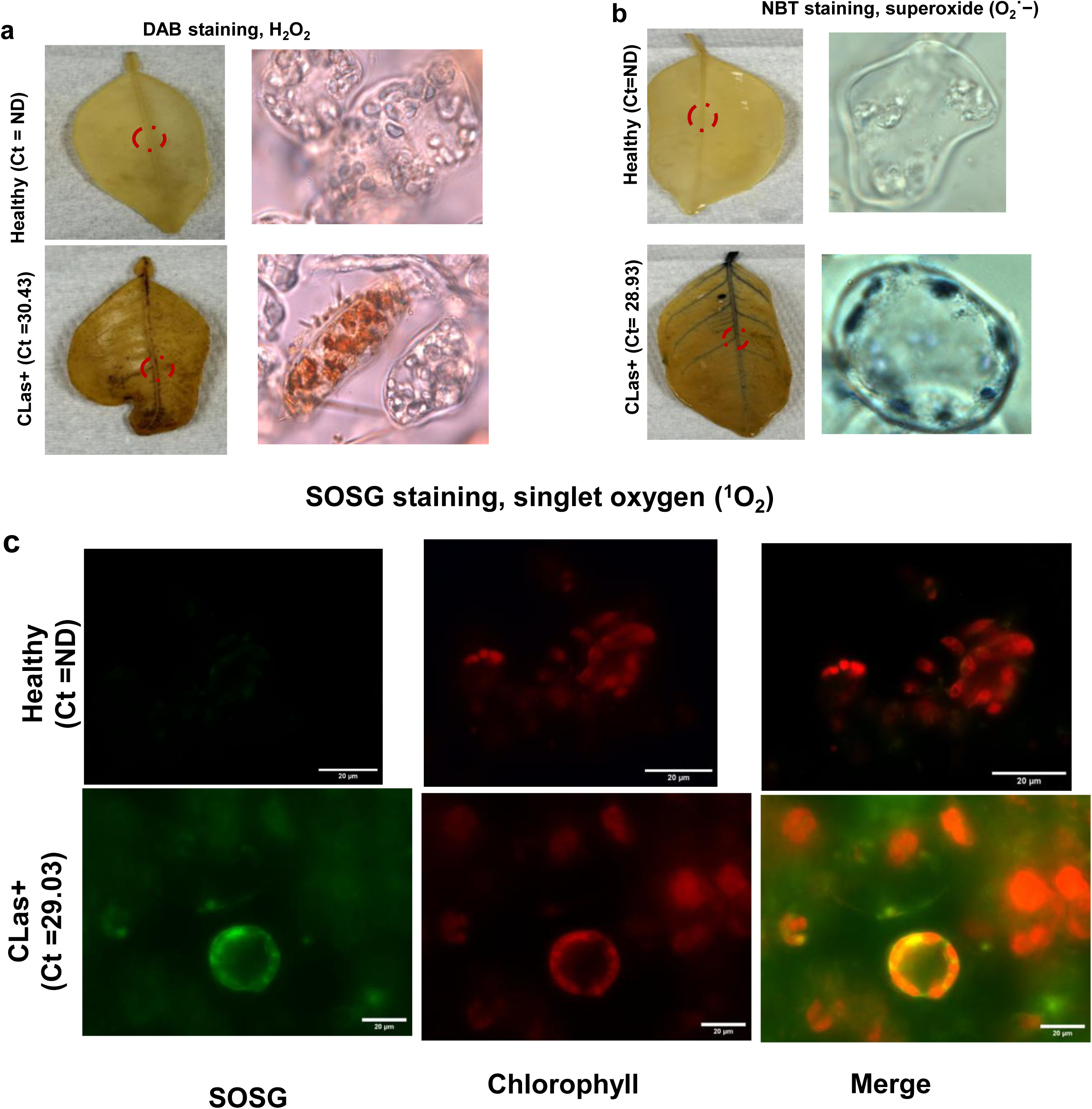
Detection of hydrogen peroxide (H_2_O_2_), superoxide (O2^·−^), and singlet oxygen (^1^O2) in the midvein of CLas infected and healthy *C. sinensis* leaves. (**a**) Detection of H_2_O_2_ in the midvein using the DAB staining method. After DAB staining and rinsing, samples were cut from midrib region (indicated in red dotted circle) to make a thin slice for microscopy. The images were taken under bright field microscopy (600X) power magnification. (**b**) Detection of O_2_ ^·−^ using the NBT staining method. NBT is reduced by free oxygen radical and forms a blue-black formazan compound indicates its presence. After NBT staining and rinsing, samples were cut (indicated in red dotted circle) to make a thin slice for microscopy. The images were taken under bright field microscopy (600X) power magnification. (**c**) Detection of ^1^O_2_ using singlet oxygen sensor green reagent (SOSG). Protoplast cells were isolated from CLas infected and healthy Valencia mature leaves by enzymatic digestion with 1% cellulase RS and 1% macerozyme R-10. After 16 hours of enzyme digestion, protoplast cells were isolated and treated with 10 µM of SOSG then incubated for 2 hours in darkness at room temperature. Treated protoplast cells were observed by fluorescence microscopy. Green signal indicates singlet oxygen detection (SOSG) and red indicates the chloroplast autofluorescence. Merge = chlorophyll + SOSG. This experiment is repeated at least twice with similar results. CLas titers were quantified by qPCR.

## CLas triggers global upregulation of immunity and downregulation of photosynthesis in leaves, but not roots, likely via the ETI pathway

Chloroplastic ROS accumulation is known to be triggered in PAMP-triggered immunity (PTI), and effector-triggered immunity (ETI) in plant responses to various microbes ^18,21^. Su et al. ^16^ reported that NLR-mediated ETI induces global inhibition of photosynthesis and generation of ROS in chloroplasts, whereas de Torres Zabala et al. ^22^ showed that PAMPs induce ROS production in chloroplasts, but had no effect on photosynthesis. We therefore investigated whether CLas induces immunity and reduces photosynthesis by conducting Gene Ontology (GO) term enrichment analysis of differentially expressed genes (DEGs) in leaf and root tissues of HLB symptomatic and healthy citrus plants. For this analysis, we accessed 8 microarray and 21 RNA-seq data sets from previous studies (Data S1). The pathways up-regulated by CLas in leaves are enriched for functions such as calcium transport, defense response, response to stress, kinase, and protein phosphorylation. In contrast, pathways primarily associated with photosynthesis including chlorophyll biosynthesis, photosystem I and II, thylakoid, and light reaction are down-regulated in Clas-infected tissues (Fig. S4). The global upregulation of immune responses alongside the downregulation of photosynthesis mirror the effects of ETI on immune responses and photosynthesis ^16^, but differs from the patterns observed in PTI ^22^. Both ETI and PTI have been shown to activate the mitogen-activated protein kinases (MPKs) MPK3/MPK6 ^23^. In particular, the prolonged activation of MPK3/MPK6 during ETI suppresses photosynthesis, leading to increased ROS in chloroplasts ^16^. CLas infection induced the expression of both *MPK3* and *MPK6* compared to that in healthy control plants and CLas also induced phosphorylation of MPK3 and MPK6, which was absent in healthy plants (Fig. S5).

In contrast to leaves, GO term enrichment analysis of root tissue of HLB symptomatic plants compared to healthy controls did not reveal any upregulation of immune-related pathways or the down-regulation of photosynthesis pathways (Fig. S6). The different patterns of expression of genes related to immunity and photosynthesis in the leaf and root tissues in response to CLas infection is consistent with significantly reduced ROS accumulation, callose deposition and phloem cell death in roots compared to leaves in response to CLas infection (Fig. 1, Figs.S1-2).

## Mitigation of chloroplastic ROS reduces HLB symptoms, starch accumulation, and ion leakage, but does not affect phloem callose deposition induced by CLas

Since chloroplastic ROS production appears to play critical roles in HLB symptom development, we further investigated its role by reducing chloroplastic ROS via overexpressing Flavodoxin (Fld) in chloroplasts. Flavodoxin is an electron shuttle that is present in both prokaryotes and algae, whose overexpression in chloroplasts decreases chloroplastic ROS production and cell death caused by ETI ^16,17^. Additionally, ROS are known to positively regulate callose deposition ^24^. We hypothesized that overexpression of Fld reduces both chloroplastic ROS and phloem callose deposition, thus decreasing HLB symptoms. We overexpressed *35S::TP* (*chloroplast transit peptide*)*-Fld* in *C. sinensis*. The localization of the chloroplast-targeted Fld in chloroplasts ^25^ was confirmed by confocal microscopy analysis of transient expression of *35S::TP-Fld* with a mVenus tag in *N. benthamiana* leaves (Fig. 3a-b). The *35S::TP-Fld* transgenic *C. sinensis* infected with CLas exhibited only minor symptoms typical of HLB including leaf yellowing whereas wild type plants had more severe HLB symptoms including smaller leaves, leaf hardening and corky veins (Fig. 3c). *Fld* overexpression reduced ROS levels (Fig. 3d), starch accumulation (Fig. 3e), and ion leakage (Fig. 3f), but not callose deposition (Fig. 3g) in response to Clas infection compared to the wild type. The reduction of Clas-triggered ROS levels (Fig. 3d) and cell death as revealed by ion leakage assays (Fig. 3f) in the *Fld* transgenic lines compared to wild type plants is consistent with a previous report that CLas triggered ROS can directly cause phloem cell death ^2^. The slight leaf yellowing of Clas-infected *Fld* transgenic lines probably resulted from starch accumulation, which was much reduced in the *Fld* transgenic lines compared to the high levels seen in CLas infected wild type plants (Fig. 3e). The starch accumulation in the *Fld* transgenic lines was potentially caused by phloem callose deposition (Fig. 3g). Additionally, our data indicates that CLas-induced phloem callose deposition occurs independently of ROS production, different from our original hypothesis.

**Fig. 3:**
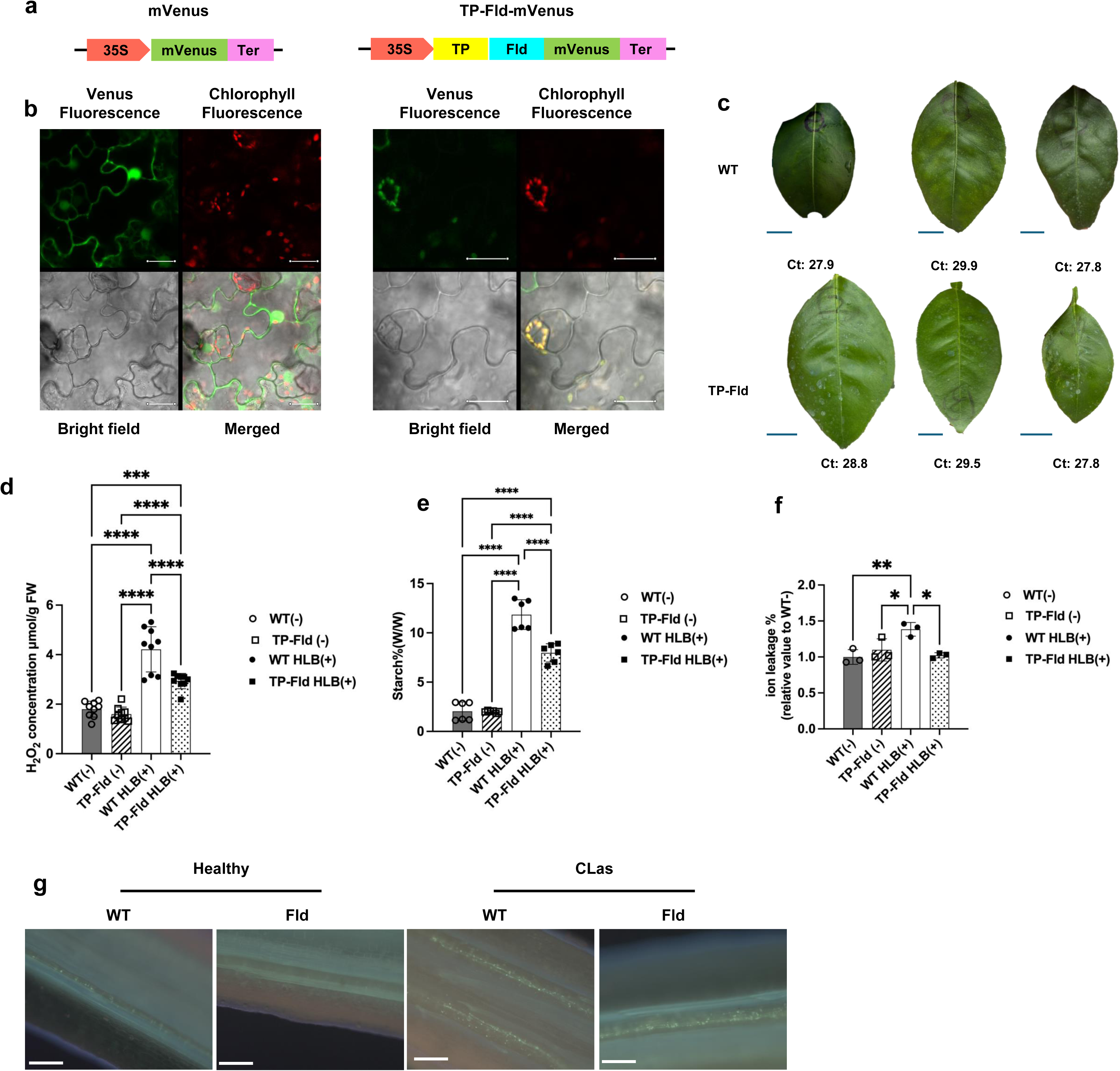
Overexpression of Fld in the chloroplast scavenges chloroplastic ROS production, reduces starch accumulation, ion leakage, and HLB symptoms, but does not affect phloem callose deposition induced by CLas. (**a**) Construct for transient expression of chloroplast-targeted *Fld* in *Nicotiana benthamiana* via agroinfiltration (right). The schematic diagram to the left is the negative control. TP: chloroplast transit peptide. Fld: flavodoxin. CaMV 35S: the Cauliflower Mosaic Virus 35S promoter. Ter: NOS terminator. mVenus: mVenus fluorescent protein. (**b**) Confocal analysis of the localization of TP-Fld-mVenus in *N. benthamiana.* Left: 35S::mVenus transiently expressed in *N. benthamiana* leaf. Right: 35S::TP-Fld-mVenus transiently expressed in *N. benthamiana* leaf. All scale bars, 20 μm. (**c**) Representative leaf symptoms of *35S::TP-Fld* transgenic *C. sinensis* cv. Hamlin and wild-type (WT) plants six months after graft-inoculation with CLas. Scale bars, 1 cm. (**d**) Quantification of H₂O₂ in leaves of TP-Fld transgenic and wild-plants in (c). Data shown are mean ± SEM, *n* = 3. (**e**) Starch content in leaves of (c). (**f**) Ion leakage assay of (c). Data shown are mean ± SEM, *n* = 9. Experiments were repeated two times with similar results. One-way analysis of variance (ANOVA) was conducted, followed by Tukey’s test at a significance level of 0.05. *, **, and ***,**** indicate *P* < 0.05, 0.01, 0.001,0.0001, respectively. The original data were processed and visualized using GraphPad Prism 8.0 software. (**g**) Callose deposition analysis of leaves in (c). Scale bars are 100 μm.

Next, we investigated the contribution of phloem callose deposition in HLB symptom development by altering the expression of phloem callose synthase 7 (*CalS7*), which is responsible for phloem sieve element callose deposition in *Arabidopsis* ^26,27^, by mutation, silencing or overexpression.

## Three *AtCalS7* homologs were expressed in the phloem tissues in *C. sinensis*

There are three callose synthase 7 genes in *C. sinensis* (Cs_ont_7g015100 (CsCalS7a, 1922 aa), Cs_ont_7g015150 (CsCalS7b, 1910 aa) and Cs_ont_7g014700 (CsCalS7c, 1904aa)) that are homologs of AtCalS7 in *Arabidopsis.* They share 1433/1923(73.61%), 1353/1905(71.02%) and 1354/1930 (70.16%) amino acid identity, respectively, with AtCalS7, based on BLASTp. The three domains (Vta1, FKS1 and Glucan synthase) are highly conserved in all three proteins (Fig. S8). Similar to AtCalS7, CsCalS7a, CsCalS7b and CsCalS7c have 18 transmembrane motifs based on analysis with TMHMM version 2.0 ^28^, with 8 located in the N terminus and 10 in the C terminus (Fig. S8c).

To test whether *CsCalS7a*, *CsCalS7b* and *CsCalS7c* are indeed expressed in phloem tissues, we generated transgenic *C. sinensis* plants expressing erGFP driven by *CsCalS7a, b* or *c* promoters. erGFP driven by the 35S promoter was used as a control to show constitutive expression patterns. As expected, 35S-erGFP exhibited ubiquitous expression whereas erGFP driven by *CsCalS7a, b* or *c* promoters was only observed in the phloem tissues of transgenic roots (Fig. S9a). Specifically, GFP signals were detected in the regions where phloem callose accumulates, as stained by aniline blue, indicating that *CsCalS7a, CsCalS7b* and *CsCalS7c* were expressed in the phloem. The phloem-specific expression of *CsCalS7a, b* or *c* was also observed by analyzing longitudinal sections of shoots (Fig. S9b). erGFP signals were detected in sieve elements, indicating that *CsCalS7a*, *b* and *c* are expressed in these cells.

## Loss of function of *CsCalS7a* and *CsCalS7b* but not *CsCalS7c* affects phloem callose deposition, cell death, starch accumulation, and causes HLB-like phenotypes

CalS7 is known to be involved in callose synthesis in the phloem and sieve pore formation ^26,27^. We therefore investigated whether *CsCalS7a*, *CsCalS7b* and *CsCalS7c* have such functions by conducting genome editing of *CsCalS7a*, and *CsCalS7c* and RNAi silencing of *CsCalS7b* in *C. sinensis* cv. Hamlin. For mutation of *CsCalS7a* and *CsCalS7c*, a multiplex CRISPR/Cas9 construct was used ^29^, and two specific guide RNAs (sgRNA) for each gene were selected (Fig. S10). Seven biallelic *CsCalS7a* edited lines and ten biallelic *CsCalS7c* edited lines were generated. The genotypes of edited lines were confirmed by whole genome sequencing (Fig. S11). To investigate any off-target mutations, whole genome sequencing of representative *cals7a* and *cals7c* lines was conducted. In three sequenced *cals7a* lines, off-target mutation of *CsCalS7c* was detected. However, the mutation of *CsCalS7c* was heterozygous indicating *CsCalS7c* remained functional (Fig. S11b). In *cals7c* mutant lines, no off-target mutations were detected. For RNAi silencing of *CsCalS7b* (Fig. S12), based on GFP screening and qRT-PCR assays, 6 *CsCalS7b-*silenced lines were generated. The relative expression of *CsCalS7b* in the 6 RNAi plants was significantly decreased compared with the empty vector (EV) control (Fig. S12d-f). Even though *CsCalS7a*, *CsCalS7b* and *CsCalS7c* share 93.6% CDS DNA sequence similarity, only the expression level of the targeted *CsCalS7b* decreased whereas that of *CsCalS7a* and *CsCalS7c* had no significant changes in expression (Fig. S12c-e), indicating that expression of *CsCalS7b* was successfully silenced in the RNAi lines.

We originally hypothesized that knockout or gene silencing of *CalS7* genes would reduce phloem callose deposition, thus decreasing HLB symptoms. Interestingly, both *cals7a* mutants and *CsCalS7b-RNAi* plants exhibited HLB-like symptoms including leaf yellowing, smaller leaves, corky veins, and reduced plant growth measured as plant height and leaf numbers and size (Fig. 4a-d; Fig. S13). In contrast, *cals7c* mutants were similar to healthy wild type *C. sinensis* (Fig. 4c, d; Fig. S13c).

**Fig. 4:**
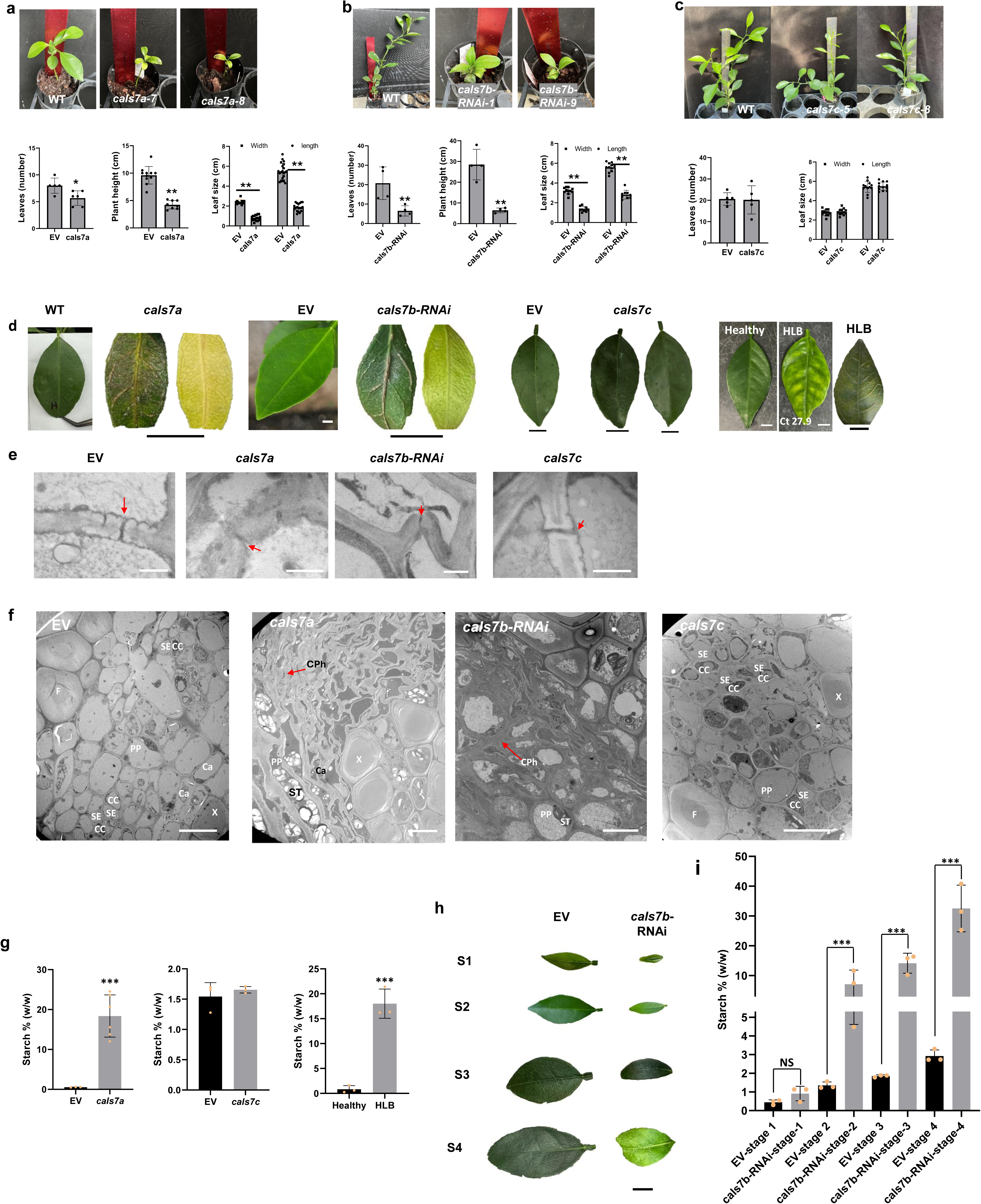
Phenotypes of loss of function mutants of *CsCalS7* genes in *C. sinensis* cv. Hamlin. (**a**) Three-month-old *CsCalS7a* gene edited lines. (**b**) Nine-month-old *CsCalS7b* silencing lines. (**c**) Nine-month-old *CsCalS7c* gene edited lines. Measurement of leaf number, plant height and leaf size were done for a-c. Student’s *t* test was used for statistical analysis. (**d**) Leaves of *CsCalS7a* edited lines and *CsCalS7b* silencing lines but not *CsCalS7c* edited lines showing HLB-like phenotypes. Healthy and HLB symptomatic leaf of *C. sinensis* cv. Hamlin were included as controls. Ct value of CLas titer quantified by qPCR. Scale bars are 1 cm. (e,f) TEM analysis of the midveins of loss of function mutants of *CalS7a*, *b*, and *c* of *C. sinensis* cv. Hamlin. (**e**) Sieve pores in midrib of young shoots (at stage 1 in h). Red arrows indicate sieve pore. Bars are 1 μm. (**f**) Cross-sections of mature leaf midrib. Scale bars are 10 μm. Phloem tissues of empty vector (EV) transgenic sweet orange cv. Hamlin, genome edited *cals7a* mutant, *cals7b-RNAi* transgenic plants and genome edited *cals7c* mutant. PP, phloem parenchyma; SE, sieve element; CC, companion cell; X, xylem; F, fiber; Ca, cambium; CPh, collapsed phloem cells; ST, starch. (g-i) Loss of function of *CalS7a* and *CalS7b*, but not *CalS7c* causes starch accumulation in *C. sinensis* cv. Hamlin. (**g**) Starch accumulation in *cals7a* and *cals7c* mutant and HLB symptomatic leaves. (**h**) Different stages of *cals7b-RNAi* and wild type plant leaves. Bar is 1 cm. (**i**) Starch accumulation in different stages (h). * indicates *P* < 0.05 ** indicates *P* < 0.01 and *** indicates *P* < 0.001. Student’s *t* test was used for statistical analysis.

The *cals7a* mutant and *CsCalS7b-RNAi* plants exhibited significantly reduced callose accumulation in the phloem tissues of leaf midveins compared to the wild type *C. sinensis*, whereas the *cals7c* mutant was similar as the wild type (Fig. S14). In accord with callose accumulation, TEM analysis showed that functional sieve pores could be seen in the wild type and *cals7c* mutant, but not in the *cals7a* mutant and *CsCalS7b-RNAi* plants (Fig. 4e). It appeared that sieve pores sizes in the *cals7a* mutant and *CsCalS7b-RNAi* plants were much smaller than that of the wild type and the *cals7c* mutant or not fully developed (Fig. 4e; Fig. S15), mimicking the reduced sieve pores of HLB symptomatic citrus plants owing to callose deposition triggered by CLas ^30,31^. Additionally, plasmodesmata were similar in size and number in the *cals7a* mutant, *cals7c* mutant, and *CsCalS7b-RNAi* plants as in the wild type, indicating they are not involved in plasmodesmata formation (Fig. S16). Together, our data suggested that *CsCalS7a* and *CsCalS7b* but not *CsCalS7c* are required for phloem callose synthesis and sieve pore formation.

Importantly, phloem cell death was observed in the midveins of leaves of both the *cals7a* mutant and *CsCalS7b-RNAi* plants (Fig. 4f; Fig. S17). Phloem cell death was not observed in the healthy wild type plants and the *cals7c* mutant (Fig. 4f; Fig. S17). Phloem cell death also occurred in the midvein during early stages of leaf development of both the *cals7a* mutant and *CsCalS7b-RNAi* plants (Fig. S18). Phloem cell death has been suggested to be a key contributor to HLB symptom development and positively correlates with CLas titers ^2^. Thus, phloem cell death of both the *cals7a* mutant and *CsCalS7b-RNAi* plants likely explains their HLB-like symptoms (Fig. 4a-d, Fig. S13). Interestingly, phloem cell death was not observed in *CalS7* mutants of *Arabidopsis* ^26,27^ even though they also have reduced phloem callose deposition and sieve pore sizes as the citrus *cals7a* mutant and *CsCalS7b-RNAi* plants.

Phloem malfunction is known to cause starch accumulation in leaf chloroplasts. For instance, trunk girdling results in such starch accumulation ^32^. Starch accumulation was significantly higher in the chloroplasts in the *cals7a* mutants and *CsCalS7b-RNAi* plants than in the healthy wild type and *cals7c* mutant (Fig. 4g-i). Starch accumulation in chloroplasts results from the imbalance of sugar production from photosynthesis and sugar transported out via the phloem. Starch accumulation in chloroplasts that is triggered by CLas has been suggested to cause chloroplast degradation, resulting in leaf yellowing/blotchy mottle symptoms, typical of HLB ^6^. Reducing sunlight by 80% on *CsCalS7b-RNAi* plants for one month reduced starch accumulation and prevented leaves from turning yellowing (Fig. S19a, b). This is consistent with a report that shade treatment of HLB symptomatic trees reduces starch accumulation and reduces obvious symptoms of HLB ^33^. However, the shade treatment did not prevent phloem cell death in leaf midveins of *CsCalS7b-RNAi* plants (Fig. S19c), indicating starch accumulation does not cause phloem cell death. Instead, phloem cell death/phloem malfunction is responsible for starch accumulation in the leaf and subsequent leaf yellowing/blotchy mottle symptoms. It is worth noting that phloem cell death was also observed in the roots of the *CsCalS7b-RNAi* plant (Fig. S20).

Phloem cell death caused by CLas triggers phloem regeneration ^34^ including hypertrophy and hyperplasia of phloem tissues, which leads to vein ruptures. Light microscopic analysis showed more phloem regeneration in the leaf midvein of the *cals7a* mutant and *CalS7b-RNAi* plants than the wild type plants (Fig. S21), similar to that seen in HLB symptomatic plants (Figs. S2 & S21c). Consistent with this observation, transcriptomic data reveal that genes contributing to phloem differentiation are highly expressed in *cals7a* and *cals7b-RNAi* as compared to healthy wild type plants (Fig. S21d). Taken together, we have found that loss of function of *CsCalS7a* and *CsCalS7b* causes phloem blockage, similar to that seen in HLB, and also demonstrate that callose deposition triggered by CLas not only directly reduces solute transportation by the phloem by affecting sieve pore size, but also contributes to phloem cell death together with ROS, further reducing phloem function. This provides clear evidence that phloem cell death causes HLB-like symptoms.

## Effect of overexpression of *CsCalS7* on phloem callose deposition and citrus growth

We further aimed to mimic Clas-triggered callose deposition via overexpression of *CsCalS7a*, *CsCalS7b* and *CsCalS7c by* driving them with the 35S promoter. We were able to clone *CsCalS7b* and *CsCalS7c* but were unable to clone *CsCalS7a* despite multiple attempts. Transformants overexpressing *CsCalS7c* could be isolated as plantlets but could not further mature (Fig. S22). Phloem cell death was observed in *CsCalS7c-*overexpressing transformants (Fig. S22d), similar to that seen in *a cscals7a* mutant and *CalS7b-RNAi* plants (Fig. 4f; Fig. S17). However, overexpression of *CsCalS7b* driven by either the 35S promoter or the phloem-specific SUC2 promoter had no effect on plant growth, callose deposition, or sieve pores (Figs.S23 & S24), suggesting that CalS7 may be activated post-transcriptionally or post-translationally. Together our observations of the *cscals7a* mutant and *CalS7b-RNAi* and *CsCalS7c* overexpressing plants suggest that callose deposition around sieve pores induced by CLas contributes to HLB symptom development by reducing phloem transport of carbohydrates and causing phloem cell death in locations where callose deposition is severe.

## Trunk injection of HLB positive *C. sinensis* with CalS7 inhibitor 2-deoxy-d-glucose (DDG) reduces fruit drop and increases yield

We tested whether suppression of phloem callose deposition can alleviate the damaging effect of HLB. 2-deoxy-d-glucose (DDG) is known to inhibit callose synthesis ^35^. In a field trial, 7-year-old Valencia sweet orange trees were treated with DDG at different concentrations (0.5 mM, 1 mM, 2 mM) by trunk injection (100 ml per tree) in May 2024. Oxytetracycline (100 ml, 5500 ppm) was included as a positive control treatment ^3^. In another field trial with 14-year-old Valencia sweet orange trees, DDG at different concentrations (1 mM, 2 mM, 4 mM) were applied in July 2024. In both trials, oxytetracycline significantly reduced fruit drop, a common symptom of HLB, compared to non-treated controls as reported previously ^3^. In trials on the young trees, DDG at 2 mM, but not 0.5 and 1 mM, significantly reduced fruit drop compared to the non-treated control (Fig. S25a). In older trees in the second trial, DDG at 2 mM and 4 mM, but not 1 mM, significantly reduced fruit drop compared to non-treated controls (Fig. S25b). In both trials, both oxytetracycline and DDG increased the yield compared to the non-treated controls. Fruit yield was highest in trees receiving the highest concentrations of DDG (Fig. S25c, d). In one trial, DDG (2 mM) conferred similar yields as oxytetracycline (Fig. S25c), whereas its effect on yield was slightly lower than that of oxytetracycline in another trial (Fig. S25d). In addition, DDG (4mM) slightly increased (not statistically significant) CLas titers (Fig. S25e). The field studies provide evidence that phloem callose deposition plays an important role in HLB disease development.

## Knockout of tomato *Eds1* and *Pad4* but not *RbohB*, *Bik1* and *Sobir1* reduces disease symptoms, ROS production, callose deposition, and phloem cell death caused by *Ca*. Liberibacter

We next investigated whether immune signaling pathways are indeed responsible for the *Ca*. Liberibacter-triggered immune disease development. *Ca*. L. psyllaurous (Lpsy, synonym *Ca*. L. solanacearum) causes increased leaf yellowing, ROS production, cell death as indicated by trypan blue staining and ion leakage assays, phloem callose deposition and starch accumulation in tomato (Fig. 5), similar to that seen in HLB. We therefore utilized tomato-Lpsy as a model to further characterize the involvement of critical genes in the immune signaling pathways to symptom development given the quick turnaround time for mutant generation and disease testing. Specifically, we generated mutants for *Eds1*, *Pad4*, and *RbohB* (which is the *RbohD* homolog in *Arabidopsis* and a key enzyme responsible for PTI-induced apoplastic ROS production in plants ^36,37^), as well as for *BOTRYTIS-INDUCED KINASE1 (Bik1)* and *SUPPRESSOR OF BIR1-1 (Sobir1)*, two central components of PTI signaling (Figs. S26-S30) ^38–40^.

**Figure 5:**
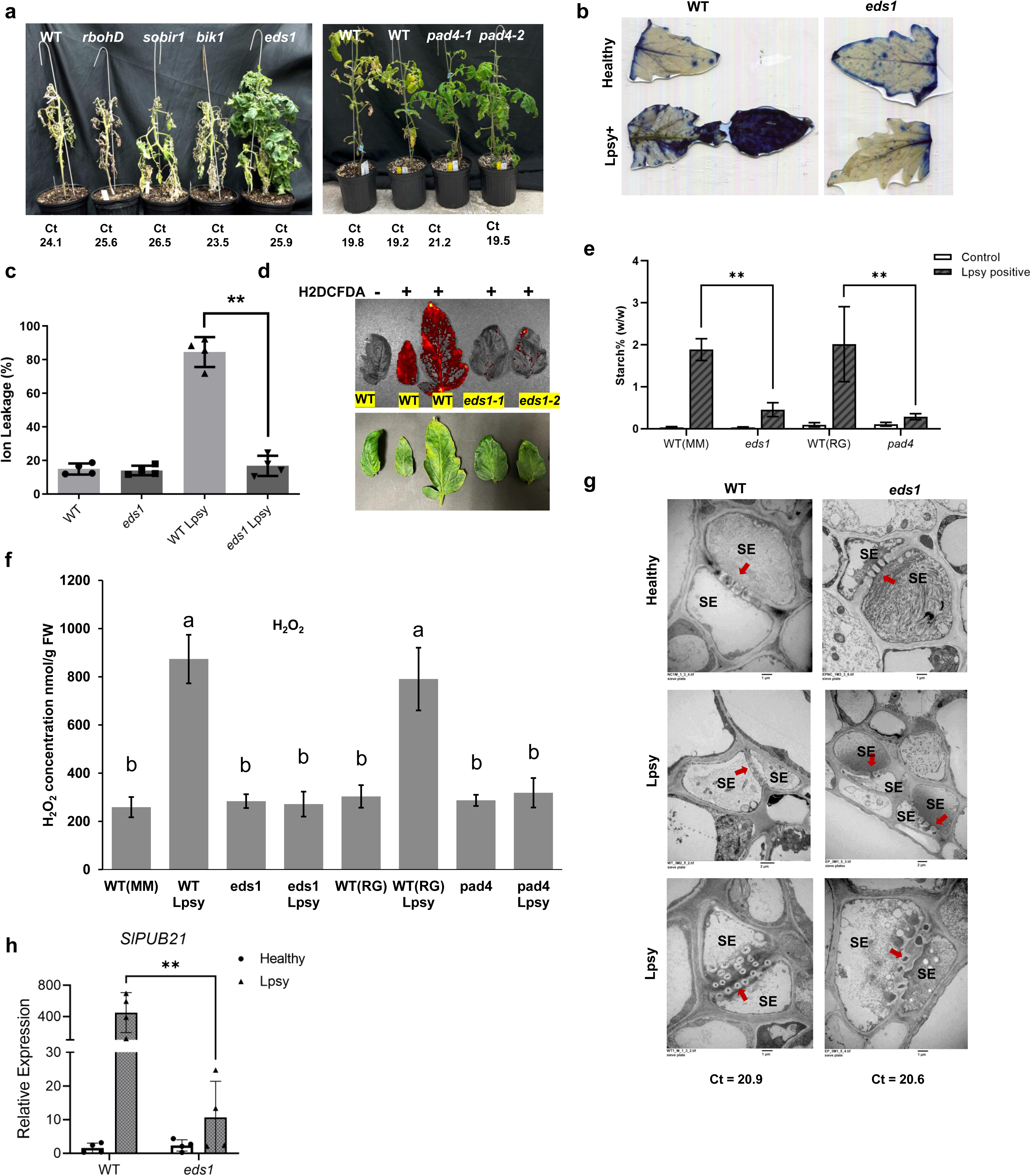
Mutation of *Eds1* and *Pad4* but not *RbohB*, *Bik1* and *Sobir1* reduces disease damages caused by *Ca.* L. psyllaurous (Lpsy), reduces ROS production, callose deposition, and phloem cell death. (**a**) Left: Tomato plants of the indicated genotypes were inoculated with *Candidatus* Liberibacter psyllaurous (Lpsy). Quantitative PCR (qPCR) was performed 21 days post-inoculation (DPI) to assess bacterial titers (Ct values). The photograph was taken at 60 DPI. Right: qPCR analysis was performed 6 weeks post-inoculation, and the corresponding image was captured at 7 weeks post-inoculation. (**b**) Trypan blue staining of tomato leaves from healthy and Lpsy-infected plants, highlighting cell death in infected tissues. (**c**) Cell death in tomato leaves of the indicated genotypes, measured by ion leakage assay. Ion leakage was calculated as the percentage of initial ion conductivity relative to total conductivity after boiling. (**d**) Reactive oxygen species (ROS) detection in H₂DCFDA-stained tomato leaves, visualized using a PerkinElmer IVIS Lumina S5 imaging system. Red fluorescence indicates ROS accumulation. (**e**) Quantification of leaf starch content in healthy (control) and Lpsy-infected tomato plants. Total starch levels were normalized to leaf fresh weight. (**f**) Quantification of H_2_O_2_ in healthy and Lpsy-infected tomato plants. (**g**) Transmission electron microscopy (TEM) analysis of callose deposition in the midveins of indicated tomato genotypes under healthy and Lpsy-infected conditions. Red arrows indicate sieve pores. SE, sieve element. (**h**) qRT-PCR analysis of *SlPUB21* relative expression levels in *eds1-*edited lines and wild-type tomato under healthy and Lpsy-infected conditions. The actin gene was used as an internal control. Scale bars represent mean ± SD, n=4. Asterisks indicate statistically significant differences by one-way ANOVA and Tukey’s test (*P* < 0.01).

The *eds1* and *pad4* tomato mutants remained relatively healthy after inoculation, whereas the wild type, *rbohB*, *bik1*, and *sobir1* showed typical disease symptoms and died approximately two months after inoculation (Fig. 5a). Interestingly, bacterial titers in the WT, *eds1* and *pad4* mutant plants were comparable. Since only *eds1* and *pad4* mutants exhibited improved disease resistance against Lpsy, we analyzed their effect on ROS production, phloem callose deposition, cell death, and starch accumulation compared to the WT. Mutation of *Eds1* or *Pad4* significantly reduced cell death (Fig. 5b, c), ROS production (Fig. 5d, f), starch accumulation (Fig. 5e), and phloem callose deposition (Fig. 5g). *PUB21* was reported to be a HLB susceptibility gene and its silencing increased disease resistance against CLas ^13^. Interestingly, mutation of *Eds1* drastically decreased the induction of *PUB21* by Lpsy (Fig. 5h).

## Discussion

In this study, we have shown that *Ca*. Liberibacter triggers the development of chronic immune disease through the Eds1 and Pad4-mediated induction of chloroplastic ROS production, and phloem callose deposition, which lead to phloem cell death. Our study provides strong evidence that phloem cell death triggered by CLas is responsible for HLB disease symptoms. While ROS have been shown to cause phloem cell death ^2^, this study suggests that phloem callose deposition can also contribute to phloem cell death when phloem callose deposition is very severe. In the HLB pathosystem, it seems that Clas-triggered chloroplastic ROS production and phloem callose deposition, which are typical immune responses, are the major factors for disease development even though we could not completely exclude disease effects from other putative virulence factors. Nevertheless, the immune disease model can explain HLB disease development and most of the known phenomena associated with HLB (Fig. S31). Importantly, trunk injection of HLB symptomatic trees with DDG, a callose deposition inhibitor, reduces fruit drop and increases fruit yield of HLB symptomatic trees. Application of antioxidants also reduces HLB disease symptoms ^2^. Such effects mirror that of treatment of human immune diseases using anti-inflammatory and immunosuppressant drugs ^1^. Although both ROS production and phloem callose deposition are regulated by Eds1, ROS do not directly influence phloem callose deposition. This contrasts with ROS-mediated callose deposition on the cell wall observed during PTI ^24^, but aligns with the Eds1-associated regulation of callose deposition in plasmodesmata ^41^. However, CalS7 is not involved in plasmodesmata callose deposition. How Eds1 triggers phloem callose deposition remains to be explored. The common phenomena associated with Liberibacter diseases such as blotchy mottle, starch accumulation, ROS production, and phloem callose deposition are reported with many other phloem-colonizing bacterial pathogens such as Phytoplasma and Spiroplasma ^42,43^. It is probable that the chronic immune responses triggered by these pathogens also contribute to their disease symptoms in addition to known virulence factors ^44^.

Mutation of key genes of the immune signaling pathway including *Eds1* and *Pad4* ^45,46^ but not *RbohB*, *Sobir1*, and *Bik1* abolishes the disease development caused by *Ca*. Liberibacter. It remains unknown how *Ca*. Liberibacter activates Eds1 and Pad4. Interestingly, multiple CLas effectors, flagellin and cold shock protein of CLas cause immune responses ^8–10,47,48^. Knockout of *Eds1* and *PAD4* improves disease resistance against Lpsy. Knockout of *Eds1* and *Pad4* also increases disease resistance against necrotrophic pathogens such as *Botrytis cinerea* ^49^. Together, the data reveals a unique feature of HLB, namely, that CLas is biotrophic, but causes disease damages though triggering chronic immune responses leading to phloem cell death, mimicking necrotrophic pathogens in pathogenicity. Similar Lpsy titers were observed in the wild type, *eds1* and *pad4* mutants, indicating that mutation of these two genes does not increase susceptibility to Lpsy. This is consistent with that DDG treatment has no significant effect on CLas growth. Together, the data suggests a slow and chronic nature of the immune responses caused by CLas, which are insufficient to kill CLas, but cause eventual phloem cell death. Nevertheless, CLas benefits from the leaf yellowing resulting from the immune responses, which is known to attract psyllid feeding and promote subsequent CLas transmission^50^. The recently identified HLB susceptibility gene *PUB21* ^13^ is required for R gene-mediated and elicitor infestin 1 (INF1)- triggered cell death ^51^. Silencing of *CMPG1*, a homolog of *PUB21*, confers resistance to *Phytophthora infestans* during the necrotrophic phase in *Nicotiana benthamiana* ^52^. Importantly, our study shows that *PUB21* is positively regulated by Eds1, which probably contributes to the increased disease tolerance of the *eds1* mutant against *Ca*. Liberibacter. Mutation of *Eds1* and *Pad4* likely improves disease resistance against necrotrophic pathogens, but reduces disease resistance against biotrophic pathogens ^49^. The benefit of improving disease tolerance against HLB using this approach likely overweighs the compromise given that commercial citrus production relies on scion varieties grafted onto rootstocks, and most commercial citrus rootstocks are resistant or tolerant to prevalent pathogens including HLB ^53^ and most scion diseases except HLB have effective control approaches. Importantly, non-transgenic *eds1* edited sweet orange in an ongoing study has received regulatory approval for commercialization by The Animal and Plant Health Inspection Service (APHIS) of the United States Department of Agriculture.

In summary, *Ca*. Liberibacter triggers chronic immune diseases through Eds1 and Pad4 mediated chloroplastic ROS production and phloem callose deposition. Genetic manipulation of *Eds1* and *Pad4* can convert *Ca*. Liberibacter into a benign endophyte, offering a promising strategy for precision breeding to enhance resistance/tolerance against HLB and potentially other *Ca*. Liberibacter associated diseases.

## Supporting information

Gene expression

## Data availability

The RNA-Seq data from *Cals7*, and whole genome resequencing data for Cals7 were deposited to NCBI under project ID PRJNA1258379.

## Acknowledgements

We thank Wang lab members for constructive suggestions and insightful discussions. This project was supported by funding from Florida Citrus Initiative Program, Citrus Research and Development Foundation, U.S. Department of Agriculture National Institute of Food and Agriculture grants 2022-70029-38471, 2025-70029-44030 and 2023-70029-41280, and Hatch project [FLA-CRC-005979] (N. Wang).

## Author Contributions

N.W. conceptualized and designed the experiments. WW, WM, XH, YF, SSP, TL, DSA, JL, JLD, YW, ZH, CR, NZ, MK, AH, and NW performed the experiments. J.X. performed bioinformatics. NW, WW, WM, XH, YF, SSP, JX, DSA, and JL wrote the manuscript with input from all co-authors. Specifically, among the co-first authors, WM led the work on CalS7b and identification phloem specific *CalS7a*, *b*, and *c*, WW led the work on CalS7a and CalS7c, XH and WW conducted work related to tomato-Lpsy, YF conducted work related to Fld, TL investigated chloroplastic ROS production, and JX conduct RNA-seq analysis.

## Competing interests

N. W. filed a PCT patent application based on the results reported in this paper. All other authors declare no competing financial interests.

**Correspondence and requests** for materials should be addressed to N. Wang.

## Materials and Methods

### Transmission electron microscopy (TEM) analyses

Sample sizes of 2-3 mm were taken from each tissue and placed in 3% glutaraldehyde in 0.2 M Sorenson’s phosphate buffer over night at 4°C. Then, the samples were postfixed in 2% osmium tetroxide prepared in 3% glutaraldehyde for 4 h at room temperature in a fume hood. The samples were dehydrated by sequential treatment with 10, 20, 30, 40, 50, 60, 70, 80, 90, and 100% (thrice) acetone for 10 min each. The samples were incubated sequentially in 50, 75, and 100% (twice) Spurr’s low-viscosity epoxy resin prepared in acetone for 8 h each. The blocks were trimmed with a surgical blade and then sectioned to 0.1 µm using a diamond knife under an ultramicrotome. The thin sections were collected on 200-mesh copper grids. The samples were stained with 2% aqueous uranyl acetate for 15 min, washed in water, and again stained with lead citrate, followed by water wash. The micrographs were prepared and analyzed using a Morgagni 268 (FEI Company, Hillsboro, OR, USA) transmission electron microscope equipped with an AMT digital camera (Advanced Microscopy Techniques Corp., Danvers, MA, USA).

### Quantification of CLas and Lpsy using quantitative real time PCR (qPCR)

Total genomic DNA from each plant sample was extracted as described previously ^2^. Briefly, samples were immediately frozen in liquid nitrogen and ground using the bead-beating method with Qiagen Tissue Lyser II (Qiagen, Hilden, Germany). Citrus genomic DNA was extracted using the CTAB (cetyltrimethylammonium bromide) method. DNA samples were then treated with the One Step PCR Inhibitor removal kit (Zymo). For qPCR assays, 100 ng of total genomic DNA was mixed with the Quantitec Probe PCR master mix (Qiagen) or SYBR Green Universal Master Mix (Applied Biosystems). Primers/probe CQULA04F-CQULAP10-CQULA04R were used for CLas ^3,4^ and Y DRAG Q-PCR-IGS-7F and Y DRAG Q-PCR-IGS-7R were used for Lpsy ^5^. All qPCR assays were performed using the QuantStudioTM 3-96-Well 0.1mL instrument (Thermofisher Scientific, Hillsboro, OR).

### H_2_O_2_ quantification

Hydrogen peroxide (H_2_O_2_) concentrations were quantified using a modified protocol from Ma et al. (2022)^6^. In brief, 0.5 g of plant samples were homogenized in 0.1% (w/v) trichloroacetic acid (TCA) and then centrifuged at 12,000 × g for 15 minutes at 4°C. The supernatant (0.3 mL) was mixed with 1.7 mL of 1.0 M potassium phosphate buffer (pH 7.0) and 1.0 mL of 1.0 M potassium iodide solution. After a 5-minute incubation, the absorbance of the resulting oxidation product was measured at 390 nm. H_2_O_2_ concentrations were calculated using a standard curve with known H_2_O_2_ concentrations and expressed as µmol/g fresh weight. For phloem-enriched bark tissue exudates, the H_2_O_2_ was estimated following the procedure as described previously ^7^. The H_2_O_2_ concentrations in phloem-enriched bark tissue exudates were expressed in mmol/L.

### Callose staining and observation

Leaf, stem and root samples were collected and flash-frozen in liquid nitrogen, then chopped into small slices. The samples were transferred to cold 4% paraformaldehyde fixative and vacuum infiltrated for 30 mins, and incubated overnight at 4°C. After one wash with 1X PBS for 15 mins, samples were dehydrated by sequential treatment with 10, 20, 30, 50, 70, 85, 95, and 100% (thrice) ethanol for 1 hour each. Samples were cleared in Ethanol:tert-Butanol (TBA) solutions (2:1, 1:1, 1:2, and 100% tert-Butanol) for 1 hour each. Liquid paraffin was infiltrated by sequentially incubating in 2:1, 1:1, and 1:2 TBA:paraffin solutions and 100% paraffin (thrice) for 6 hours each. Samples were embedded in liquid paraffin and solidified at room temperature. 8 µm sections were cut and heat-fixed on a heater and dewaxed to remove paraffin by placing the slides in two changes of Histoclear II for 15 minutes each. The sections were rehydrated in a rehydration ethanol series (100, 75, 50, 25% and water) for 5 minutes each. Slides were stained using the 0.05% aniline blue solution. One minute after staining, green fluorescence was observed under a fluorescent microscope in the UV channel. Blue fluorescence indicates xylem and fiber autofluorescence. Callose spots were quantified per slide area for all sample types.

### Starch assays

Root samples and vein tissues were harvested from cleaned leaves and roots and subsequently powdered using the TissueLyser II (Qiagen, Hilden, Germany) according to the manufacturer’s protocol. For starch quantification, 100 mg of the powdered samples were utilized. Starch estimation was carried out using the Total Starch Assay Kit (AA/AMG) from Megazyme (Bray, Ireland), following the manufacturer’s instructions. All experiments, unless otherwise noted, were repeated twice with consistent results.

### Histochemical detection of H_2_O_2_ and O_2_^·^− in citrus leaf tissues

Citrus leaves were submerged in a freshly prepared staining solution. For H_2_O_2_ detection, 1mg/ml solution of 3,3’-diaminobenzidine (DAB) was prepared by dissolving in low pH (3.8) water. For O_2_^·^− detection, 1 mg/ml of nitro blue tetrazolium (NBT) (dissolved in 25 mM HEPES-KOH) solution was prepared. The submerged leaves were stained for 24 hours under bench light conditions at room temperature with orbital shaking. After treatments, the samples were briefly rinsed with distilled water twice and decolorized with ethanol (95%). Samples were stored in ethanol and photographed with a digital camera by using an Olympus BX61 epifluorescence microscope (Olympus Corporation, Center Valley, PA, USA). H_2_O_2_ and O_2_^·^− were visualized as reddish-brown and dark-blue coloration spots localized within cell organelles, respectively.

### Total ROS and singlet oxygen (^1^O_2_) detection by fluorescence microscopy

Total intracellular ROS was detected by using 10 μM 2′,7′-dichlorodihydrofluorescein diacetate (H_2_DCFDA) dye (Cat. #D399, Thermo-Fisher Scientific). This dye is non-fluorescent in its reduced form but converts into fluorescent form inside cells when oxidized by H_2_O_2_ and O_2_^·^− and various free radical products downstream from H_2_O_2._ Mature leaves were used for cell isolation. Cells were isolated after enzymatic digestion (∼16 hours) with 1% cellulase RS and 1% macerozyme R-10 (PhytoTech labs, Lenexa, KS, USA). The isolated cells were treated with 10µm of H_2_DCFDA (stock was prepared in DMSO and the working solution was prepared in 1XPBS) then incubated at 37^0^C for 15 minutes in darkness and rinsed briefly with 1xPBS then visualized by fluorescence microscopy (BX61 epifluorescence microscope, Ex; 488nm, Em: 525 nm). To eliminate the damages of low osmolarity on protoplasts, the procedure was done within a short time.

For singlet oxygen detection, Singlet oxygen sensor green reagent (SOSG) was used (Cat# S36002, Thermo Fisher, Willow Creek Road, Eugene, OR, USA). 10 µm of SOSG working solution was prepared in 1X PBS by dissolving the stock (prepared in methanol). The isolated citrus cells from HLB-infected (mildly symptomatic) and healthy leaves were incubated in the dark for 2 hours at room temperature and then visualized by fluorescence microscopy (BX61 epifluorescence microscope, Ex: ∼504 nm, Em: ∼525 nm). The presence of singlet oxygen emits a green fluorescence in cells. All results were repeated at least twice. It is noteworthy that the procedure caused cell damages, but obvious differences were observed between HLB positive and healthy samples.

### RNA-Seq sequencing and data analysis

Total RNA was extracted from mature leaves using the RNeasy plant kit (Qiagen, Valencia, CA), followed by treatment with RQ1 RNase-Free DNase (Cat. #M6101, Promega, Madison, WI).

RNA-Seq Libraries were constructed with the NEBNext Ultra II RNA Library Prep Kit from Illumina (San Diego, CA, U.S.A.). Samples were sequenced to generate 150-bp paired-end reads using the Illumina NovaSeq 6000 platform (Illumina). Data were filtered to remove low-quality reads and adapters by Novogene. High quality reads were then aligned to the *Citrus sinensis* v3 genome ^8,9^ with HISAT2 ^10^ and SAMtools ^11^ and raw counts were quantified using HTSeq-count ^12^ to generate the gene expression read count profile. Based on the normalized gene expression data using the DESeqVS and log2 scale methods ^13^, samples were clustered based on Euclidean distance. Differentially expressed gene (DEG) analysis was performed using DESeq2 packages in R ^13^. Genes were considered significantly expressed with an adjusted *P* < 0.05 (FDR method) ^14^. Gene Ontology (GO) term and KEGG pathways enrichment of DEGs was analyzed using g:Profiler web tool ^15^. Heatmap plots of gene expression data were drawn using the ComplexHeatmap package in the R program ^16^.

### GO enrichment analyses of previously published transcriptome data between HLB and healthy citrus samples

To generate the comprehensive expression pattern of citrus plants in response to CLas infection, we collected 8 microarray and 21 RNA-seq data sets across different genotypes and tissues from NCBI SRA and GEO databases (Data S1). For RNA-seq data sets, the high-quality short reads were aligned to the *Citrus sinensis* genome ^8,9^ with HISAT2 ^10^ and SAMtools ^11^. The read counts were quantified using HTSeq-count ^12^ to generate the gene expression profile. The differentially expressed genes (DEGs) were determined using Limma version 3.46.0 ^17^ and DESeq2 version 1.30.1 ^13^ packages in R version 4.2 for microarray and RNA-seq data, respectively (adjusted *p* value <0.05). The up- or down-regulated genes were determined based on the strategy that DGEs showed the same expressed trends in at least 75% of the comparisons. The Gene Ontology (GO) term enrichment analysis of up- or down-regulated genes was performed using g:Profiler web tool ^15^.

### Constructs

Unless otherwise specified, all gene cloning in this study was achieved using the seamless gene assembly technology (In-Fusion Snap Assembly Master Mix, Takara Bio, San Jose, CA, USA).

To generate the expression construct erGFP-1380N-CaMV35S-TP-Fld for citrus transformation, we synthesized the DNA fragment encoding TP-Fld from IDT (Integrated DNA Technologies, Coralville, IA). The fragment introduced into the erGFP-pCAMBIA-1380N-CaMV35S binary vector at the *Xba*I and *Sal*I sites and fused with the C-terminal 3×HA tag. To generate the transient expression construct TP-Fld-mVenus for subcellular localization in *Nicotiana benthamiana*, the pCAMBIA1380 vector was digested with *Sbf*I and *Eco*RI and ligated with a CaMV35S-mVenus insert. The *TP-Fld* coding sequence was amplified and cloned into the *Xba*I and *Bam*HI sites of the pCAMBIA1380-CaMV35S-mVenus binary vector.

To generate plasmids used for RNAi silencing of *CsCalS7b* in *C. sinensis* cv. Hamlin, 531 bp sequence spanning part of the 5’UTR and CDS were amplified from Hamlin cDNA. Primers Cals7a-sens-F/R were used to amplify the sense strand (S), Cals7a-AS-F/R were used to amplify antisense strand (AS), and Intron-F/R were used to amplify a 199 bp segment of the first intron of potato GA20 oxidase as the loop (Table S1). The S, loop, and AS fragments, along with the binary vector erGFP-1380N plasmid ^4^ digested by *Xba*I and *Stu*I, were ligated using T4 DNA ligase, generating the pCals7b-RNAi plasmid.

To generate 35S_pro_:RUBY reporter for transgenic plant screening, the RUBY system was introduced into the erGFP-1380N plasmid. Specifically, three key genes, *CYP76AD1*, *DODA*, and *glucosyl transferase* were fused with 2A peptide and inserted downstream of the 35S promoter at the *Xba*I and *Sal*I sites of the erGFP-1380N plasmid, generating the 1380-erGFP-35s_pro_:RUBY construct.

To generate *CsCalS7_pro_*:erGFP-35S_pro_:RUBY plasmids, the promoters and erGFP were amplified and inserted into the erGFP-1380N plasmid digested by *Sbf*I and *Sac*I to replace CsVMVpro:erGFP. *CsCalS7a*, *CsCalS7b*, and *CsCalS7c* promoters were amplified from Hamlin genomic DNA. The promoters were predicted using Softberry (Solovyev VV, Shahmuradov IA, Salamov AA. 2010). Specifically, 1988 bp, 1166 bp, and 1578 bp fragments upstream of the translation start codon (ATG) of *CsCals7a*, *CsCalS7b,* and *CsCalS7c* were amplified as the promoters, respectively.

The plasmids pXH1-CmYLCV and pCXH2-CmYLCV were used to generate the constructs for gene editing of *CsCals7a* and *CsCalS7c* in citrus ^18^. N-terminal specific gRNAs were selected using CRISPR-P (http://crispr.hzau.edu.cn/cgi-bin/CRISPR2/CRISPR#) and assembled into the vector by Golden gate cloning, as described previously ^18^. The cassettes were cut from pXH1-CmYLCV-*CsCals7a* and pXH1-CmYLCV-*CsCals7c* by *Fsp*I and *Xba*I and inserted into the *Zra*I/*Xba*I site of the binary vector PCXH2 to make the final constructs PCXH2-CmYLCV-*CsCals7a* and PCXH2-CmYLCV-*CsCals7c*.

To generate the *CsCalS7b* overexpression construct for citrus transformation, the coding region of *CsCalS7b* was amplified from sweet orange leaf cDNA. Due to the high complexity and the secondary structures of the sequence, we divided the full length (5769 bp) into 6 fragments, which include 4 fragments amplified using gene-specific primers and 2 fragments synthesized by IDT (Integrated DNA Technologies, Coralville, IA) (Table S1). All fragments were assembled by NEBulider HiFi DNA Assembly Cloning Kit (New England Biolab) with the binary vector erGFP-1380N-AtSUC2 or erGFP-1380N-CaMV35S ^4^ and fused with C-terminal 3xHA protein tag. For the 35S-*CsCalS7c* overexpression construct, the full length of 5712 bp was divided into two fragments for amplification. Two fragments were assembled by NEBulider HiFi DNA Assembly Cloning Kit (New England Biolab) into the binary erGFP-1380N-CaMV35S, also fused with C-terminal 3xHA protein tag. All constructs were confirmed by full-length Sanger sequencing. An empty vector (EV) was used as a negative control.

### Phloem proportion measurement

For calculating the phloem proportion, the midvein diameter was divided by the phloem width (the distance from the inner phloem fiber to the outer xylem). Images generated during callose staining observation were used to measure the phloem. Seven-11 leaves were collected from different lines, and 5 sections from each leaf were measured.

### Reverse transcription quantitative PCR (RT-qPCR) and semi-quantitative PCR

RT-qPCR was performed as described previously ^6^. Briefly, total RNA was extracted using the RNeasy plant kit (Qiagen, Valencia, CA), followed by treatment with RQ1 RNase-Free DNase (Cat. #M6101, Promega, Madison, WI). RNA concentration and quality were measured by a Nanodrop One Microvolume UV-Vis Spectrophotometer (Thermo-Fisher Scientific, Waltham, MA). 1 μg RNA was added for first-strand cDNA synthesis using ImProm-II Reverse Transcription System (Promega) and diluted 5 times for qRT-PCR to detect expression of callose synthase coding genes. 12 μL of qPCR reaction consisted of 6 µL of 2×KiCqStart SYBR Green qPCR ReadyMix (Sigma-Aldrich), 2.4 µL of each primer (1.25 µM), and 1.2 µL of diluted cDNA template. The PCR cycling consisted of an initial activation step at 95°C for 3 min, followed by 40 cycles of 95°C for 15 s and 60°C for 40 s. Relative gene expression was calculated using the 2−ΔΔCT method ^19^. All cDNA samples were run in triplicate. The *glyceraldehyde-3-phosphate dehydrogenase* (*GADPH*) gene was included as an endogenous control. For semi-quantitative PCR, gene-specific primers were used to amplify the *CsCalS7b* and *CsCalS7c* from cDNA templates. The *GAPDH* gene was used as an internal control. The primer sequences for specific genes and endogenous control genes are listed in Table S1.

### Epicotyl transformation and transgenic plants validation

*C. sinensis* cv. Hamlin was used for *Agrobacterium*-mediated epicotyl transformation. In brief, binary vectors were transferred into *A. tumefaciens* strain EHA105. Etiolated epicotyl segments of *C. sinensis* cv. Hamlin were used for *A. tumefaciens*-mediated transformation. After co-culturing with EHA105 harboring the binary vectors for 3 days, epicotyls were moved to screening agar plates. Positive shoots selected by kanamycin-resistance and erGFP fluorescence were micro-grafted onto 1-month-old rootstock (Carrizo (*Citrus sinensis* × *Poncirus trifoliata*)) seedlings and cultured in liquid MS medium. After one month, the surviving plants were transferred into commercial potting soil and grew in the growth chamber.

Plant Direct PCR Master Mix (Thermo Fisher) was used to amplify the integrated DNA fragment. Specifically, a 0.35 mm punch disc was collected from transgenic plant leaves and ground in 50 μL dilution buffer provided in the kit. After centrifuging, 2 μL of supernatant was added as template for PCR.

### DNA extraction and genotyping

Genomic DNA was extracted from genome edited or wild type citrus leaf using the CTAB method as described previously ^20^. Click or tap here to enter text. For genotyping, a ∼500 bp fragment spanning two gRNAs was amplified and cloned into the cloning vector (Zero Blunt™ TOPO™ PCR Cloning Kit, Invitrogen). 10-15 colonies were randomly selected and sent out for Sanger sequencing using M13 primers.

### Root generation mediated by *Rhizobium rhizogenes*

Roots were generated from citrus branches or epicotyl mediated by *Rhizobium rhizogenes* ^21^. Briefly, excised branches from transgenic plants or cut epicotyls from wild-type Carrizo seedlings were soaked in *Rhizobium rhizogenes* (OD_600_=1.0), followed by vacuum infiltration for 30 mins. Then, explants were transferred to the inert vermiculite matrix in pots. Pots were put in a tray with lid to maintain high humidity. Roots were generated around 1 month after treatment.

### Confocal microscopy

Transgenic Carrizo roots and shoots expressing RUBY were sectioned in both cross and longitudinal directions using razor blades. Specimens were soaked in ClearSee solution (Fujifilm) until they became transparent. Samples were rinsed with PBS buffer three times and then transferred to 0.05% aniline blue solution for 10 mins. GFP fluorescence (Ex488nm/Em510nm) and callose (Ex405nm/Em480nm) stained by aniline blue were detected using a confocal laser scanning microscope.

### Western Blotting

Western blotting analysis using anti-HA antisera was performed to detect Fld expression in ‘Hamlin’ mutant plants. Briefly, leaf tissue was ground to a fine powder in liquid nitrogen and added CelLytic™ P Cell Lysis Reagent (Sigma-Aldrich, MO, U.S.A. #C2360) for lysis at 4°C, and centrifuged to separate leaf debris from aqueous proteins. The supernatant was collected and mixed with 4× Laemmli buffer (Bio-Rad, #1610747), boiled for 10 min, then run on a 12% polyacrylamide gel (Bio-Rad, #1610175) via sodium dodecyl sulfate polyacrylamide gel electrophoresis. The proteins were transferred from the gel to nitrocellulose membrane (Bio-Rad Hercules, #1620112) by Trans-Blot Turbo System (Bio-Rad, model Trans-Blot^R^ Turbo™ System). Membranes were blocked in 5% nonfat skim milk solution for 2 hours, incubated with anti-HA antibodies at a dilution of 1:1,000 (Sigma-Aldrich,# 11867423001) overnight, followed by a goat anti-rat HRP secondary antibody (1:1,000, Abcam, ab97057) for 2 hours, and washed 3 time with TBS-T(Tris Buffered Saline with 0.1% Tween 20 (sigma, #P1379)). Detection was carried out using the SuperSignal West Pico Chemiluminescent Substrate kit (Fisher Scientific, #PI34578), and signals were captured on an Azure Imaging System (Azure Biosystems, model: Azure 300). Ponceau staining (Aqua Solutions, SPE842A) was used to indicate protein loading amount.

### MAPK phosphorylation assay

**The** MAPK phosphorylation assay was conducted following the previously reported method ^22^. Healthy and CLas-positive leaves were collected. The samples were flash-frozen in liquid nitrogen and ground using a TissueLyser II (Qiagen, Germany). To each sample, 200 μL of extraction buffer was added, which contained 50 mM HEPES (pH 7.5), 50 mM NaCl, 10 mM EDTA, 0.2% v/v Triton X-100, Pierce Protease Inhibitor Mini Tablets (EDTA-free, Cat. #A32955, Thermo), and Pierce Phosphatase Inhibitor Mini Tablets (Cat. #A32957, Thermo). The samples were then centrifuged at 15,000 × g for 10 minutes to pellet the cell debris.

Protein concentrations were determined using the Bio-Rad Protein Assay Dye Reagent (Cat. #5000006, Bio-Rad). MAPKs were detected by immunoblotting with an anti-p44/42 MAPK antibody (1:2,000, Cell Signaling Technology #4370L), followed by a goat anti-rabbit HRP secondary antibody (1:3,000, Bio-Rad #170-5046). Membranes were developed using the SuperSignal West Pico Chemiluminescent Substrate kit (Fisher #PI34578) and visualized on an Azure Imaging System (Azure Biosystems, USA).

### Ion leakage

Four leaf disks (0.35 mm in diameter) per leaf were collected and submerged in 3 mL of sterilized water with gentle shaking overnight. The sample ion leakage was measured using a CON 700 conductivity/°C/°F bench meter (Cat. #WD-35411-00, OAKTON Instruments, Vernon Hills, IL, USA). To determine the total ion leakage, the samples were boiled for 10 minutes, then cooled to room temperature and measured again using the same meter. The final ion leakage was calculated as the percentage of sample ion leakage divided by total ion leakage. The experiments were repeated twice with similar results. For ion leakage analysis of roots, samples were collected and washed with water to remove soil. Four fibrous roots of ∼2cm long were cut using scissors for each replicate. Immediately after collection, samples were immersed in a tube containing 2 mL of DI water for 16 hours by orbital shaking at 125 rpm. The ion leakage test was measured by using a CON 700 conductivity/°C/°F bench meter. Initial conductivity was measured in non-boiled samples, and total conductivity of the samples was measured after boiling of samples in water for 15 minutes. The total cell death percentage of each sample was calculated by using the initial and final conductivity values as (initial/final) x 100.

### Agrobacterium-infiltration assay in N. benthamiana

*A. tumefaciens* strain EHA105 carrying the binary vector was grown overnight in LB medium supplemented with 50 µg/mL rifampicin and 50 µg/mL kanamycin, then pelleted and resuspended in induction medium (10 mM MgCl₂, 10 mM MES pH 5.6, 200 µM Acetosyringone). After a 4 h incubation at room temperature with shaking, cultures were adjusted to an OD₆₀₀ of 0.4. For each construct, three leaves of young *N. benthamiana* plants were infiltrated (in triplicate) with the diluted bacterial suspension.

### Confocal localization of TP-Fld-mVenus in *N. benthamiana*

Three days after infiltrating *N. benthamiana* with EHA105 carrying the binary vectors, the infiltrated leaf tissue was collected and green fluorescence of the mVenus and chloroplast autofluorescence were visualized by confocal laser scanning microscopy (CLSM) (Leica TCS-SP5, Mannheim, Germany) with excitation/emission at 488 nm /510 nm.

### Field trials

Two field trials were conducted at commercial citrus groves in Avon Park and Polk City, Florida respectively. Both groves are 10 acres or more with approximately 220 trees/acre. The Avon Park grove trees were 7-year-old Valencia sweet orange (*C. sinensis* L. Osbeck) trees on US-802 (*C. grandis* ‘Siamese’ × *Poncirus trifoliata* ‘Gotha Road) rootstock. The Polk City grove trees were 14-year-old Valencia sweet orange (*C. sinensis* L. Osbeck) trees on X639 rootstock (*P. trifoliata* x *C. reticulata* cv. Cleopatra mandarin). Trunk injections were performed on sunny days during the first week of May and July 2024 for the Avon Park and Polk City grove, respectively. Injections were completed using FLexInject® Injectors (TJ BioTech LLC, Buffalo, SD) following the manufacturer’s instructions. Briefly, a hole on the trunk was drilled 15 cm below the first branch to a depth of 2 to 3 cm using a 7.14-mm drill bit. An injector was directly inserted into the drilled hole and was removed once all treatment materials were taken up by the tree. FLexInject is a pressure fluidized latex injection device that releases liquid at 200-300 kPa. One injector was used for each injection, with 50 or 100 ml of compound formulation dissolved in water. The Avon Park trial consisted of six treatments: (i) untreated control (injected with water); (ii) OTC: one injection of 100 ml oxytetracycline (5500 ppm); (iii) OTC2: two injections of 50 ml oxytetracycline (5500 ppm) at opposite sides of the tree; (iv) T500: 2-Deoxy-D-glucose at 0.5 mM in 100 ml water; (v) T1000: 2-Deoxy-D-glucose at 1.0 mM in 100 ml water; and (vi) T2000: 2-Deoxy-D-glucose at 2.0 mM. T4000 in 100 ml water. The OTC formulation was ReMedium TI® Injectors (TJ BioTech LLC, Buffalo, SD) and the amount of OTC used was based on the manufacturer’s recommendations. A whole row of 95 trees was injected for each treatment and a total of six rows were used for this study. The Polk City trial included five treatments: (i) untreated control (injected with water); (ii) OTC: one injection of 100 ml oxytetracycline (5500 ppm); (iii) T1000: 2-Deoxy-D-glucose at 1 mM in 100 ml water; (iv) T2000: 2-Deoxy-D-glucose at 2.0 mM in 100 ml water; (v) T4000: 2-Deoxy-D-glucose at 4.0 mM in 100 ml water. A full row of 80 trees were injected per treatment, with five rows used in total for the study. For both trials, pre-harvest fruit drop was examined in February 2025 and fruit yield was quantified in April 2025 during harvest.

### Generation of tomato mutants

The tomato mutants were generated using CRISPR/Cas12a-based method or CRISPR/Cas9-based method described previously ^23,24^. *SlRbohD* mutants were described previously ^24^. The crRNA for *SlEDS1* (Solyc06g071280) is GAGAGAAAGAGATCAACACCACT. The crRNA for *SlBIK1a* (Solyc10g084770) is ATGAACTTCGATCTGCAACCAGA. The crRNA for *SlBIK1b* (Solyc04g011520) is AGATTCGACGATTCAAGTATTTC. The crRNA for *SlSobir1a* (Solyc06g071810) is CCAATAGCAGAAGACAAAGTTCC. The crRNA for *SlSobir1b* (Solyc03g111800) is CTGGCAGAATACCTCAATCCTTG. The gRNA for *SlPAD4* (Solyc02g032850) is GGCGTCAACGCCGTTGCTGG.

### Infection of tomato plants with *Ca.* L. psyllaurous (Lpsy)

To inoculate tomato plants with Lpsy, we utilized the tomato psyllid *Bactericera cockerelli* western biotype ^25^. This genetic line of *B. cockerelli* was infected with Lpsy haplotype B. For each mutant, five plants were inoculated with one plant as a biological replicate. For inoculation, 15 Lpsy biotype B infected-male adult psyllids were kept within a cage around each plant biological replicate. Male psyllids were used to ensure no eggs are laid during inoculation trials because eggs can result in continued inoculation via nymphs and lead to unequal inoculations across replicates. Tomato plants used for inoculation were approximately six-week-old and psyllids were allowed to feed for one week inside of the cage. After one week of feeding psyllids were subsequently removed to ensure all plant biological replicates were inoculated for the same duration of time. For each batch, five wild type tomato plants were inoculated with Lpsy as controls. To confirm that inoculations were successful the top-tier leaves were screened for Lpsy infection using qPCR one week after psyllids were removed from plants with Lpsy specific primers following the plant DNA extraction and qPCR protocols detailed in ^5^.

### ROS imaging

Tomato branches were placed in the solution containing 10 μM 2′,7′-dichlorodihydrofluorescein diacetate (H_2_DCFDA) (Cat. #D399, Thermo-Fisher Scientific) for 24 hours. Leaves at the upper parts, without contact with the solution, were detached for imaging in a PerkinElmer IVIS Lumina S5 imager with GFP channel.

### Trypan-blue staining

Trypan Blue staining was performed with modifications based on the method described by Fernández-Bautista et al. ^26^. Briefly, freshly collected leaves were placed in Petri dishes and fully immersed in 10 mL of 0.4% Trypan Blue solution (Cat. #15250061, Thermo Fisher). Samples were incubated with gentle shaking overnight. Following staining, leaves were decolorized in 98–100% ethanol until the green pigmentation was completely removed. Then, the samples were covered in 60% glycerol for observation.

### DNA Library construction, sequencing, and data analysis for confirmation of genome-edited plants

The DNA library was prepared following the manufacturer’s protocol of short read DNA sequencing from Illumina ^27^. After quality control, quantification, and normalization of the DNA libraries, 150-bp paired-end reads were generated using the Illumina NovaSeq 6000 platform according to the manufacturer’s instructions at Novogene, China. The raw paired-end reads were filtered to remove the low-quality reads using fastp program version 0.22.0 ^28^. To assess the target site mutations of mutated plants, the high quality paired-end short genomic reads were mapped to the sweet orange (*C. sinensis*) ^8,9^ reference genome using Bowtie2 software version 2.2.6 ^29^. The mutations (single nucleotide polymorphisms, deletions and insertions) for the mutated plant genomes were generated using the SAMtools package version 1.2 ^11^ and deepvariant program version 1.4.0 ^30^. The generated mutations were filtered by quality and sequence depth (mapping quality > 10 and mapping depth > 10). The mutations of target sites were visualized using IGV software version 2.15.4 ^31^. Based on the mapping results, mutations of off-target sites were detected using the SAMtools package version 1.2 and deepvariant program version 1.4.0. Potential off-target regions were also amplified and validated by Sanger sequencing.

---

**Supplementary Table 1.**
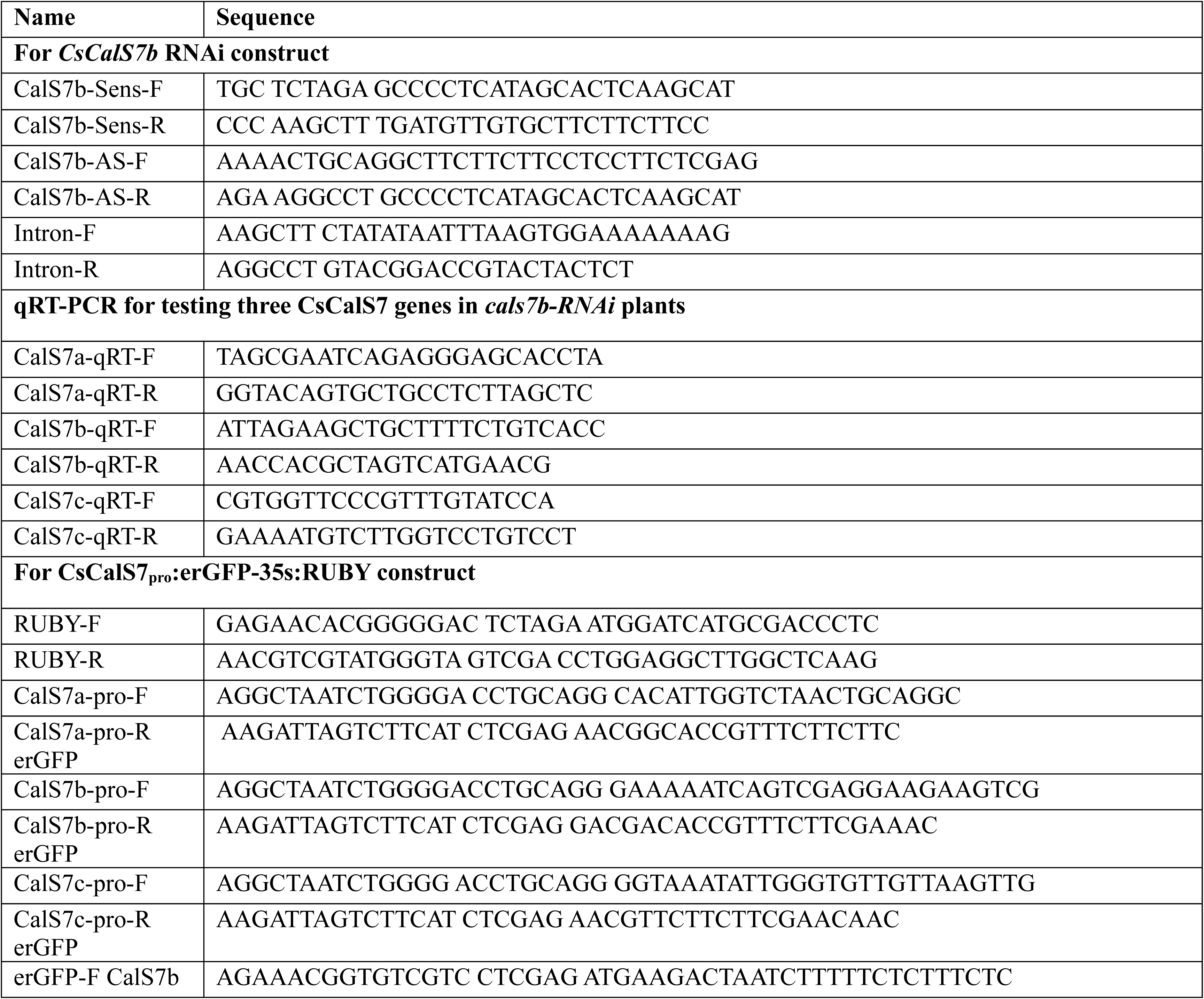
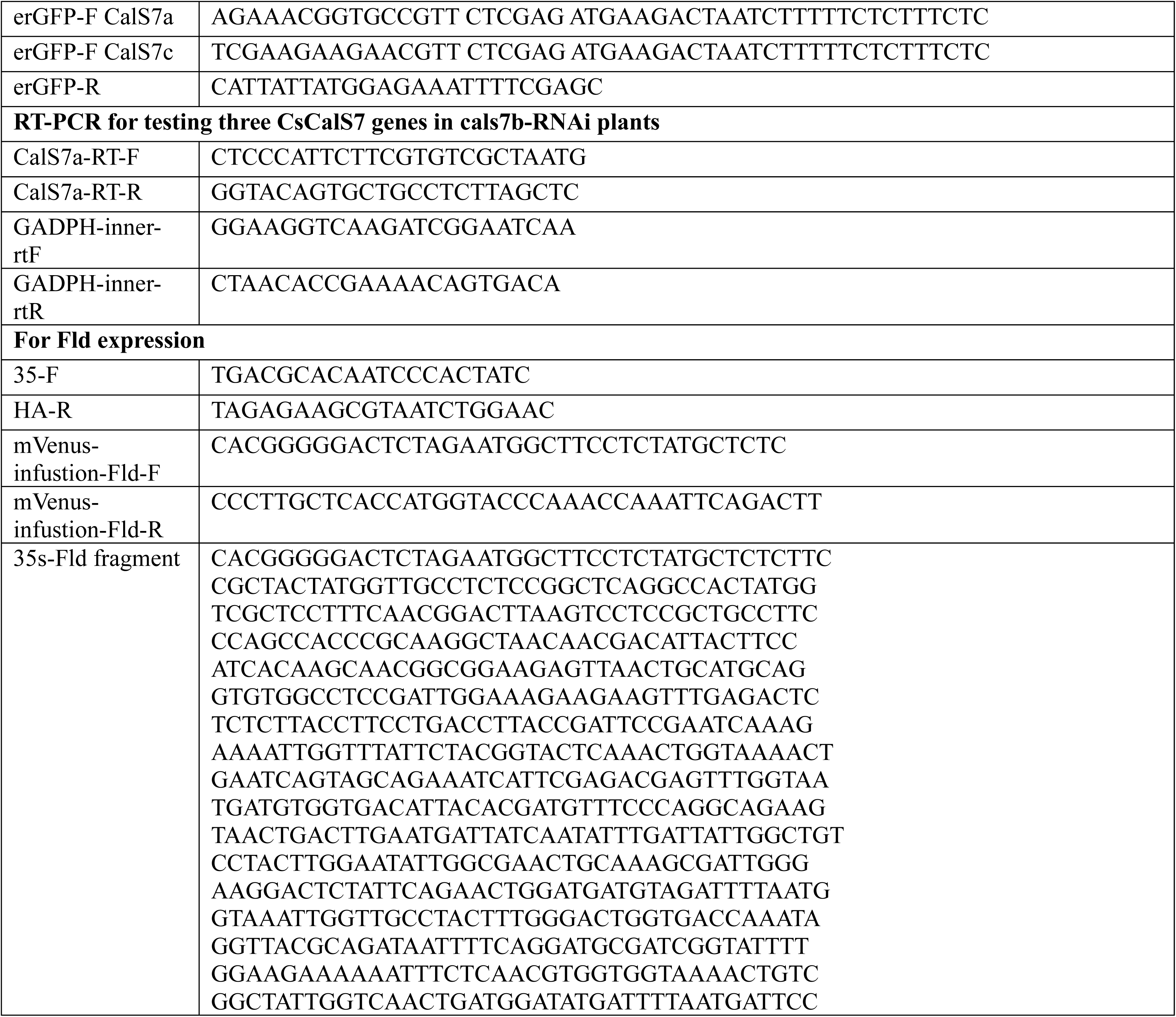
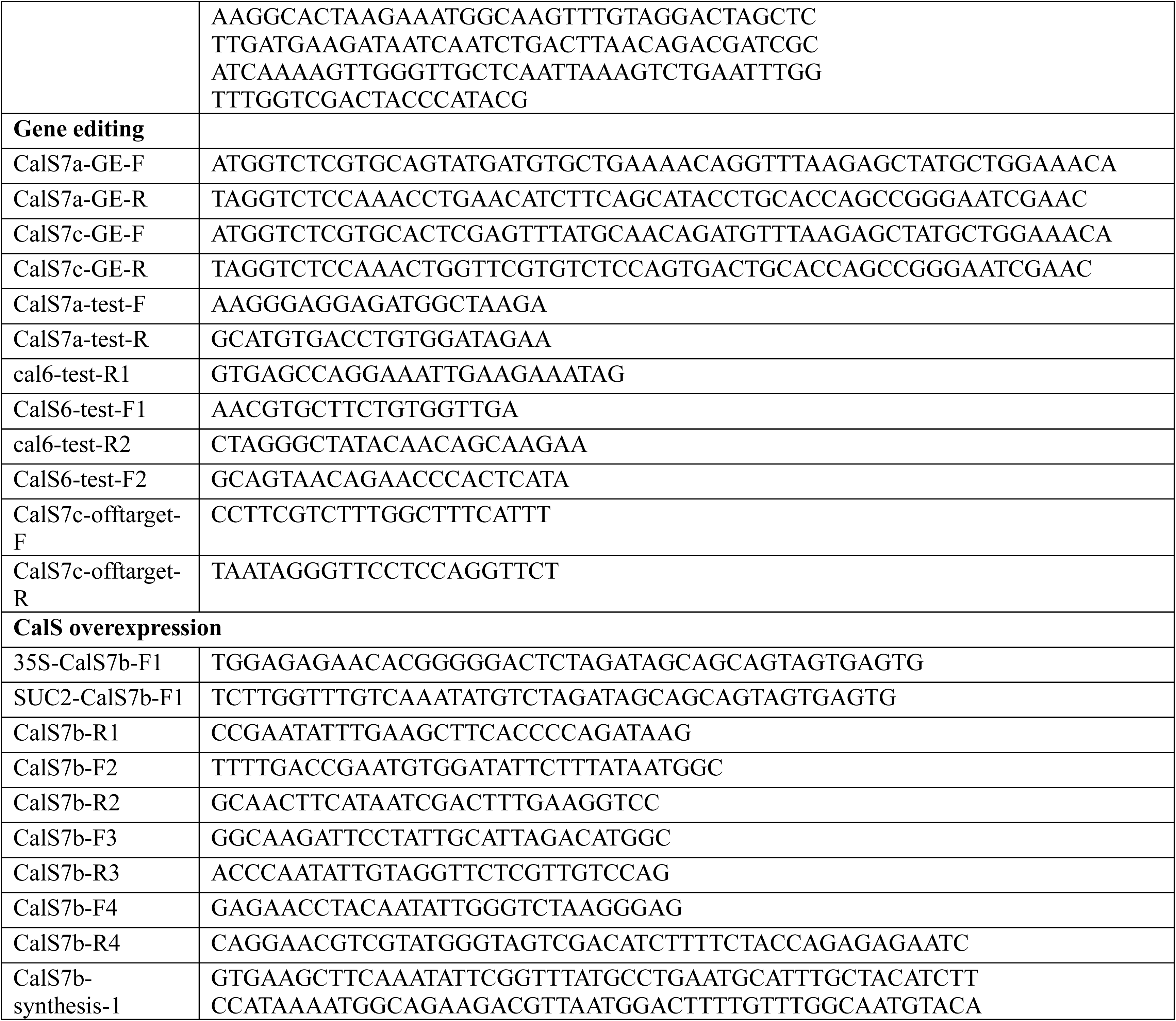
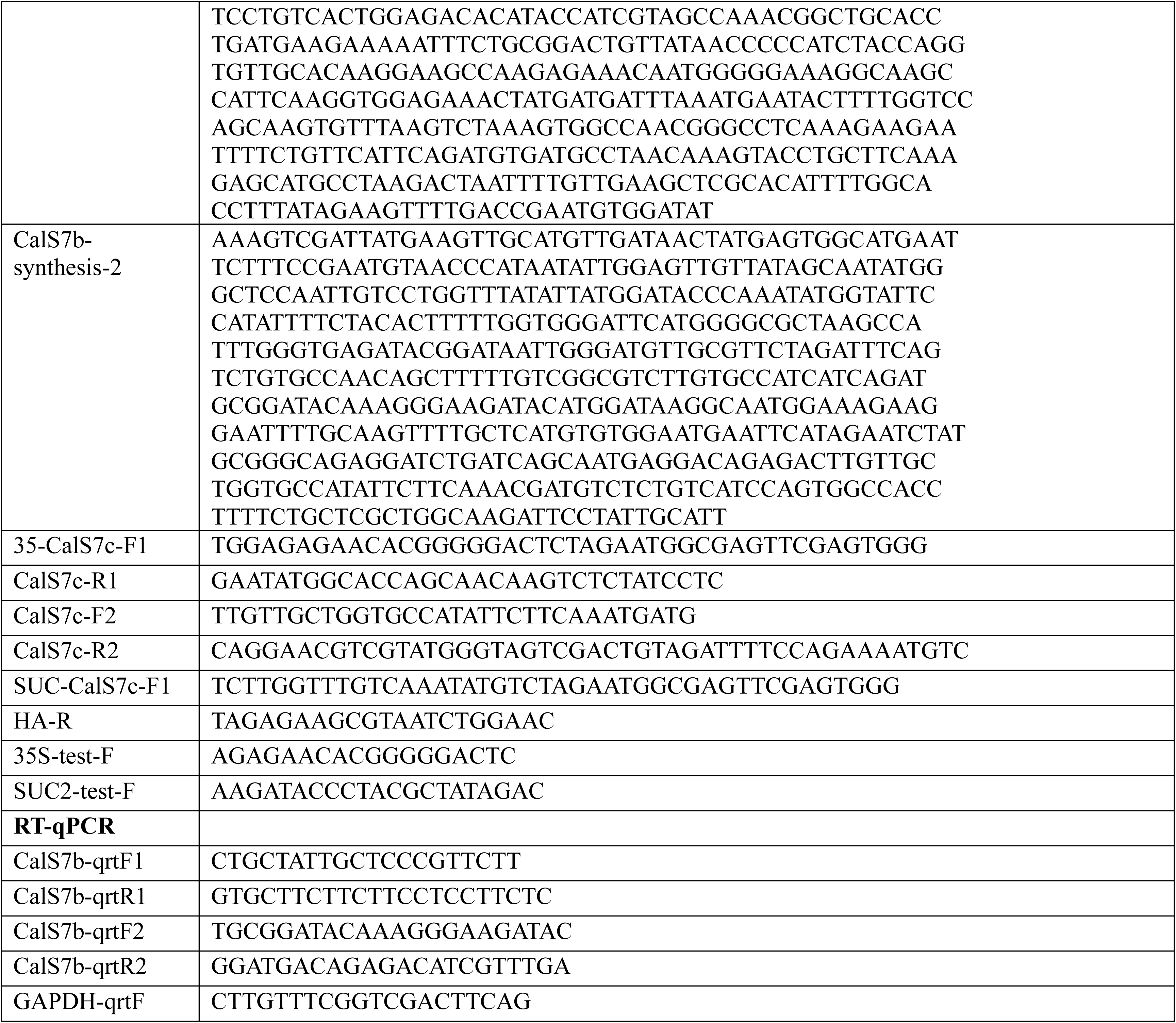
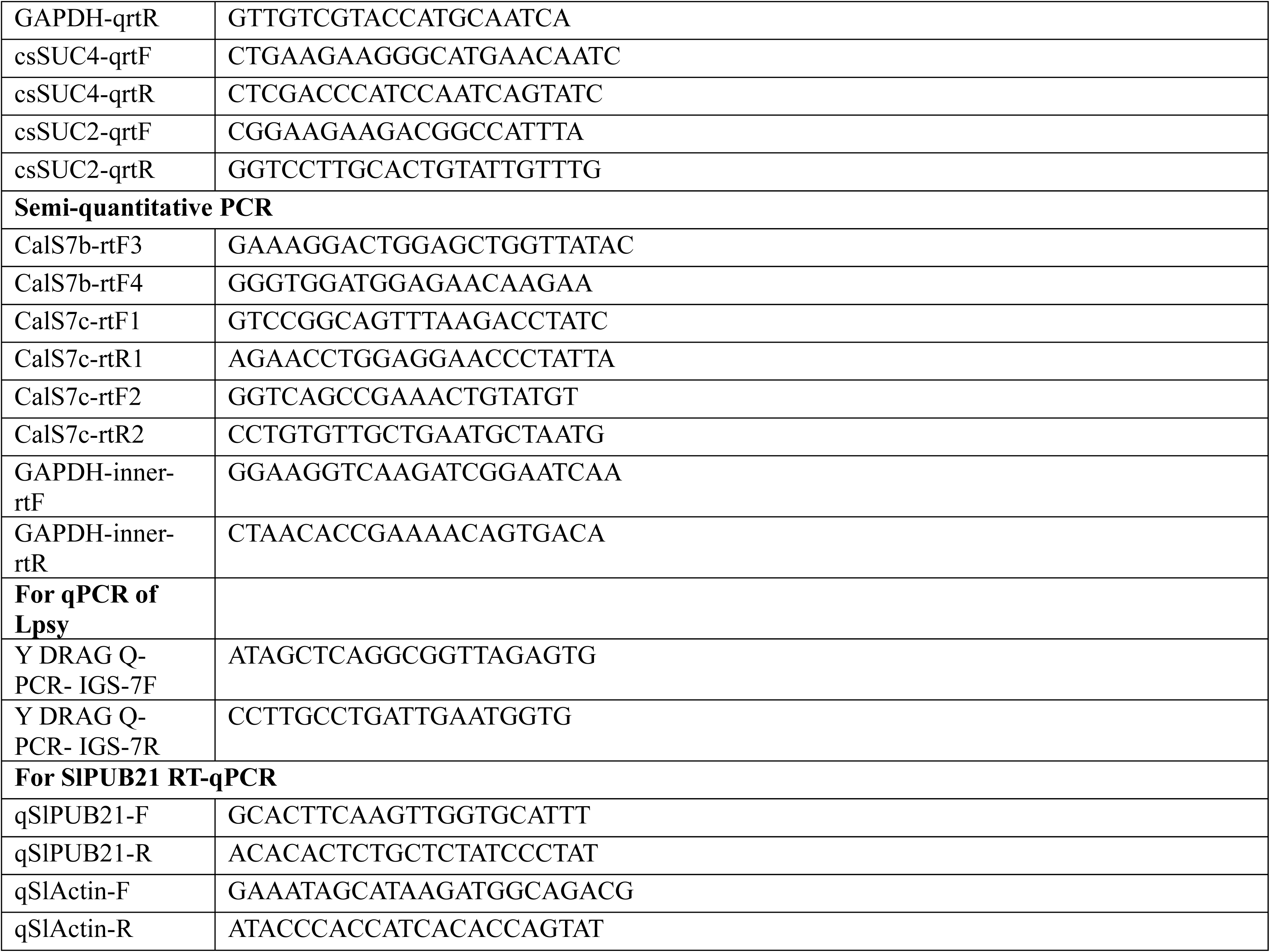
Primers, oligos, and gRNAs used in this study.

**Fig. S1:**
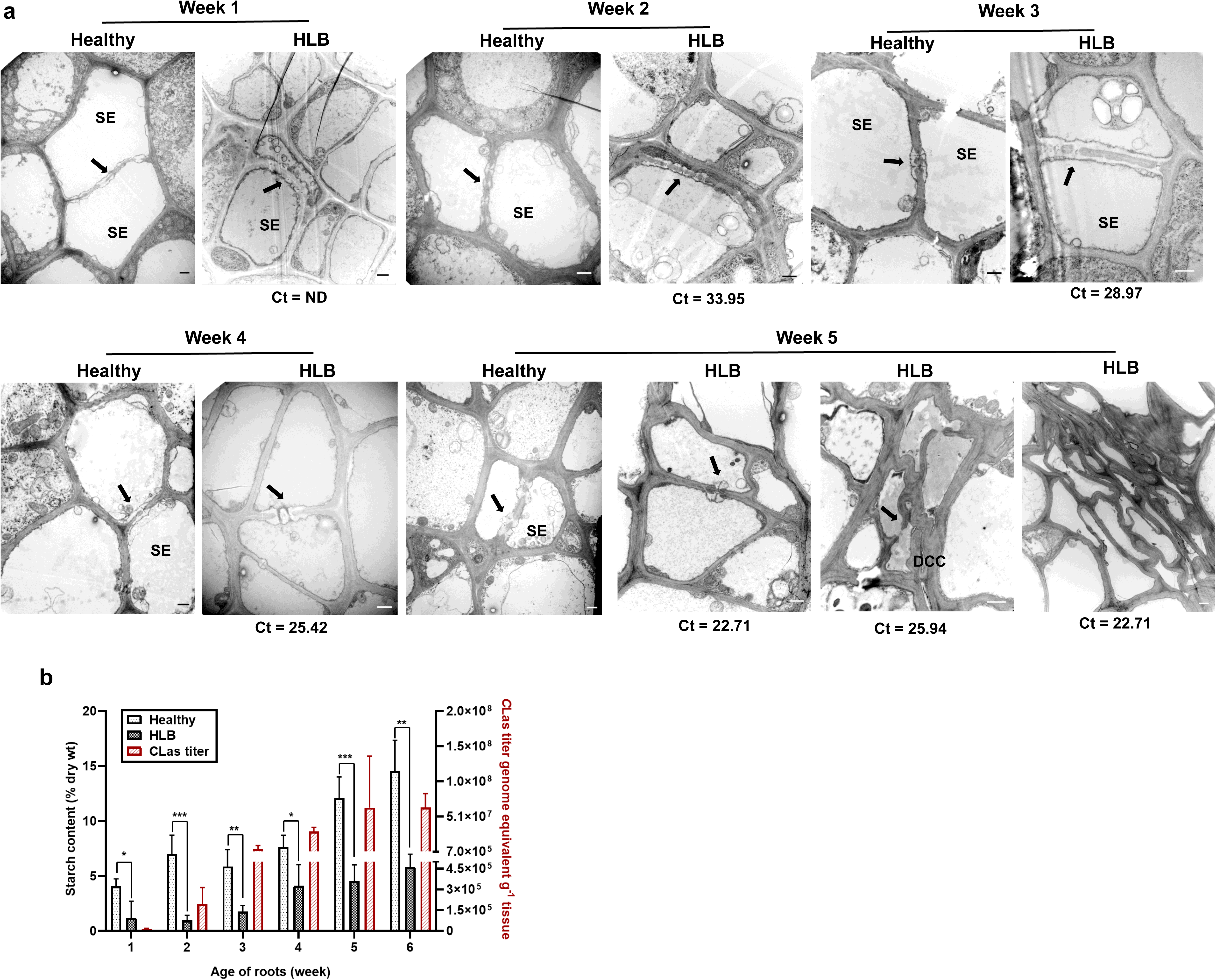
Comparison of newly generated roots of *C. sinensis* cv. Valencia scion on the Cleopatra mandarin rootstock. All young roots of approximately HLB symptomatic and healthy citrus plants were removed to promote root growth. Newly generated roots were tested for TEM, CLas titers and starch accumulation. (a) TEM analysis of newly generated roots. SE: sieve element, DCC: dead companion cells, arrow: sieve plate and sieve pores, (b) Starch accumulation. Mean + SD were shown. n=4.

**Fig. S2:**
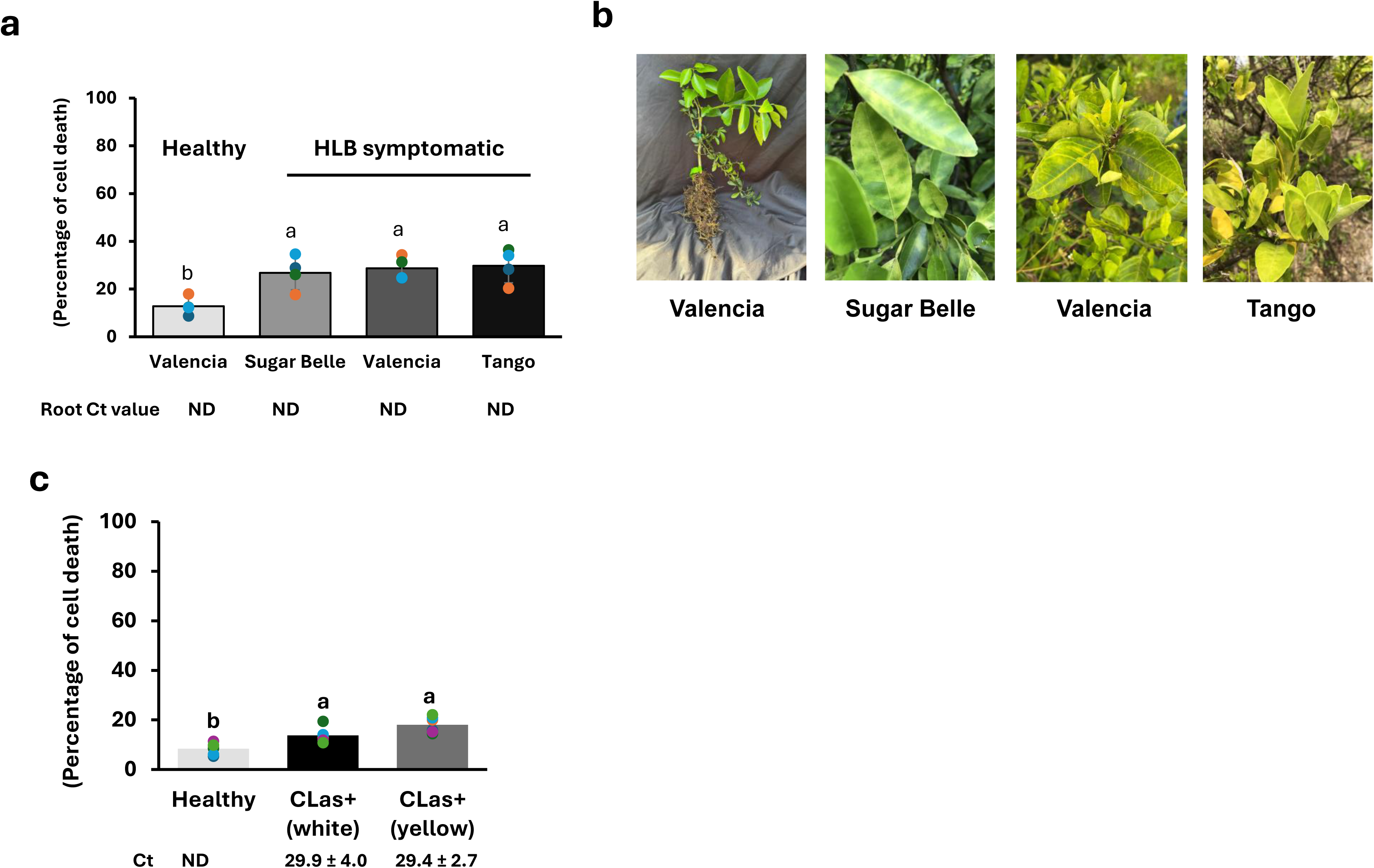
Ion leakage assays in citrus roots. (**a**) Ion leakage of US942 root. Three different scion varieties (*Citrus sinensis* cv. Valencia, Sugar belle and Tango) on US 942 rootstock were used for this assay. CLas infected roots samples were collected from the field and healthy root samples were collected from the screen house as a control. Non-degraded healthy-looking root samples were collected from HLB symptomatic trees. Ion leakage test was measured by using a CON 700 conductivity/°C/°F bench meter. The relative electrolyte leakage was represented in bar graph in percentage as a ratio of (live/total dead) x 100. The error bar represents the mean ± SD of (n=4). Statistical analysis was conducted by using one-way Anova Tukey HSD post hoc test (*P* < 0.05). Different letters indicate significant difference. (**b**) HLB symptoms of the sampled trees. (**c**) Ion leakage of Kuharske root. *C. sinensis* cv Valencia on Kuharske rootstock plants were used. White and yellow indicate the colors of the sampled roots with white roots generated later than yellow roots. CLas titers were determined by qPCR. ND: non-detected.

**Fig. S3:**
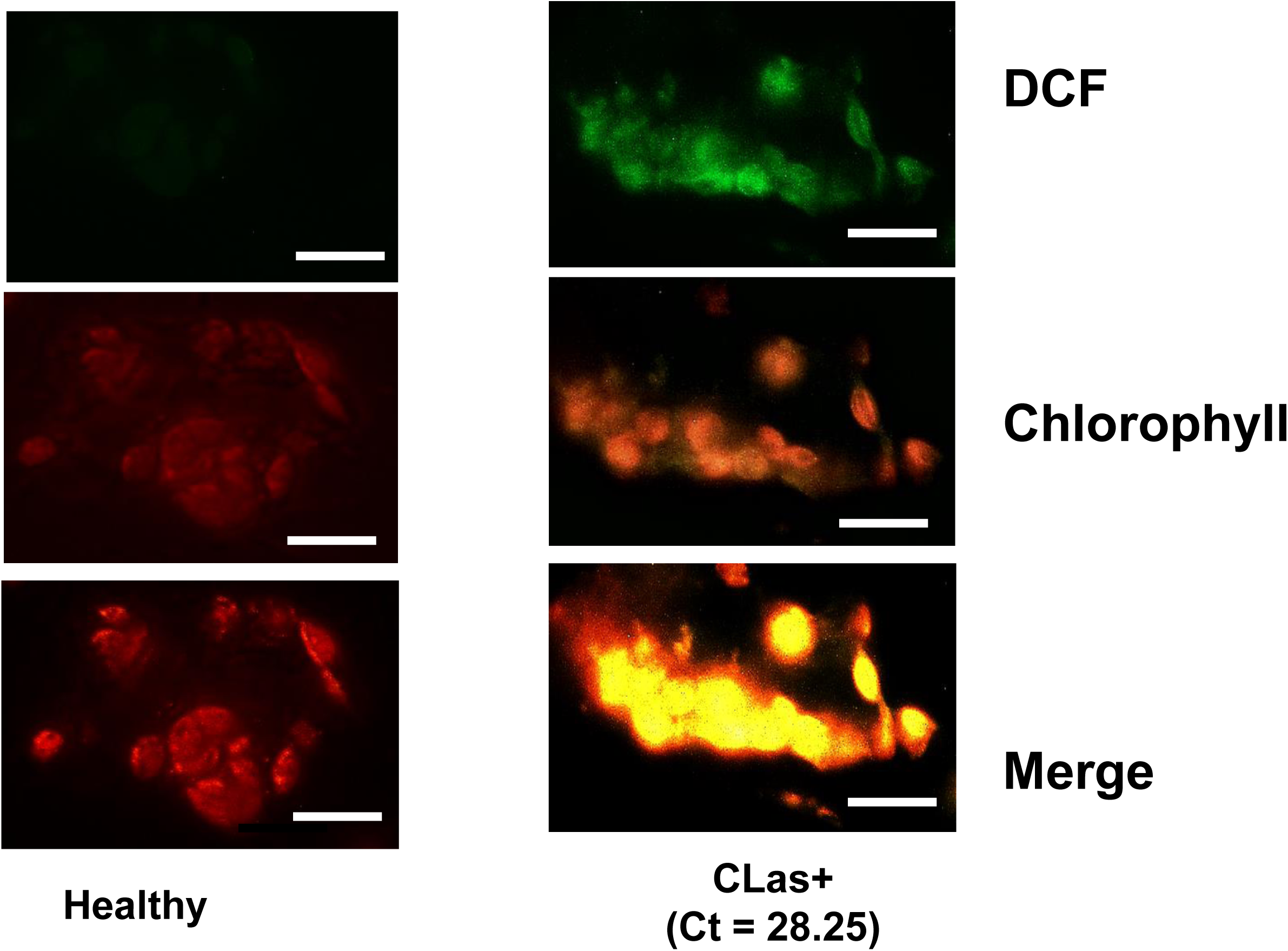
Subcellular total ROS detection in healthy and HLB symptomatic citrus leaves. Subcellular ROS were determined using the fluorescence of DCF, the oxidation product of H2DCF-DA. Protoplast cells were isolated from CLas infected and healthy Valencia mature leaves by enzymatic digestion with 1% cellulase RS and 1% macerozyme R-10. After 16 hours of enzyme digestion, protoplast cells were isolated and treated with 10 µm of H2DCF-DA then incubated at 37^0^C for 15 minutes in darkness. Treated protoplast cells were observed by fluorescence microscopy. Green signal indicates the GFP of DCF and red indicates the chloroplast autofluorescence. DCF: 2’,7’-dichlorofluorescein. Merge: chlorophyll + DCF. CLas titers were quantified by qPCR.

**Fig. S4:**
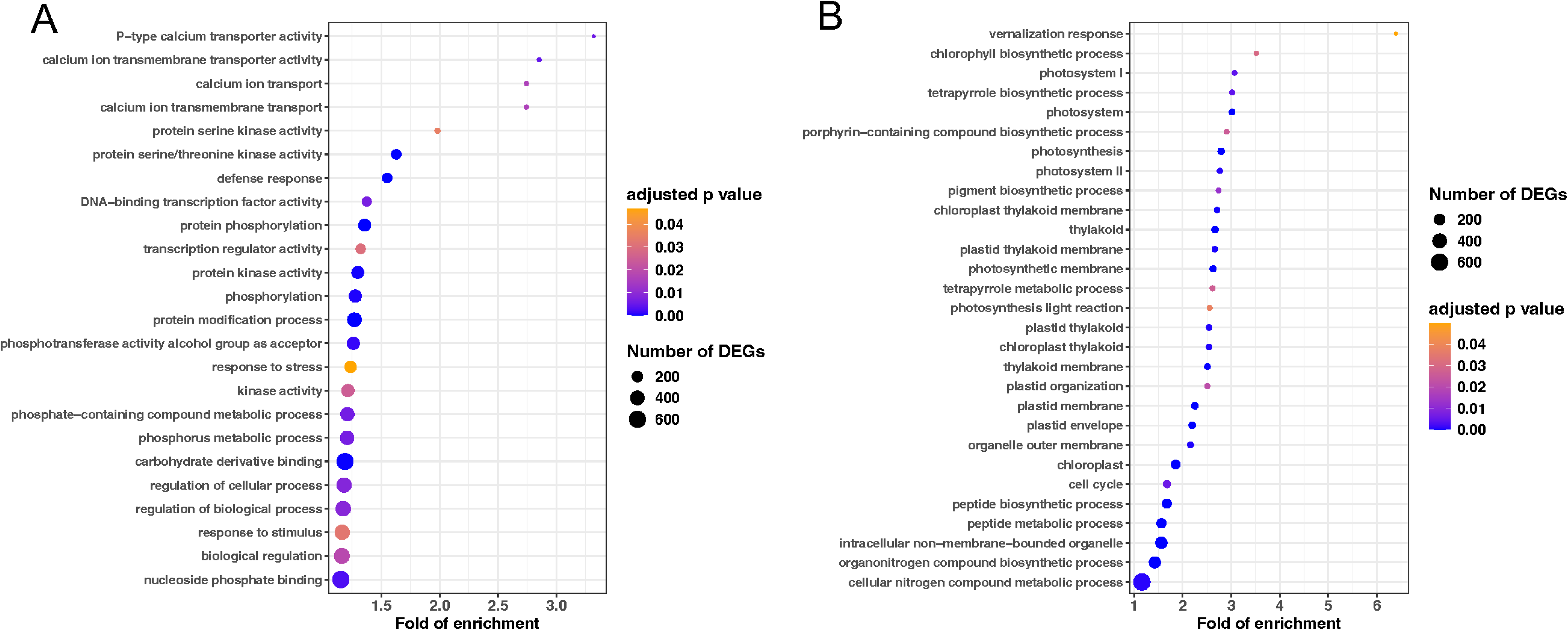
GO functional enrichment analysis of differentially expressed genes (DEGs) between HLB and healthy leaf for *Citrus sinensis*. (**a**) Upregulated genes for HLB v.s. Healthy. (**b**) Downregulated genes for HLB v.s. Healthy. DEGs were determined based on *P* ≤ 0.05 and showed the same trend at least 75% of significantly changed comparisons.

**Fig. S5:**
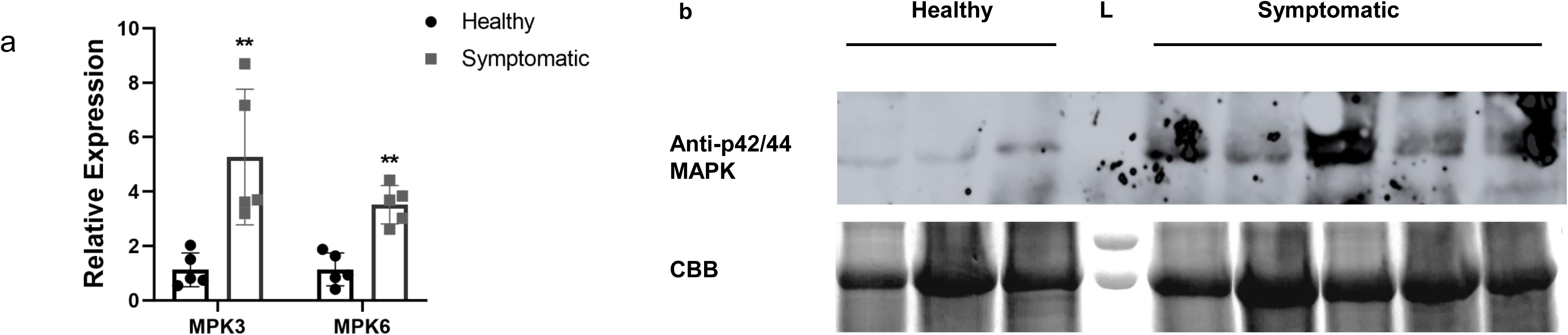
CLas activates MPK3*/*MPK6 in citrus. (**a**) Relative expression of *MPK3* and *MPK6* in healthy and CLas-positive *C. sinensis* cv. Valencia plants using quantitative reverse transcription PCR. The CsGAPDH gene was used as an endogenous control. Asterisks (*) indicate the statistically significant differences (*P* < 0.05, Student’s two-tailed *t*-test). (**b**) MAPK phosphorylation in healthy and symptomatic leaves in Valencia using anti-p42/44 MAPK immunoblotting. L, ladder; CBB, Coomassie Brilliant Blue.

**Fig. S6:**
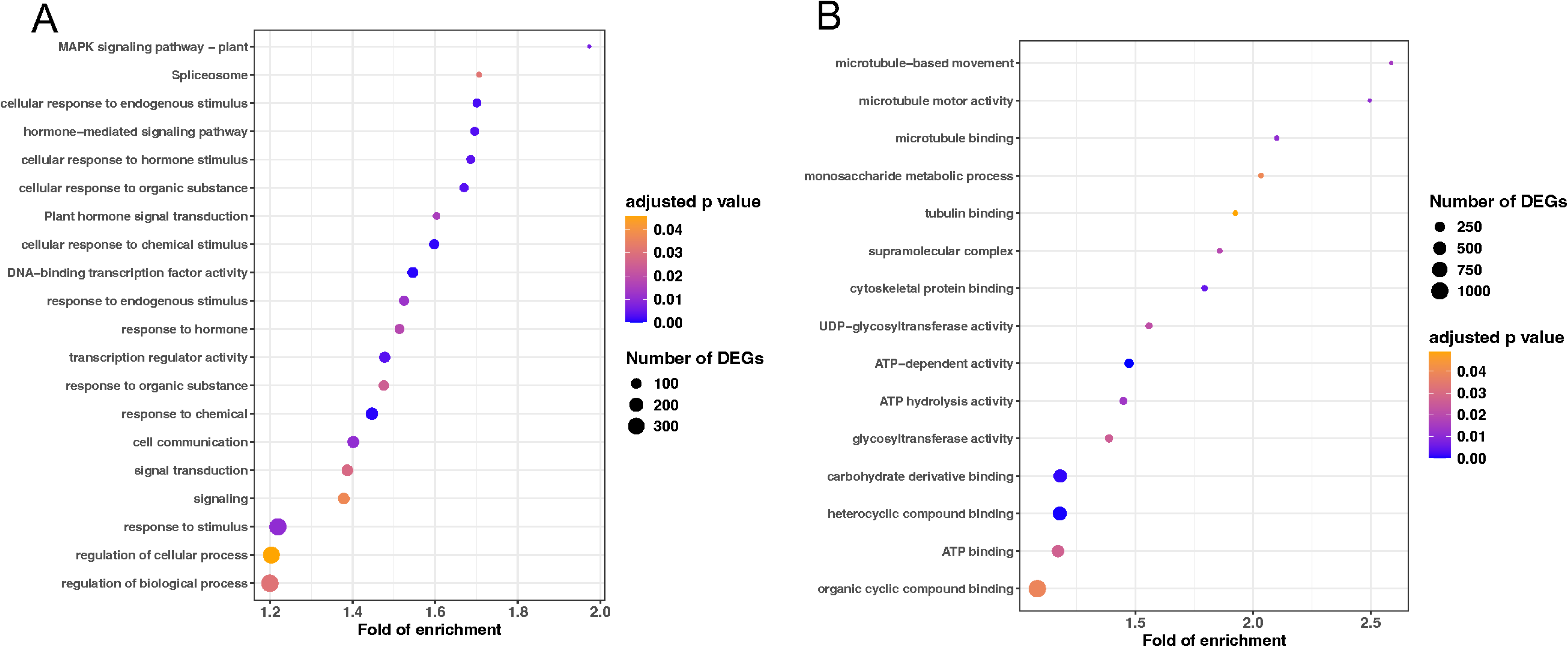
GO functional enrichment analysis of differentially expressed genes (DEGs) between HLB and healthy citrus root for multi citrus genotypes, such as Carrizo, Swingle citrumelo, Mandarin, and trifoliata orange. (A) Upregulated genes for HLB v.s. Healthy. (B) Downregulated genes for HLB v.s. Healthy. DEGs were determined based on the *P* ≤ 0.05 and showed the same trend at least 75% of significantly changed comparisons.

**Fig. S7:**
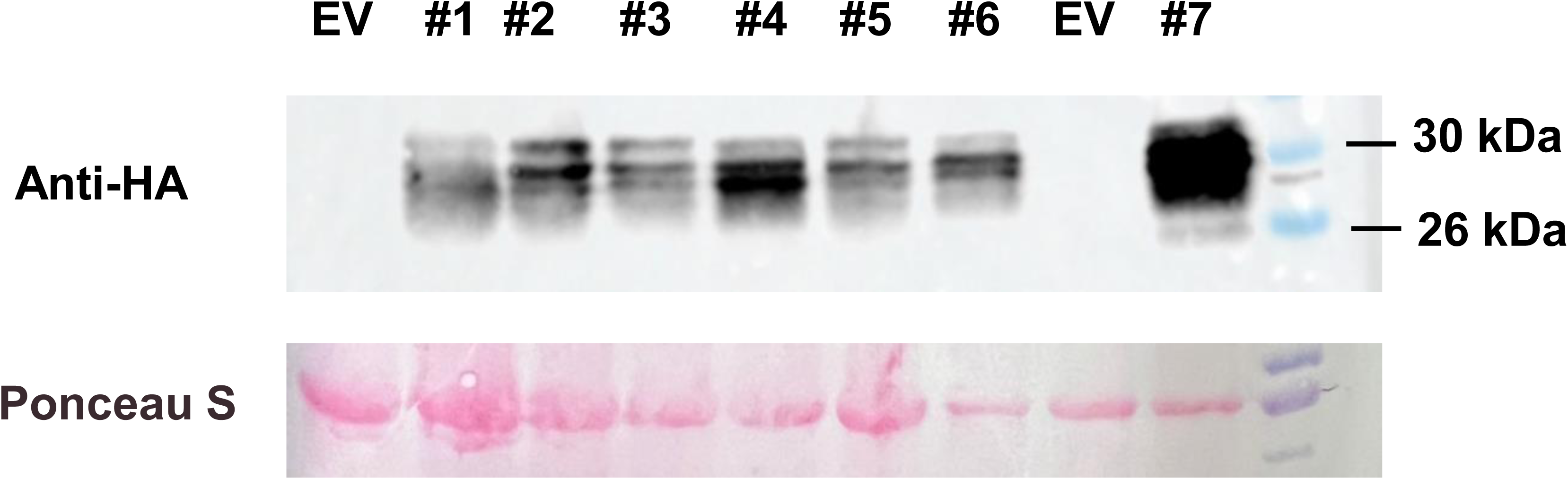
Confirmation of transgenic *35S::TP-Fld Citrus sinensis* cv. Hamlin plants using immunoblotting with the anti-HA antibody. Ponceau S–stained membrane was used as a loading control. Citrus plants transformed with an empty vector (EV) was used as the negative control.

**Fig. S8:**
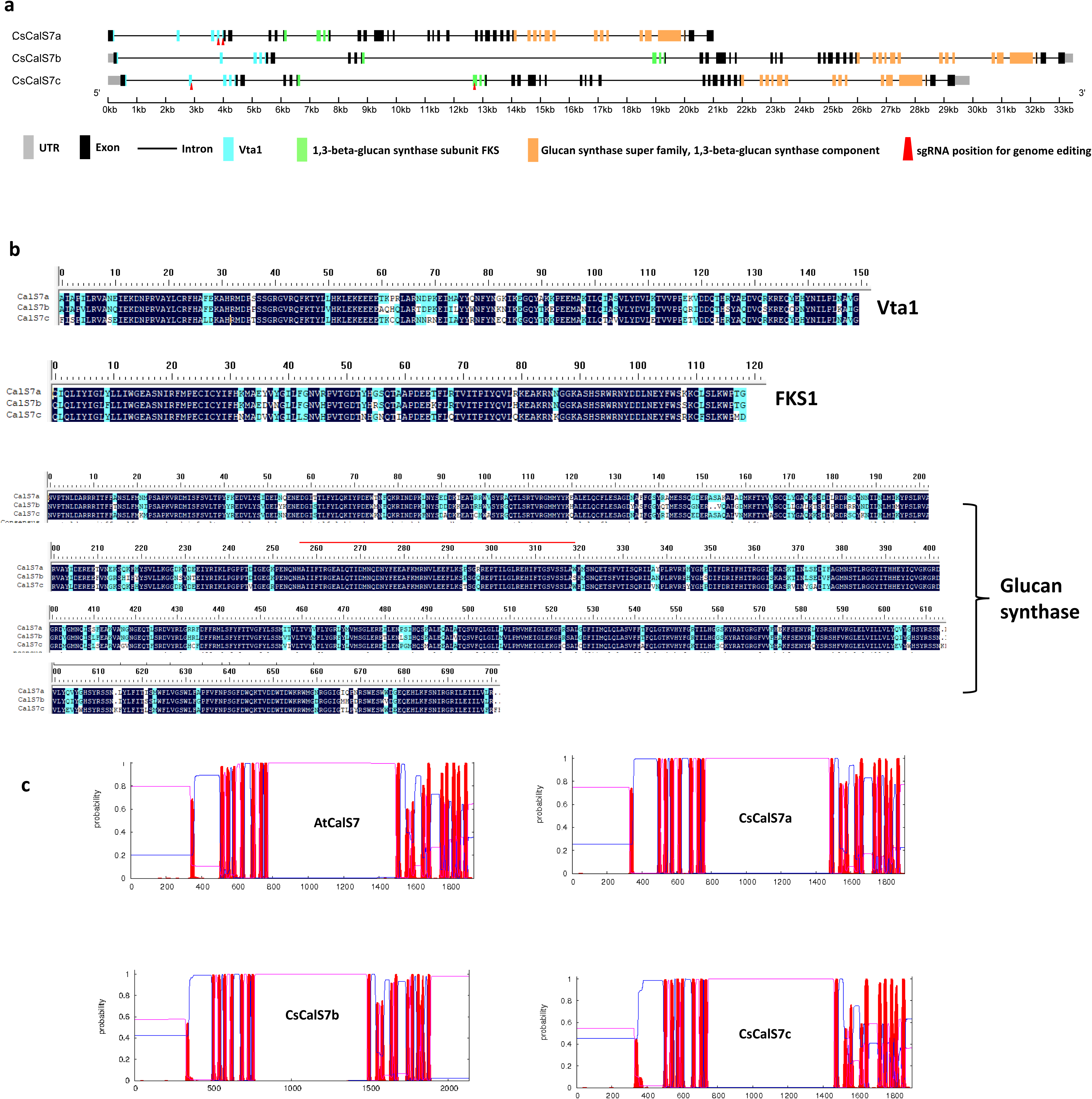
*CsCalS7* genes and proteins in *C. sinensis*. **(a)** Three *CsCalS7* homologs are present in *C. sinensis.* Grey bar, UTR; Black bar, Exon; Black line, Intron; Blue bar, Vta1 domain; Green bar, FKS domain; Orange bar, 1,3-beta-glucan synthase domain. (**b**) Protein alignment of Vta1, FKS1 and glucan synthase domains of three CsCalS7. (**c**) Prediction of CsCalS7 transmembrane motifs using https://services.healthtech.dtu.dk/service.php?TMHMM-2.0.

**Fig. S9:**
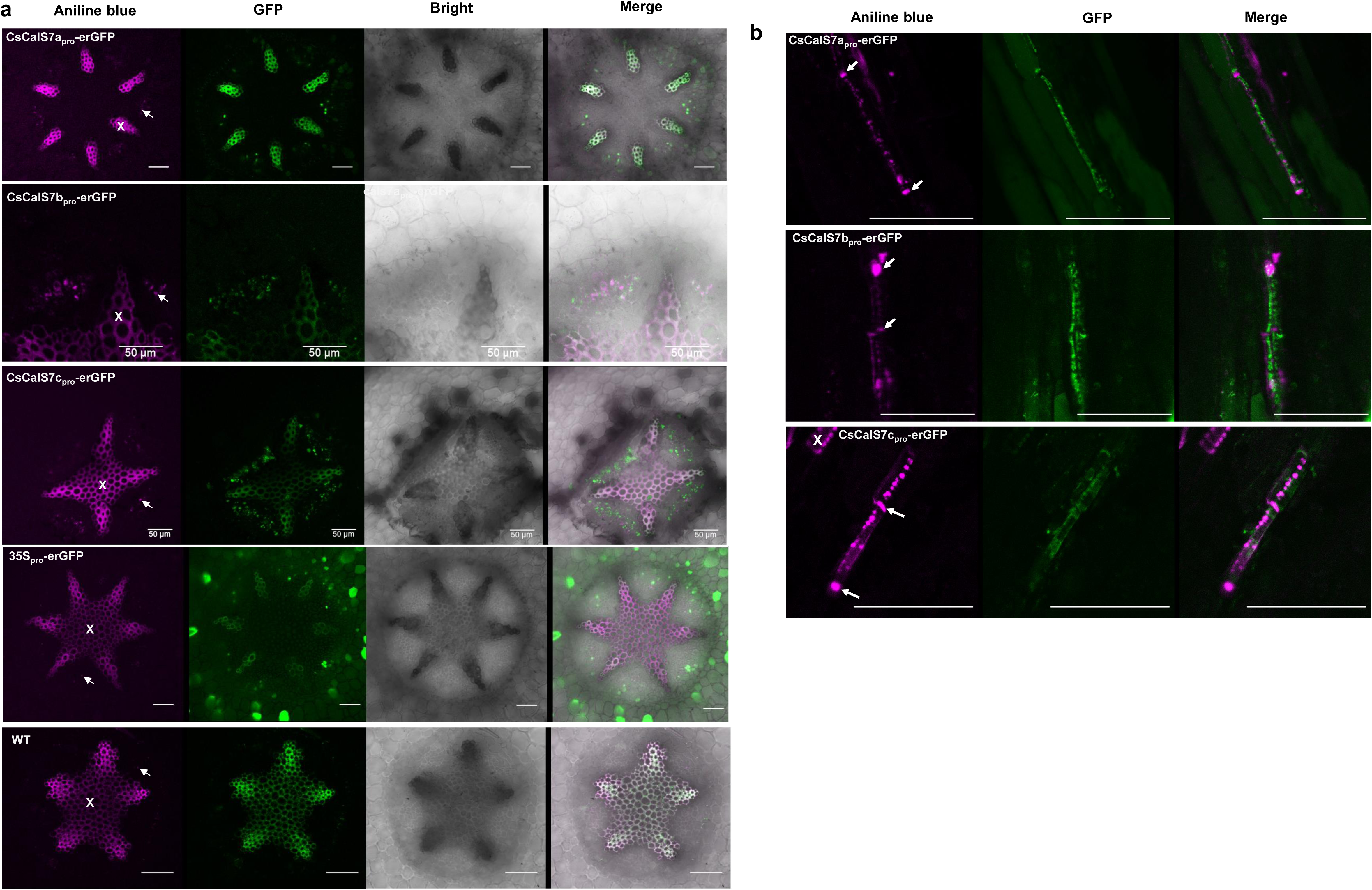
*CsCalS7*_pro_ driving erGFP are expressed in sieve elements. Root cross (**a**) and shoot longitudinal (**b**) sections were observed under confocal. Scale bars are 50 μm. X: xylem. Arrows in (a) indicate callose that deposit in phloem and in (b) indicate sieve plates.

**Fig. S10:**
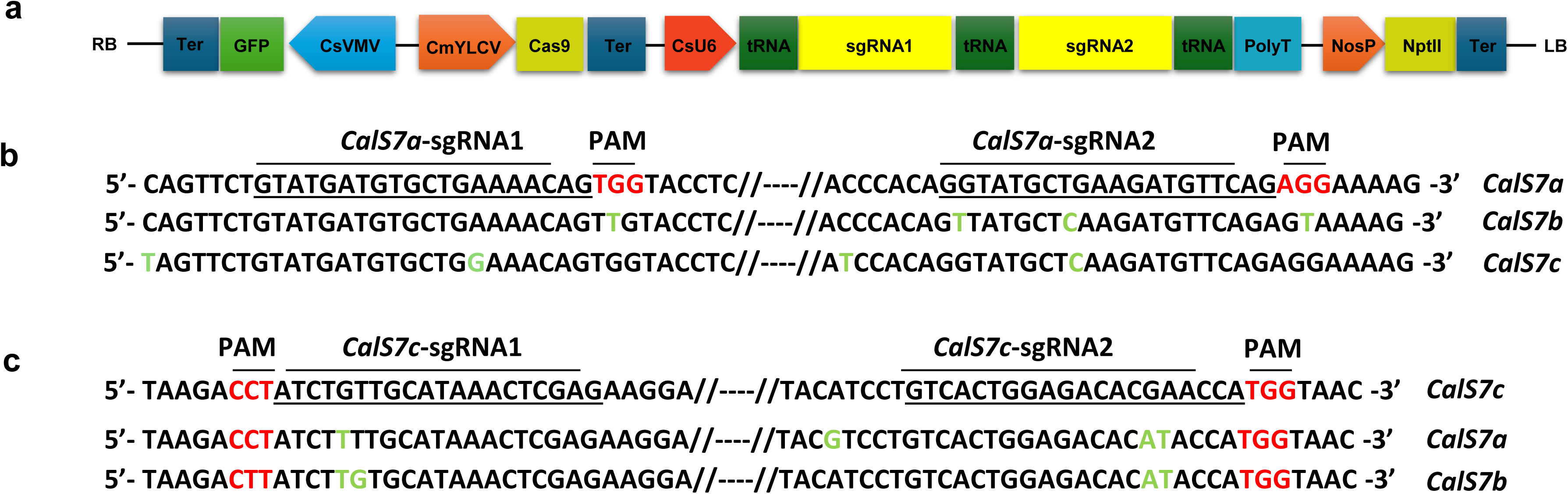
Plasmid construct and sgRNAs used for CRISPR mediated genome editing of *CsCalS7a* and *CsCalS7c*. (**a)** Schematic representation of plasmid construct used for *CsCalS7a* and *CsCalS7c* editing. sgRNAs of *CsCalS7a* (**b)** and *CsCalS7c* (**c)** are listed. sgRNAs are underlined, PAM is in red. The sequences corresponding to sgRNAs in the other two *CsCalS7* genes are listed below. Nucleotide variations are indicated in green.

**Fig. S11:**
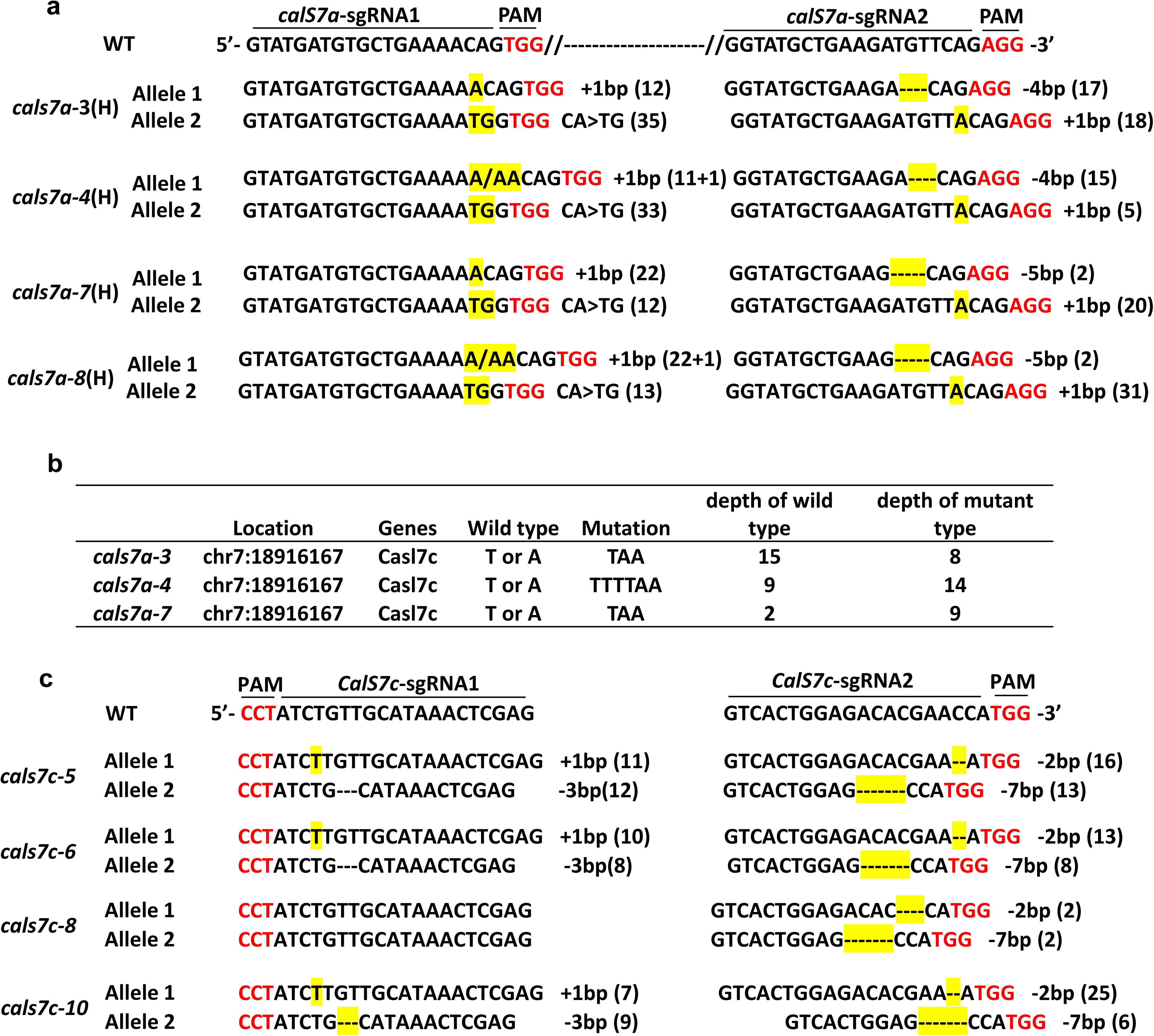
Representative *CsCalS7* genotypes of edited lines. (**a)** *CsCalS7a* genotyping of four genome edited lines. (**b**) Whole genome sequencing analysis of the off-target mutation on *CsCalS7c* in the cals7a *edited* plants. (**c)** *CsCalS7c* genotyping of four genome edited lines. The numbers in parentheses represent numbers of sequencing results.

**Fig. S12:**
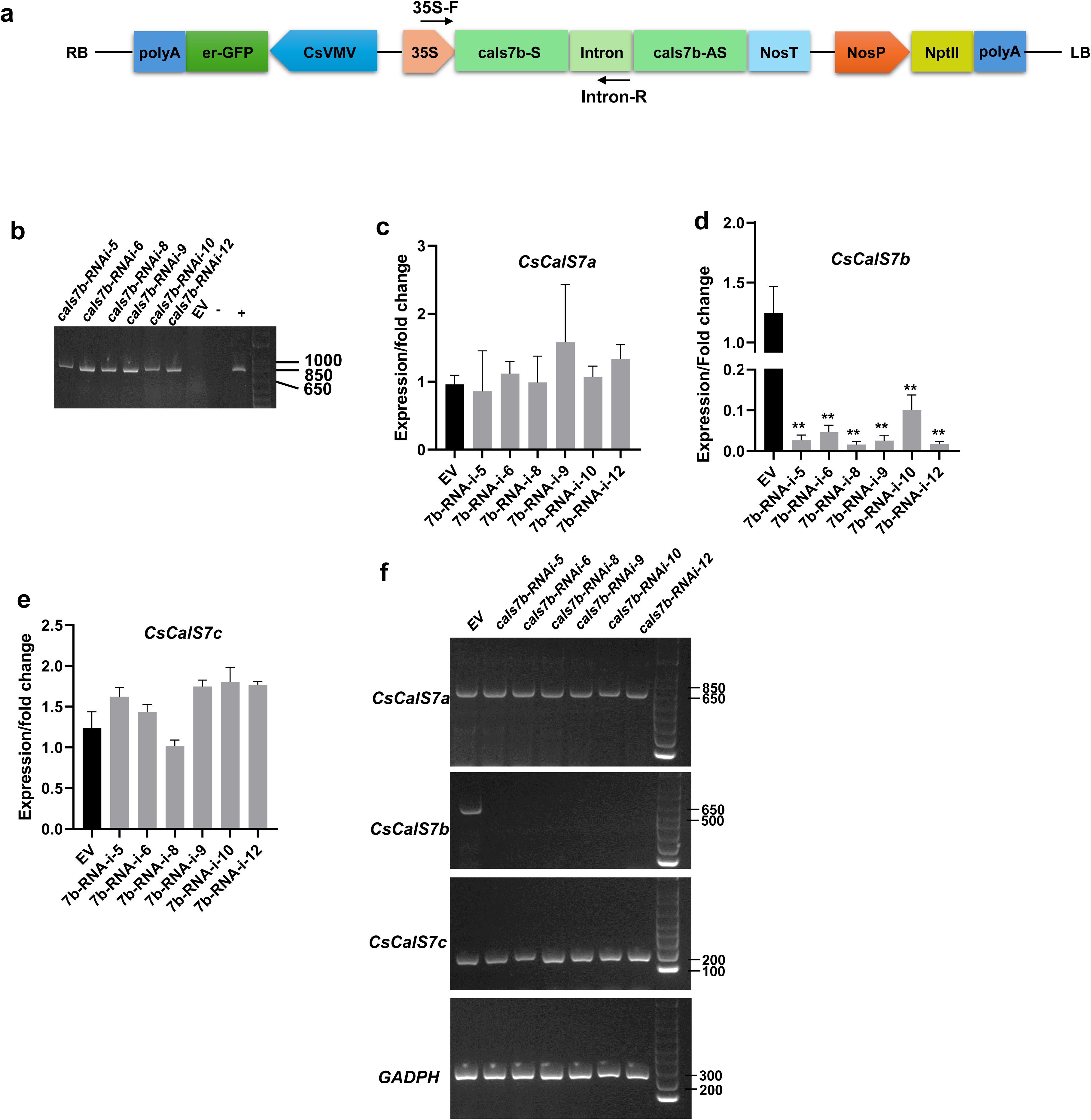
Generation of *CalS7b-RNAi* sweet orange plants. (**a)** Schematic representation of plasmid construct used for *CsCalS7b* silencing. *cals7b-S* is the sense strand while *cals7b-AS* is the antisense strand of targeted fragment of *CsCalS7b* cDNA. (**b)** Genomic DNA PCR to validate representative transgenic lines. Primers 35s-F and intron-R (arrows in panel A) were used. “-” indicates non template PCR control, “+” indicates positive control using plasmid as template. (**c-e)** qRT-PCR analysis of *CsCalS7a/b/c* relative expression levels in RNAi lines compared with that in EV transgenic plant. Mean ± SD (n=3) are shown. Experiments were repeated three times and representative results are shown. 3 independent EV lines were used and average value is presented. ** Indicates *P* value < 0.01, compared with EV. Student’s *t* test was used for statistical analysis. (**f)** Reverse transcription-PCR (RT-PCR) analysis of 3 *CsCalS7* genes using *GADPH* as an endogenous control.

**Fig. S13:**
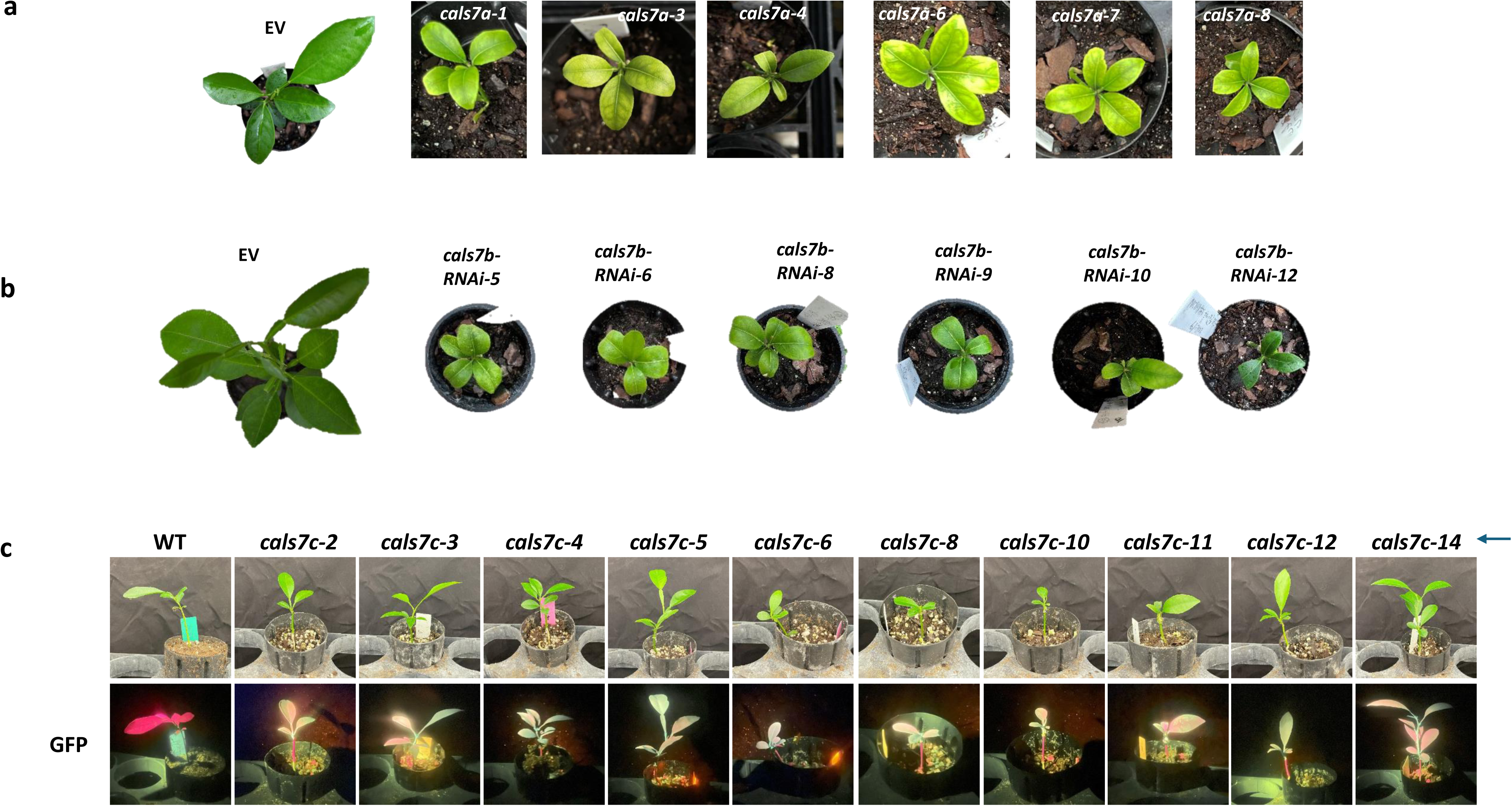
Phenotypes of loss of function mutants of *CsCalS7* genes in *C. sinensis* cv. Hamlin. (**a**) Representative *CsCalS7a* genome edited sweet orange plants at age of 4 months (time growing in soil pot). (**b**) Six months old (time growing in soil pot) *cals7b-RNAi* transgenic plants. EV are empty vector transgenic plants. (**c**) Representative *CsCalS7c* genome edited sweet orange plants at age of 2.5 months (time after micrografting).

**Fig. S14:**
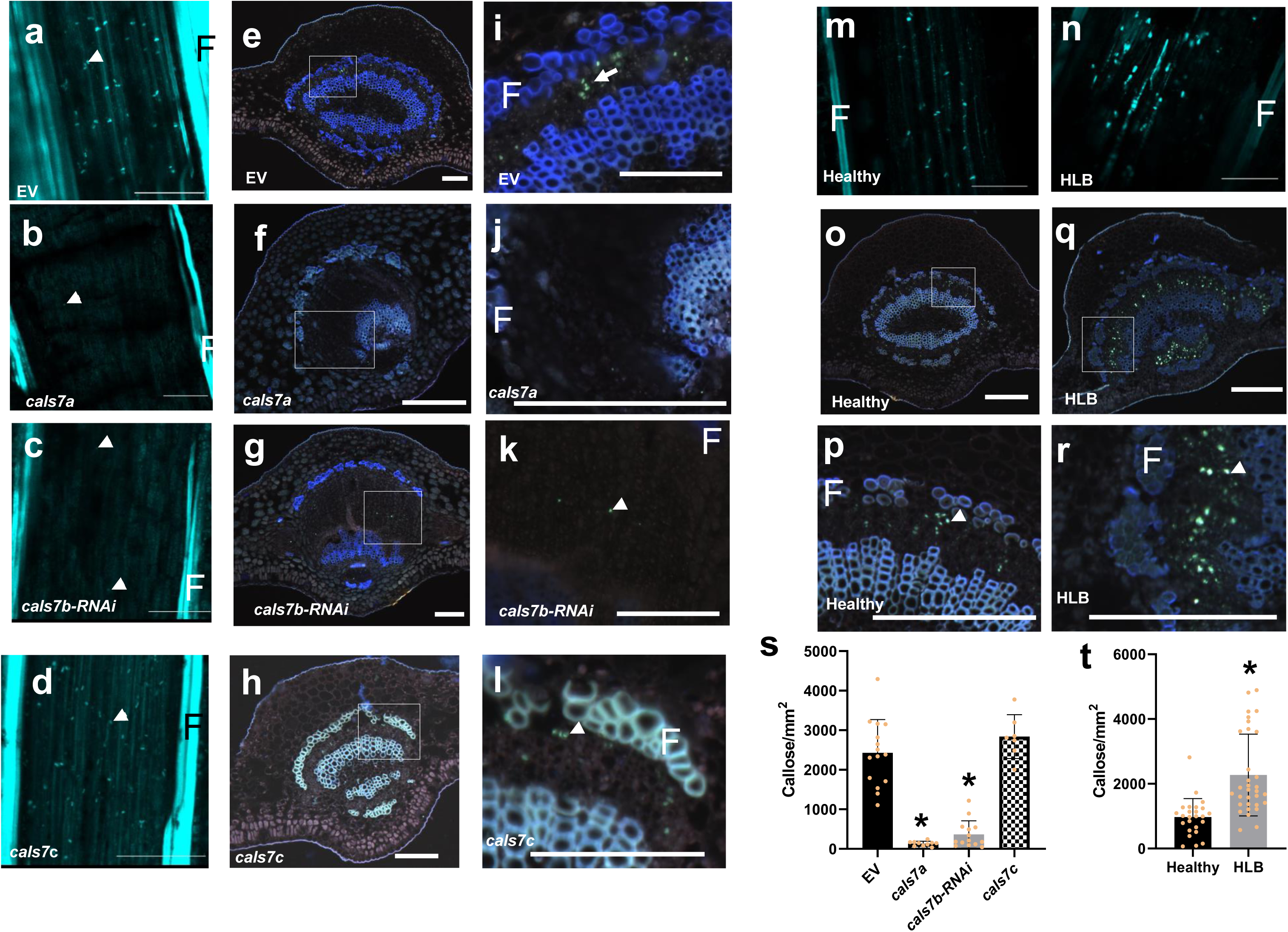
Callose staining in leaf midvein of *cals7a, CalS7b-RNAi* plants and *cals7c* mutant plants. 8 μm sections were stained with aniline blue. (**a-d)** Callose from longitudinal sections were viewed using confocal. Scale bars are 100 μm. (**e-l)** Callose in cross sections. Scale bars are 50 μm. (**i-l)** are magnified boxed-region from (**e-h)** respectively. (**m-n)** Callose from longitudinal sections of healthy and HLB leaf midveins. (**o-r)** are callose from cross sections of healthy and HLB leaf midveins. (**s, t)** callose quantification in longitudinal sections of EV and *cals7a* and *cals7c* mutants and *cals7b-RNAi* plants (**s**) and HLB plants (**t**). F indicates phloem fiber. White arrow heads indicate callose. Scale bars are 50 μm. * indicates *P* < 0.01 when compared with EV (**s**) or Healthy (**t**).

**Fig. S15:**
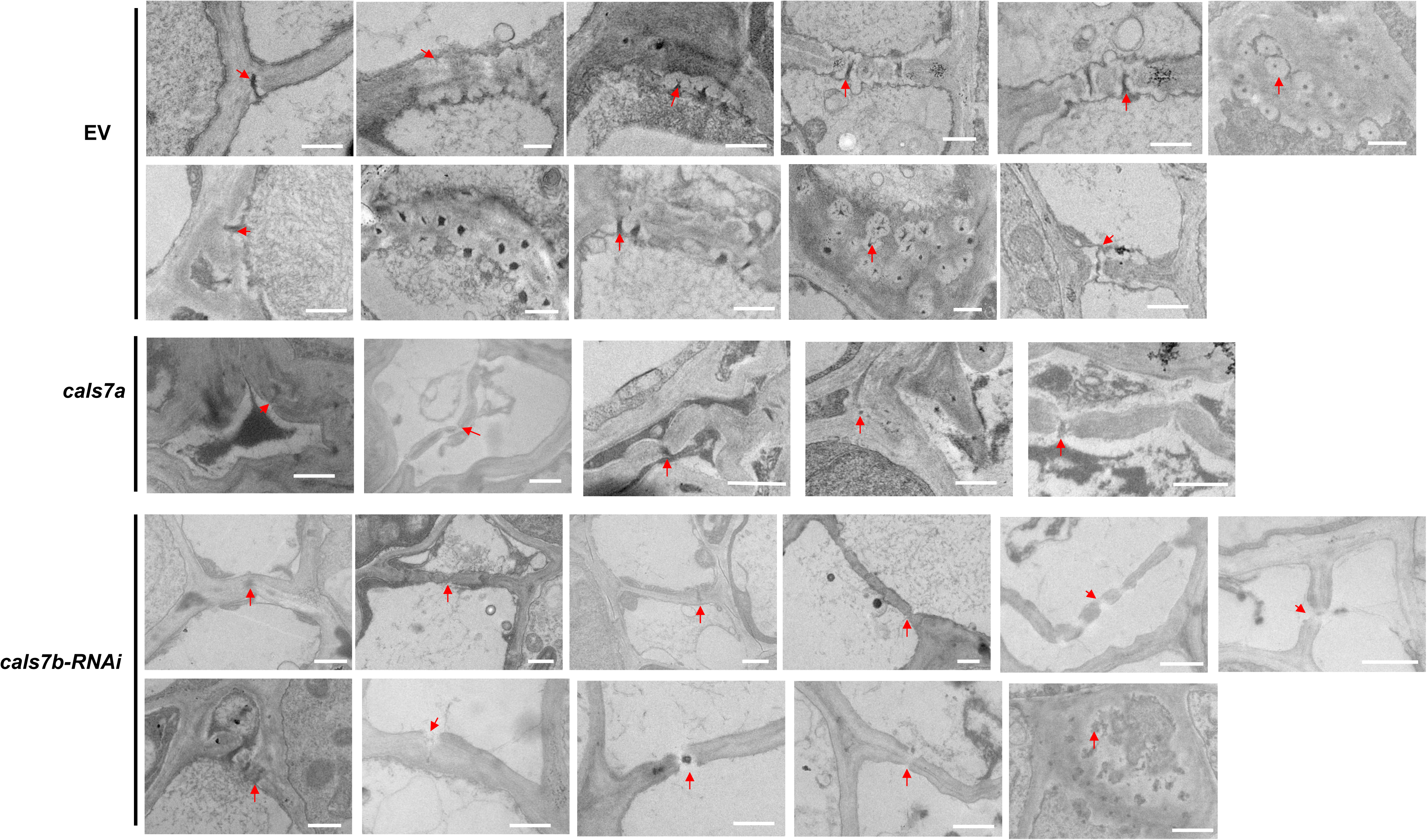
TEM micrographs showing sieve pores in midrib of young shoots of *C. sinensis* cv. Hamlin (at stage 1 in Fig.4). Red arrows indicate sieve pore. Scale bars are 1 μm.

**Fig. S16:**
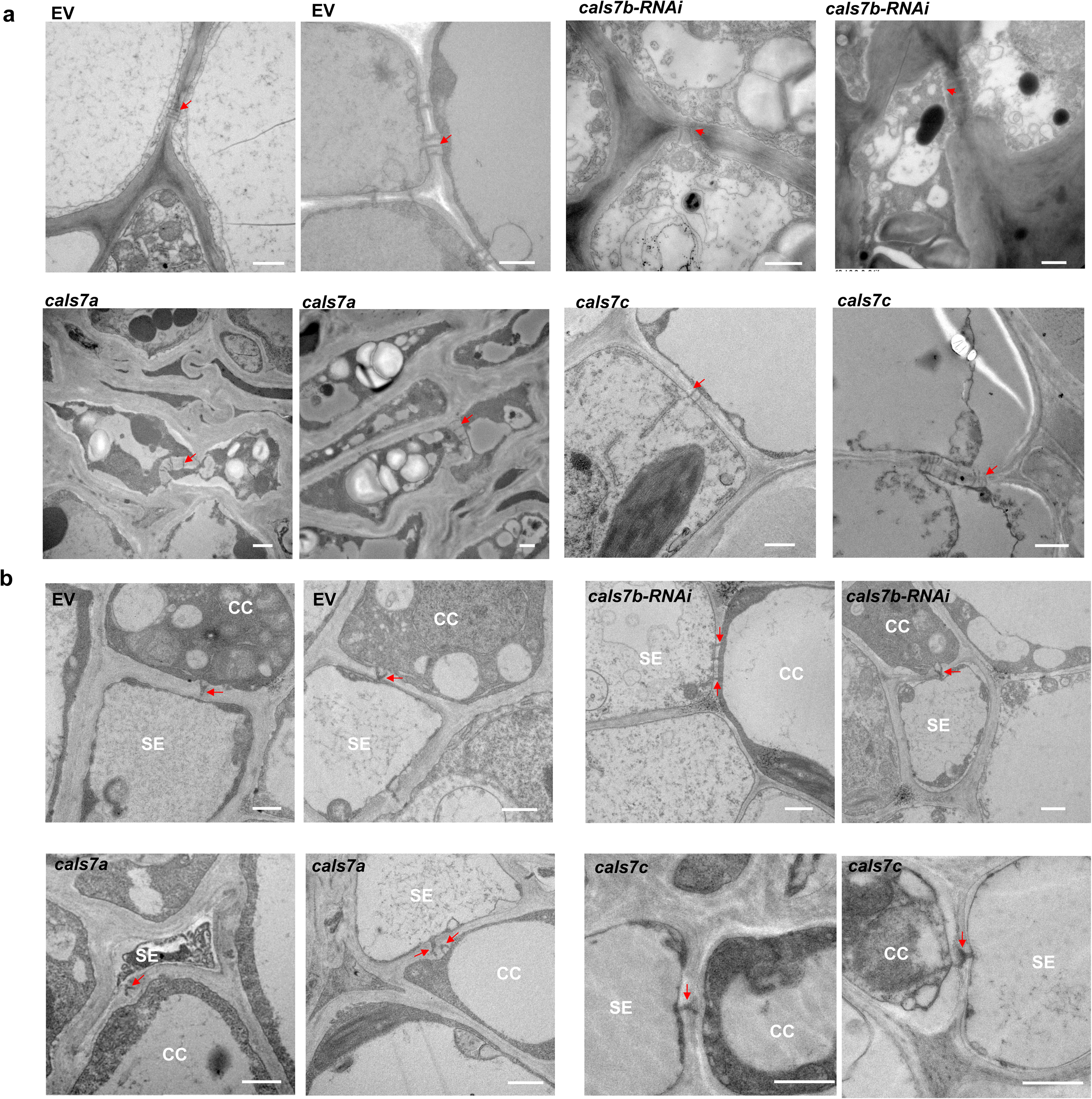
TEM micrographs of plasmodesmata. (**a**) Cross-sections of *C. sinensis* cv. Hamlin leaf midrib showing phloem parenchyma plasmodesmata (red arrows). Scale bars are 1 μm. (**b**) Transmission electron micrographs of cross-sections of *C. sinensis* cv. Hamlin shoot mid-rib showing plasmodesmata between sieve element and companion cell (red arrows). Scale bars are 1 μm.

**Fig. S17:**
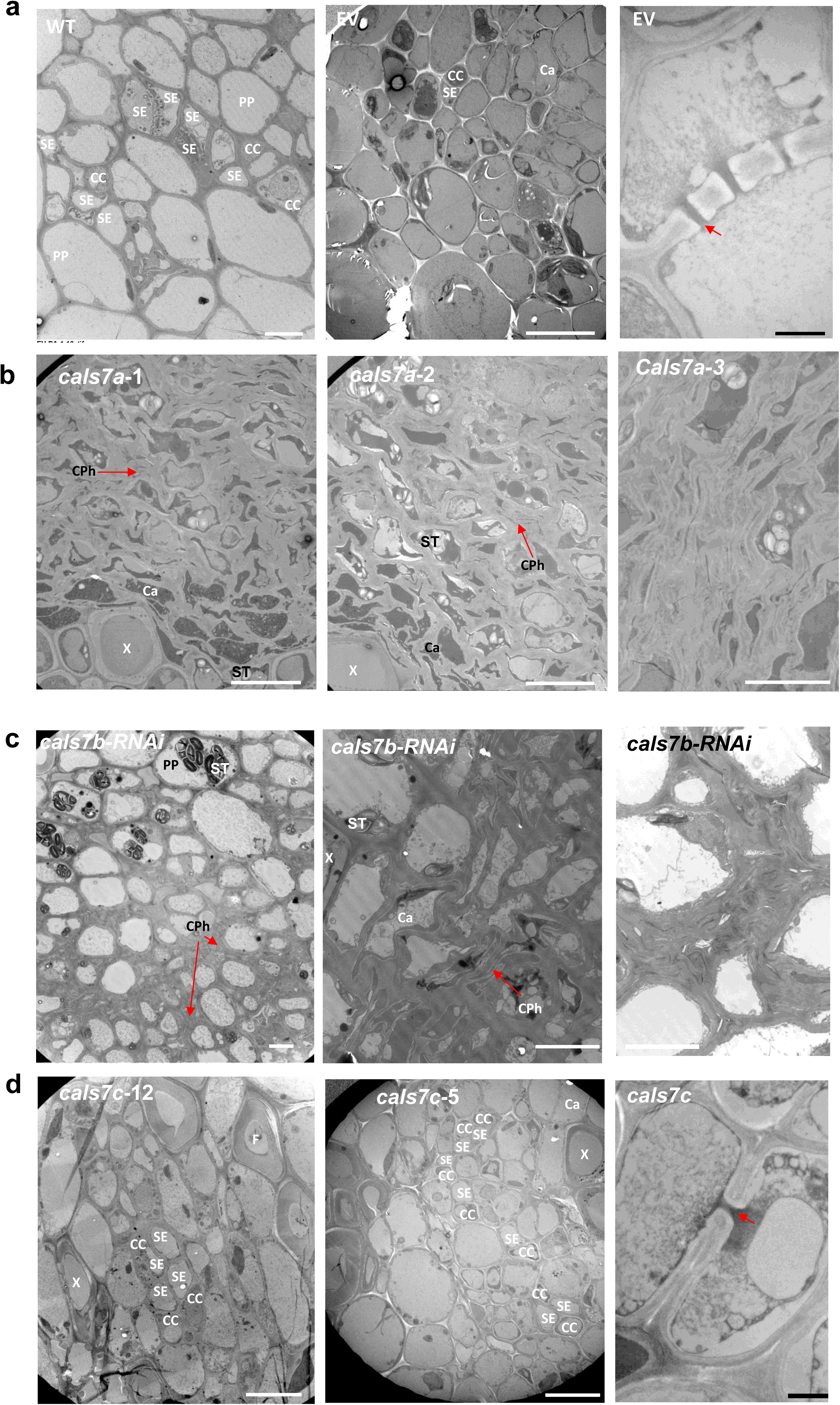
Transmission electron micrographs of *C. sinensis* cv. Hamlin mature leaf midrib. Phloem tissues of wild type (WT) and empty vector (EV) transgenic sweet orange cv. Hamlin **(a)**, genome edited *cals7a* mutant (**b**), *cals7b-RNAi* transgenic plants **(c)** and genome edited *cals7c* mutant (**d**). PP, phloem parenchyma; SE, sieve element; CC, companion cell; X, xylem; F, fiber; Ca, cambium; CPh, collapsed phloem cells; ST, starch. White scale bars are 10 μm and black scale bars are 1 μm.

**Fig. S18:**
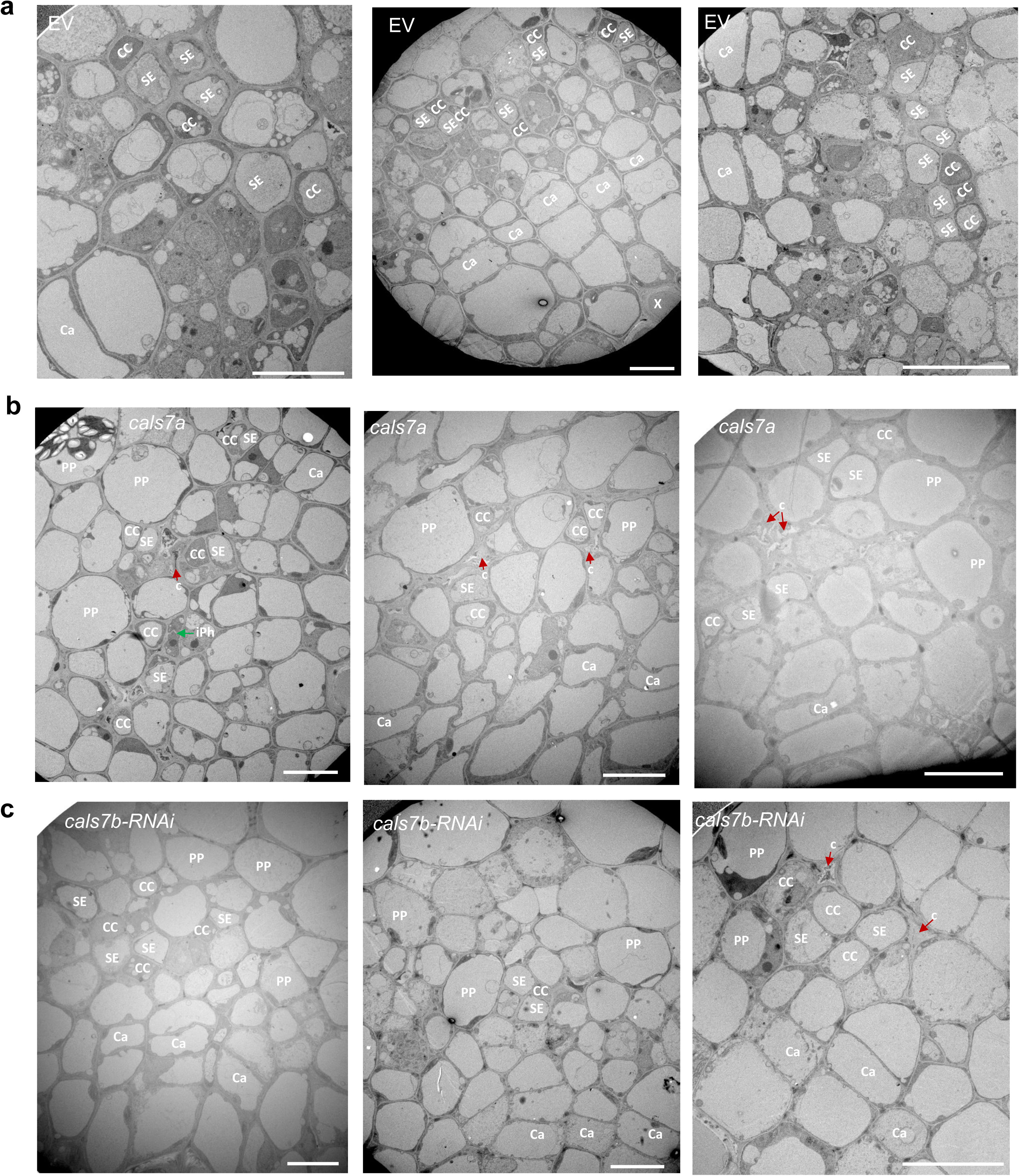
Wide view of stage-1 leaf midrib phloem under TEM. Phloem tissues of empty vector (EV) transgenic sweet orange cv. Hamlin **(a)**, genome edited *cals7a* mutant (**b**), *cals7b-RNAi* transgenic plants **(c)**. CC, companion cell; SE, sieve element; Ca, cambium; PP, phloem parenchyma cell; iPh, immature phloem cells; c, collapsed phloem cells or phloem cell death; Scale bars are 10 μm.

**Fig. S19:**
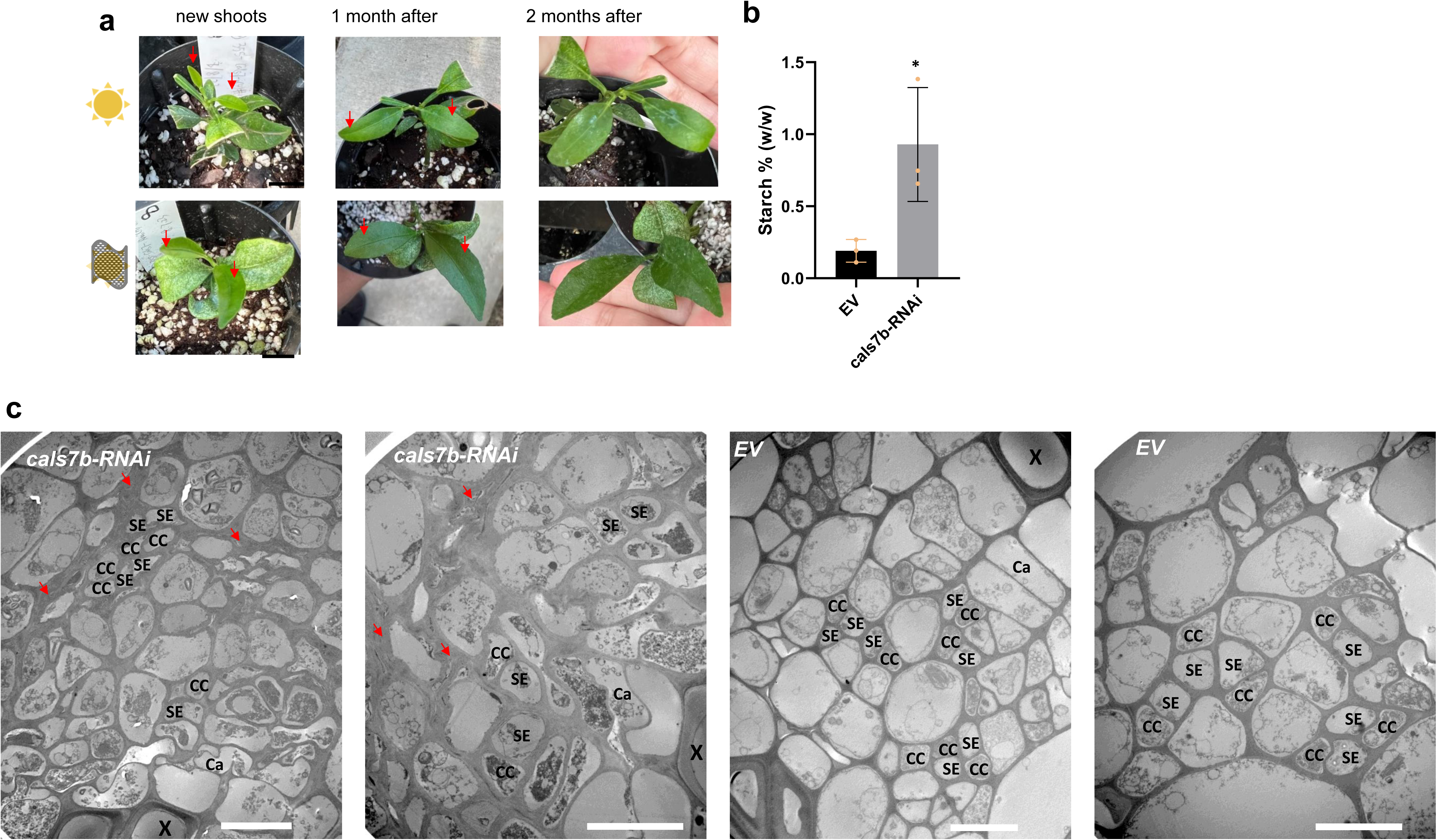
Shade treatment of *cals7b-RNAi* plants reduces starch accumulation but not phloem cell death. (**a**) Pictures of *cals7b-RNAi* plants under natural sun and 80% sunlight block. (**b)** Starch contents in plants under sunlight block for 2 months. * indicates *P* < 0.05. Students’ *t* test was used for statistical analysis. (**c**) Transmission electron micrographs of leaf mid vein of plants under 80% sunlight block. SE, sieve element; CC, companion cell; X, xylem; Ca, cambium. Arrows indicate collapsed cells or phloem cell death. Bars are 10 μm.

**Fig. S20:**
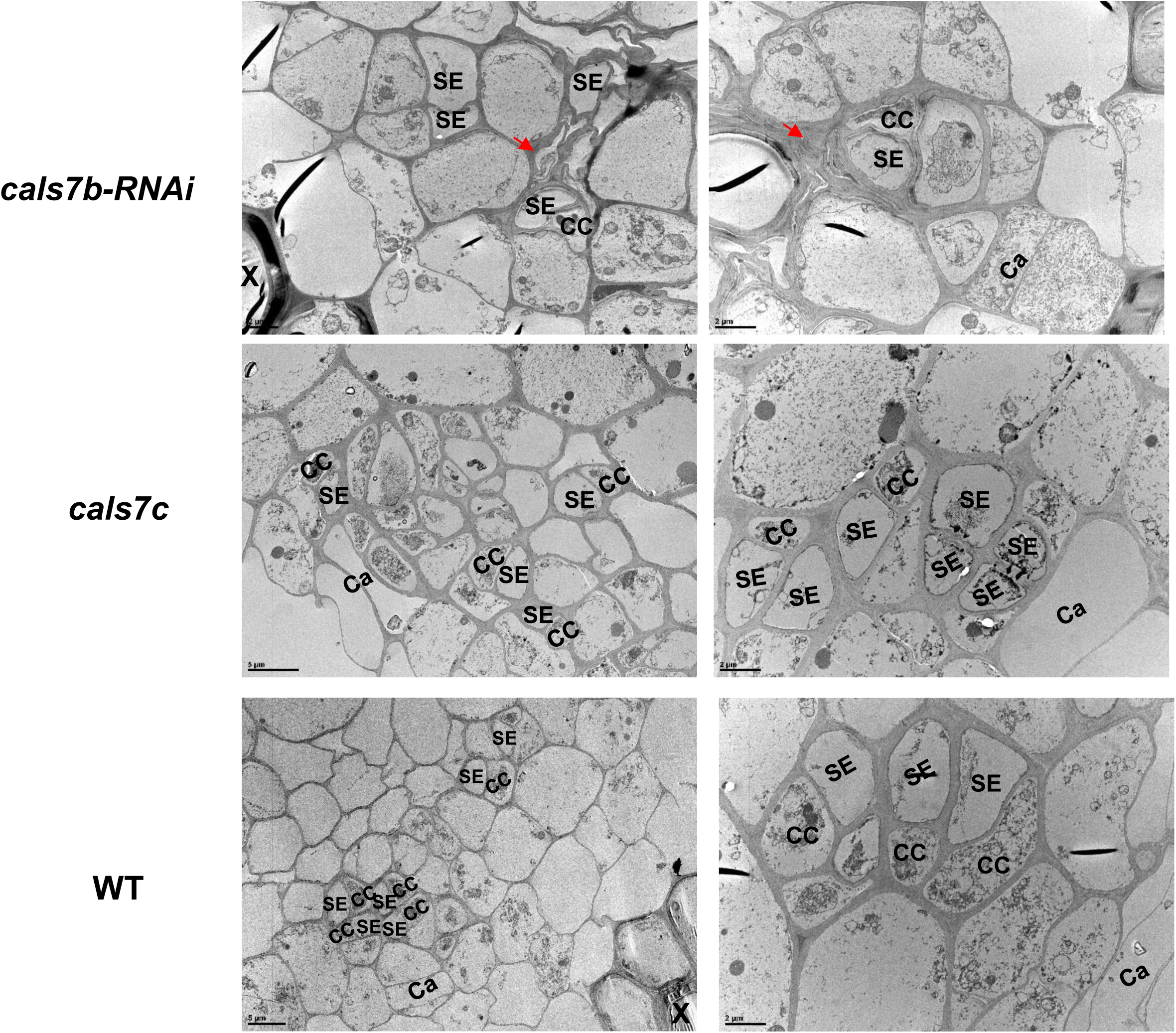
Transmission electron micrographs of cross-sections of regenerated root from *cals7b-RNAi*, *cals7c* edited and wild type (WT) *C. sinensis* cv. Hamlin branches. Red arrows indicate collapsed cells. SE, sieve element; CC, companion cell; X, xylem.

**Fig. S21:**
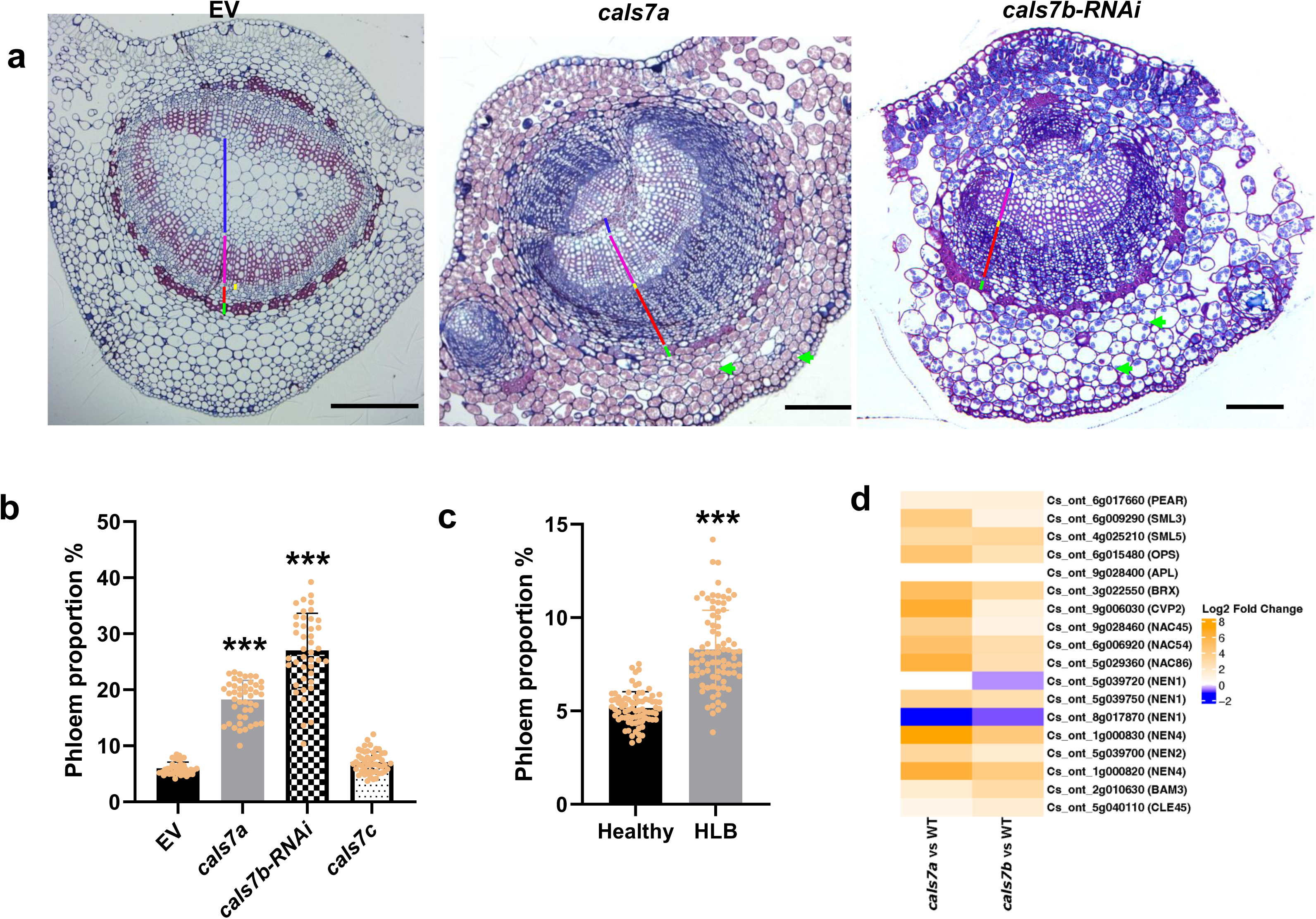
The midveins of *cals7a* mutant and *cals7b-RNAi* plants of *C. sinensis* cv. Hamlin generate more phloem. (**a**) Cross sections (1 µm) of leaf midvein were stained with methylene blue/Azure A and Basic fuchsin. Scale bars are 50 μm. Blue lines: pith. Pink lines: xylem. Yellow lines: cambium. Red lines: phloem. Green lines: fiber. Green arrows indicate starch. (**b**) Quantitative measurement of phloem proportion (phloem width/midvein diameter*100). (**c**) Quantitative calculation of phloem proportion in healthy and HLB mid vein. *** indicate significant difference at *P* < 0.0001 using student’s *t* test. (**d**) Heatmap of differently expressed genes that are involved in phloem differentiation in *cals7a* vs WT and *cals7b-RNAi* vs WT.

**Fig. S22:**
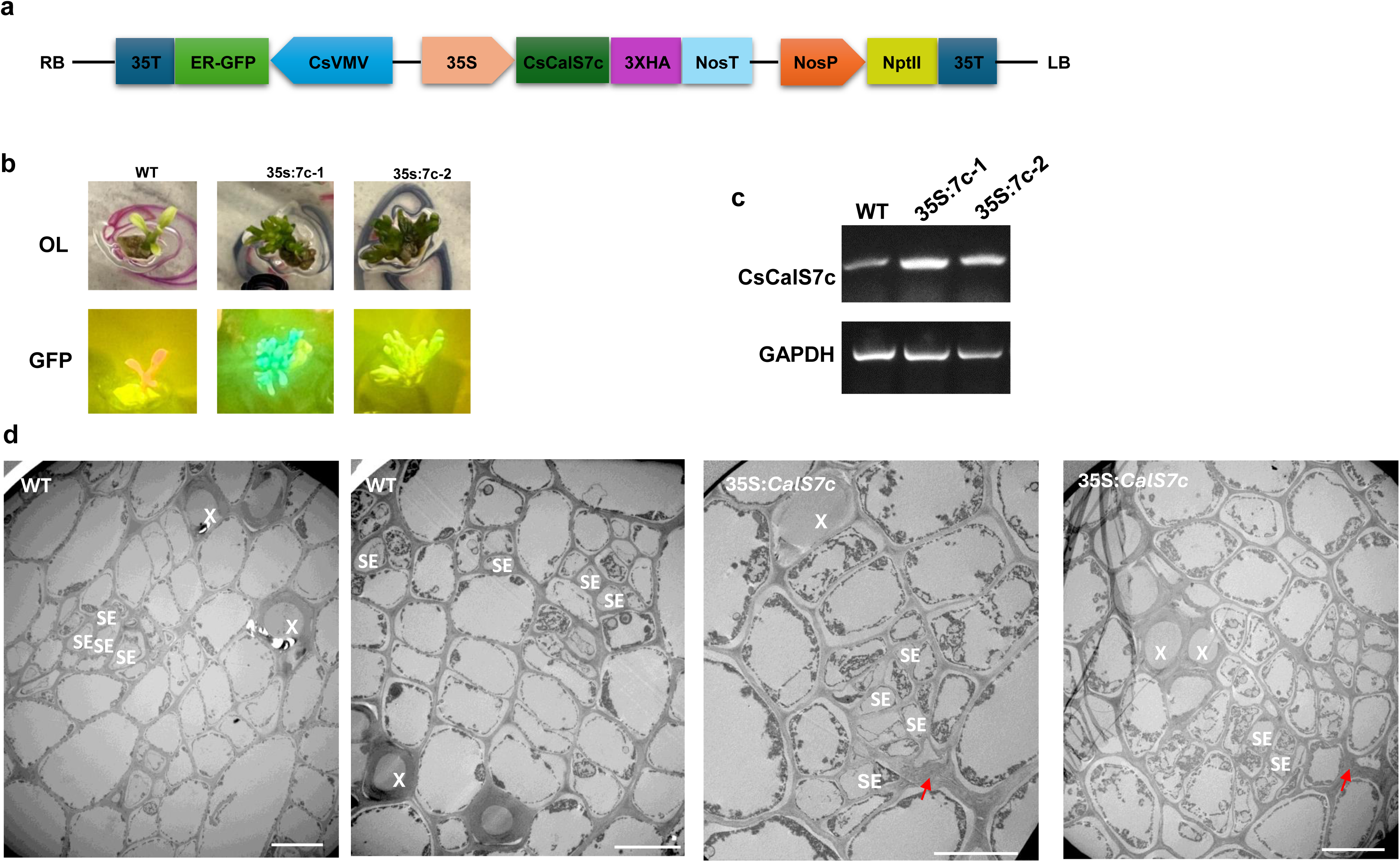
Overexpression of *CsCalS7c* driven by the 35s promoter in *C. sinensis* cv. Hamlin. (**a**) Illustration of construct used for overexpression of *CsCalS7c* driven by the 35s promoter. (**b**) GFP positive transgenic shoots grown on MS agar plates. (**c**) Semi-quantitative PCR using cDNA as the template with *CalS7c* primers. *GAPDH* is used as the endogenous control. (**d**) TEM images showing phloem cells of wide type and 35s:*CsCalS7c* shoots grown on MS agar plates. Arrows indicate collapsed phloem cells. Scale bars are 10 μm. X: xylem, SE: sieve element.

**Fig. S23:**
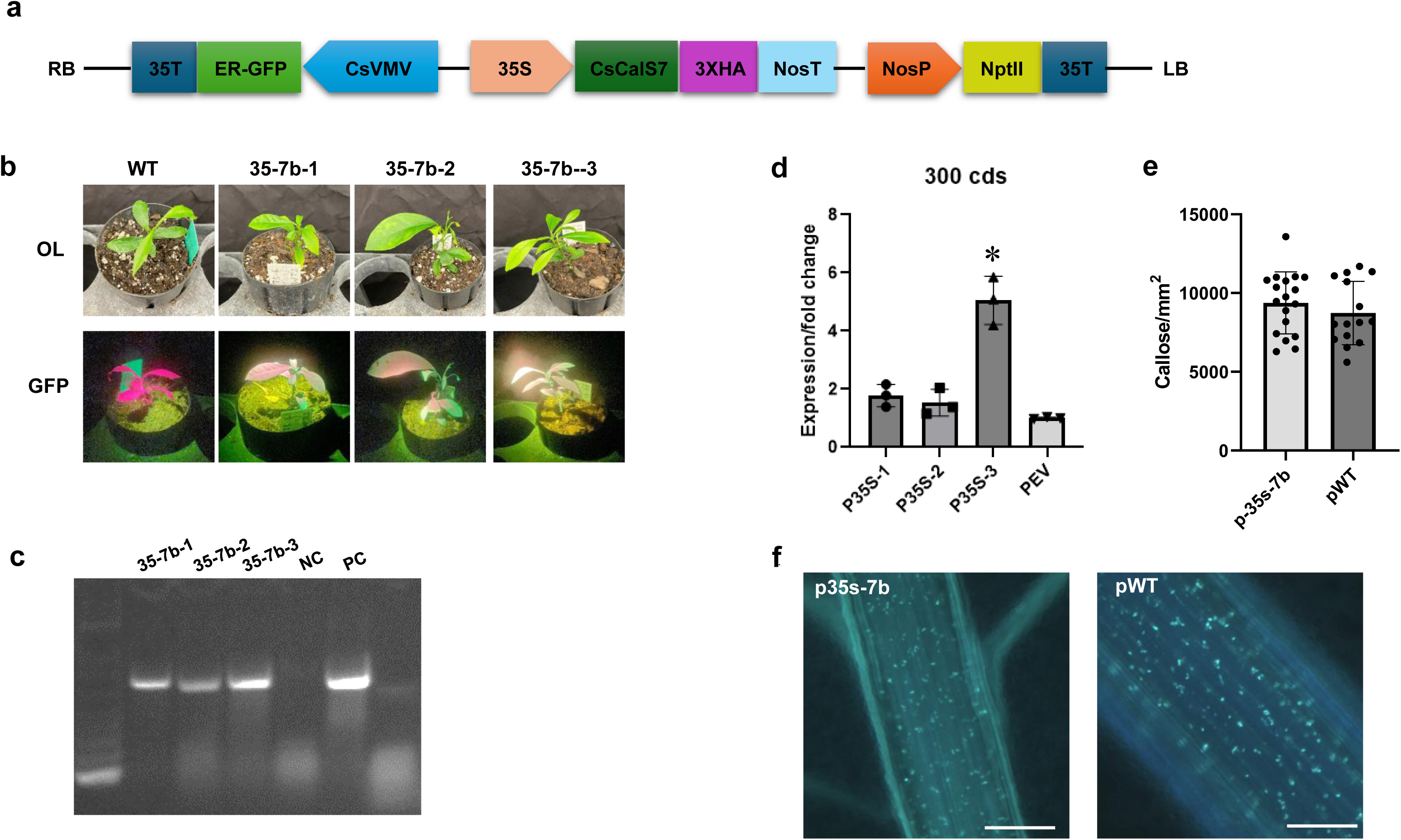
Overexpression of 35S promoter driving *CsCalS7b* in sweet orange (Pineapple). **(a)** Illustration of overexpression construct of the binary vector CsVMV-erGFP-pCAMBIA-1380N-35S for citrus. (**b).** Transgenic *C. sinensis* plants overexpressing *CsCalS7b* driven by the 35S promoter. WT, wild-type plant without GFP fluorescence signal. Top, under the optical light; bottom, erGFP fluorescence screening positive transgenic citrus under GFP light. (**c)** PCR validation for *CsCalS7b* with the 35S specific forward primer and gene-specific reverse prime. The plasmid was used as a positive control. (**d)** The relative expression level of *CsCalS7b* was tested by RT-qPCR. *CsGAPDH* was used as endogenous control. (**e)** Callose quantification in 35S-*CsCalS7b* (p-35s-7b) transgenic and WT leaf midvein. (**f)** Pictures showing callose deposition in leaf midvein. Scale bars are 50 μm. Asterisks (*) indicated the statistically significant differences (*P* < 0.05, Student’s two-tailed *t*-test).

**Fig. S24:**
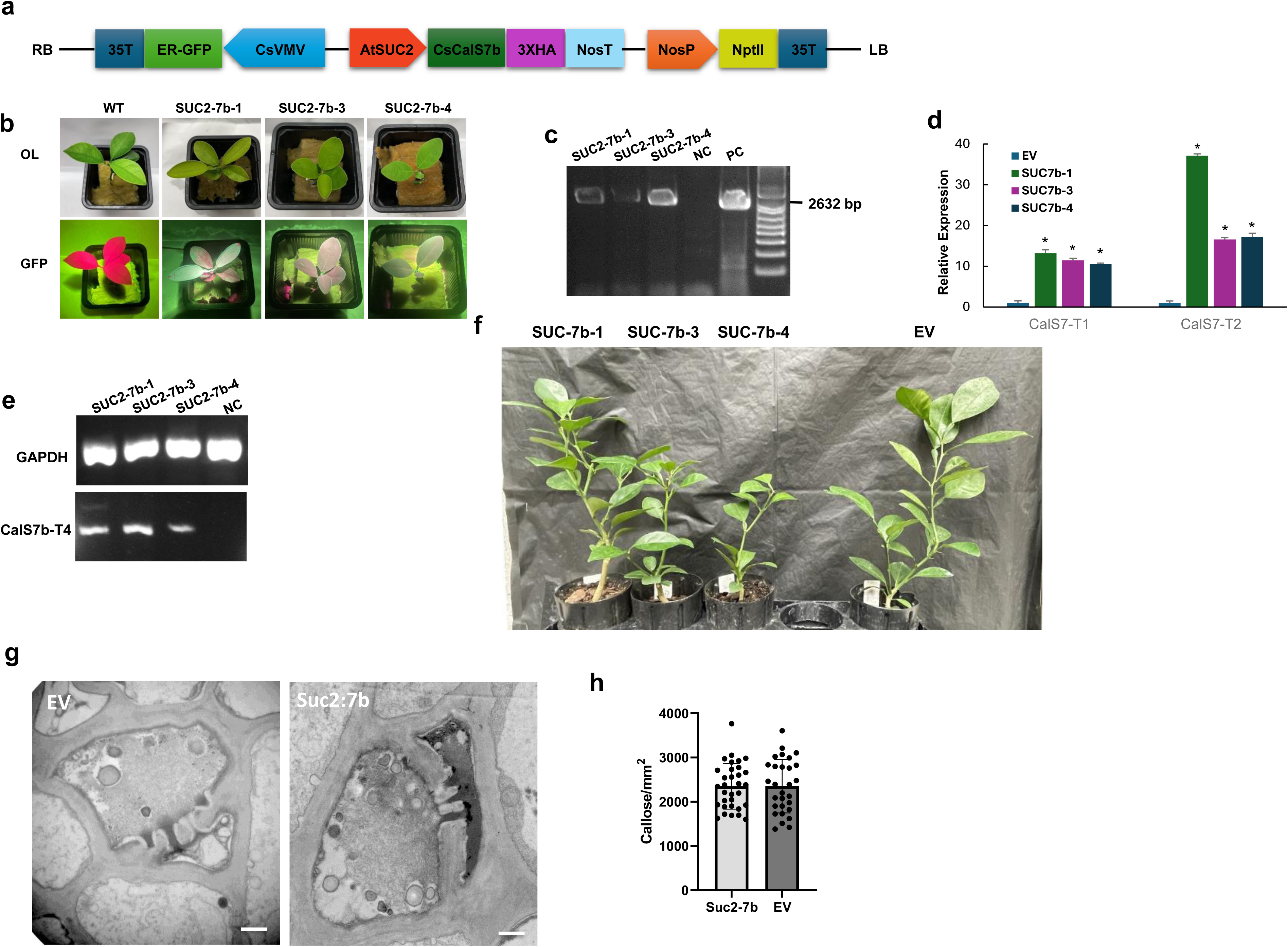
Overexpression of *CsCalS7b* driven by the AtSUC2 promoter in *C. sinensis* cv. Hamlin. **(a)** Illustration of construct used for overexpression of *CsCalS7b* driven by the Suc2 promoter. (**b)** Transgenic Hamlin plants overexpressing *CsCalS7b* driven by the AtSUC2 promoter (SUC-7b). WT, wild-type Hamlin plant without GFP fluorescence signal. Top, under the optical light; bottom, erGFP fluorescence screening positive transgenic citrus under GFP light. (**c)** PCR validation for *CsCalS7b* with the AtSUC2 specific forward primer and gene-specific reverse primer. The plasmid was used as a positive control (PC). The wild type plant was used as a negative control (NC). (**d)** The relative expression level of *CsCalS7b* was tested by RT-qPCR with two pairs of gene-specific primers. *CsGAPDH* was used as an endogenous control. Asterisks (*) indicated statistically significant differences (*P* < 0.05, Student’s two-tailed *t*-test). (**e)** Semi-quantitative PCR using cDNA as the template with *CsCalS7b* forward primers and HA reverse primer. *CsGAPDH* was used as the internal reference. (**f)** *Suc2*:*CalS7b* (SUC-7b) transgenic plants at age of 12 months. (**g)** TEM images showing phloem cells of EV and *Suc2*:*CalS7b* midveins. Scale bars are 1 μm. (**h**) Quantification callose deposition in Suc2-7b and EV stem longitudinal sections.

**Fig. S25:**
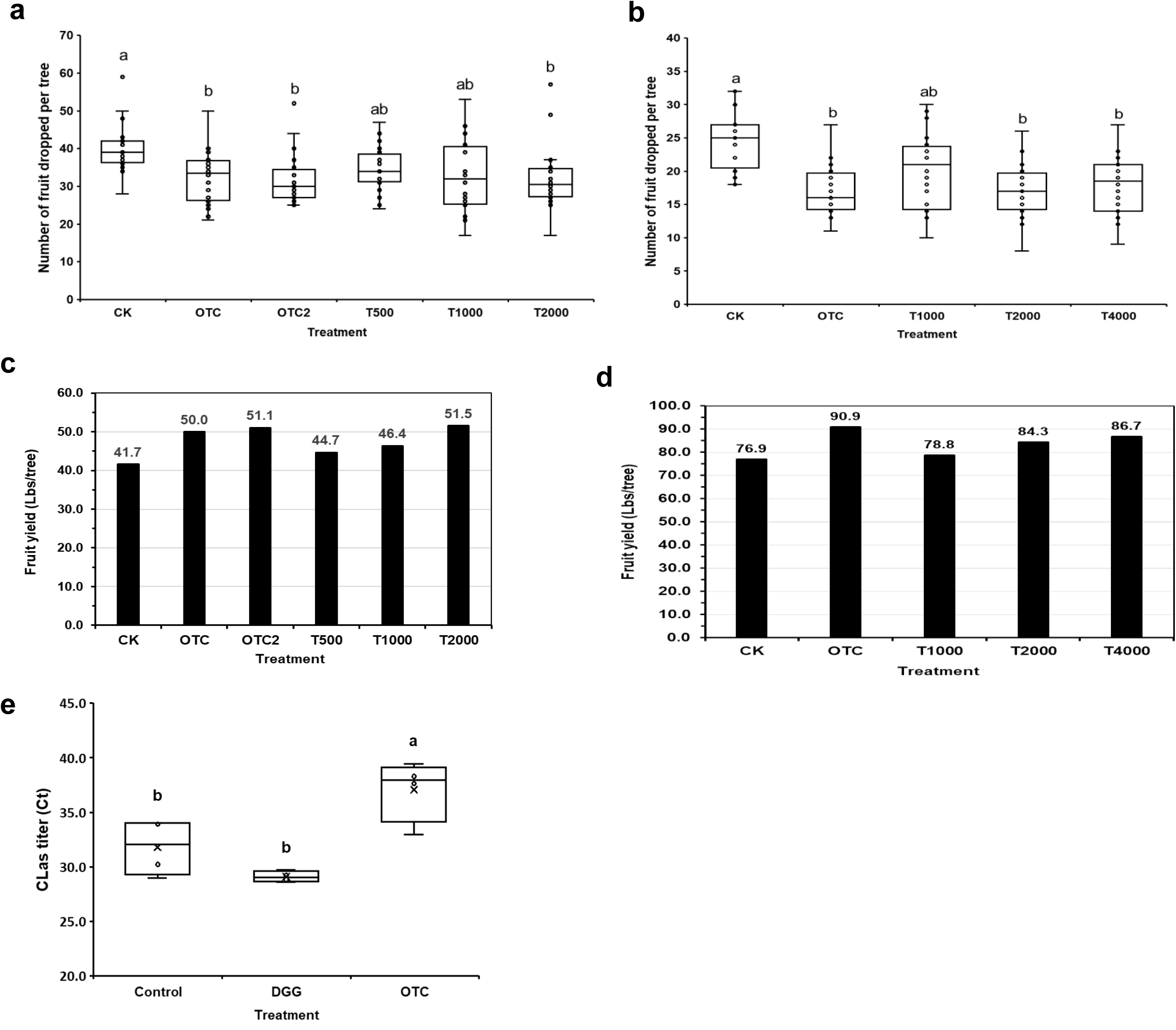
Field trials of DDG treatment on fruit drop and yield of HLB symptomatic trees. (**a**) Pre-harvest fruit drop of 7-year-old Valencia sweet orange trees in a grove at Avon Park, FL. The trees were treated with different treatments by trunk injection at 100 ml per tree in May 2024. (**b**) Pre-harvest fruit drop of 14-year-old Valencia sweet orange trees in a grove at Polk City, FL. The trees were treated with different treatments by trunk injection at 100 ml per tree in July 2024. CK: untreated control. OTC: one injection of 100 ml oxytetracycline (5500 ppm). OTC2: two injections of 50 ml oxytetracycline (5500 ppm) at opposite sides of the tree. T500: 2-Deoxy-D-glucose at 0.5 mM. T1000: 2-Deoxy-D-glucose at 1.0 mM. T2000: 2-Deoxy-D-glucose at 2.0 mM. T4000: 2-Deoxy-D-glucose at 4.0 mM. 20 trees from each treatment (*n* = 20) were examined for fruit drop in February 2025. Statistical analysis was performed using Welch’s analysis of variance and Games-Howell post-hoc test. Different lowercase letters indicate significant difference at *P* < 0.05. Box plots: central line represents the median, box edges delimit lower and upper quartiles, and dots above the boxes are outliers. (**c**) Fruit yield of 7-year-old Valencia sweet orange trees in the experimental grove at Avon Park, FL. (**d**) Fruit yield of 14-year-old Valencia sweet orange trees in the experimental grove at Polk City, FL. (**e**) CLas titers in leaves of fall flush of Valencia sweet orange trees in the treatment of (b). Four trees from each treatment (n = 4) were examined. Statistical analysis was performed using Welch’s analysis of variance and Games-Howell post-hoc test. Different lowercase letters indicate significant difference at P < 0.05. Box plots: central line represents the median, box edges delimit lower and upper quartiles, and X is mean marker.

**Fig. S26:**
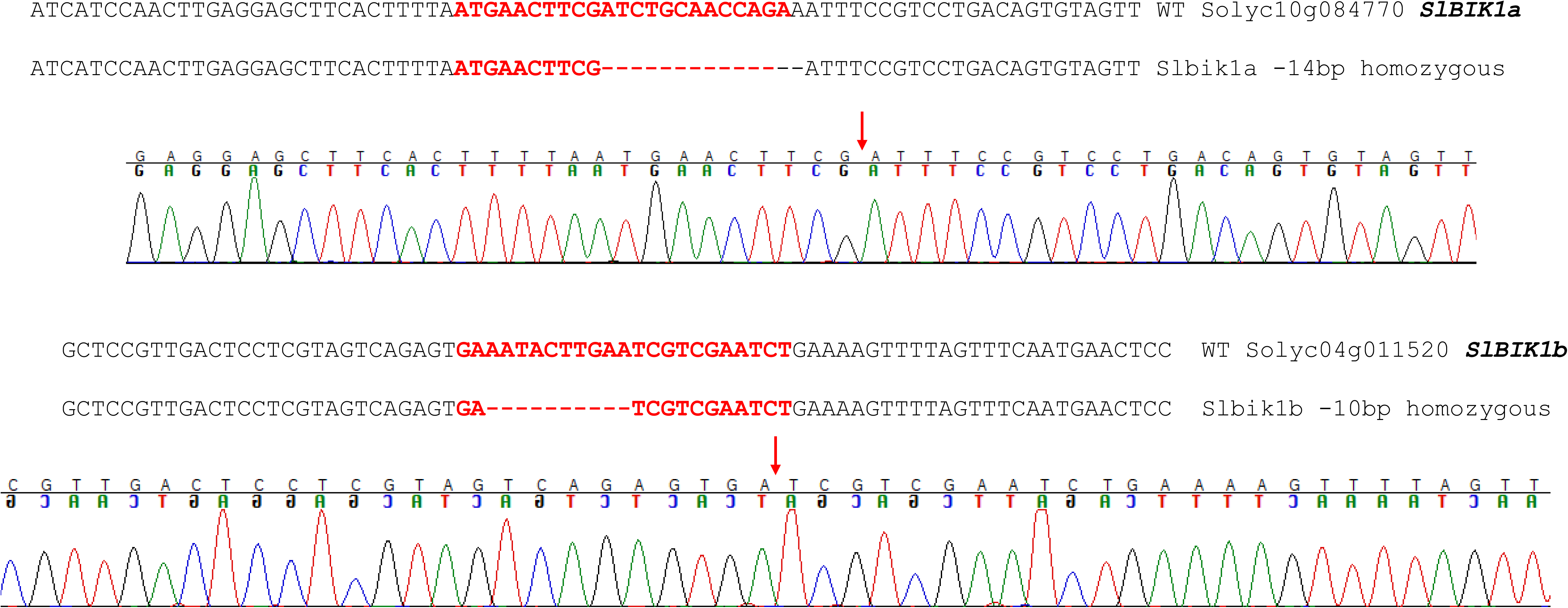
Genotype of *slbik1a/slbik1b* (*SlBIK1a* and *SlBIK1b* double mutant). The crRNA target site used for CRISPR/Cas12a-mediated gene editing is highlighted in red.

**Fig. S27:**
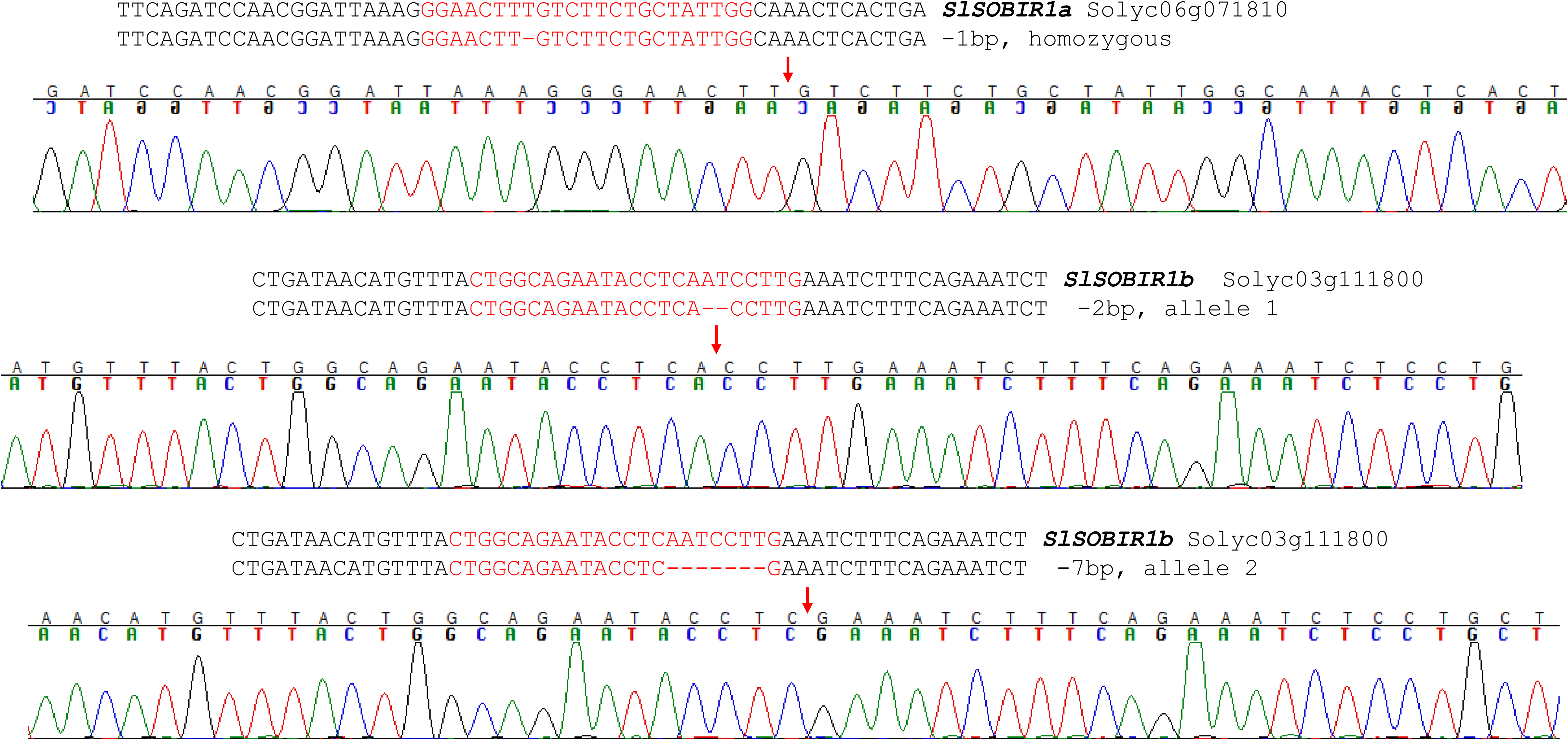
Genotype of *slsobir1a/ slsobir1b* (*SlSOBIR1a* and *SlSOBIR1b* double mutant). The crRNA target site used for CRISPR/Cas12a-mediated gene editing is highlighted in red.

**Fig. S28:**
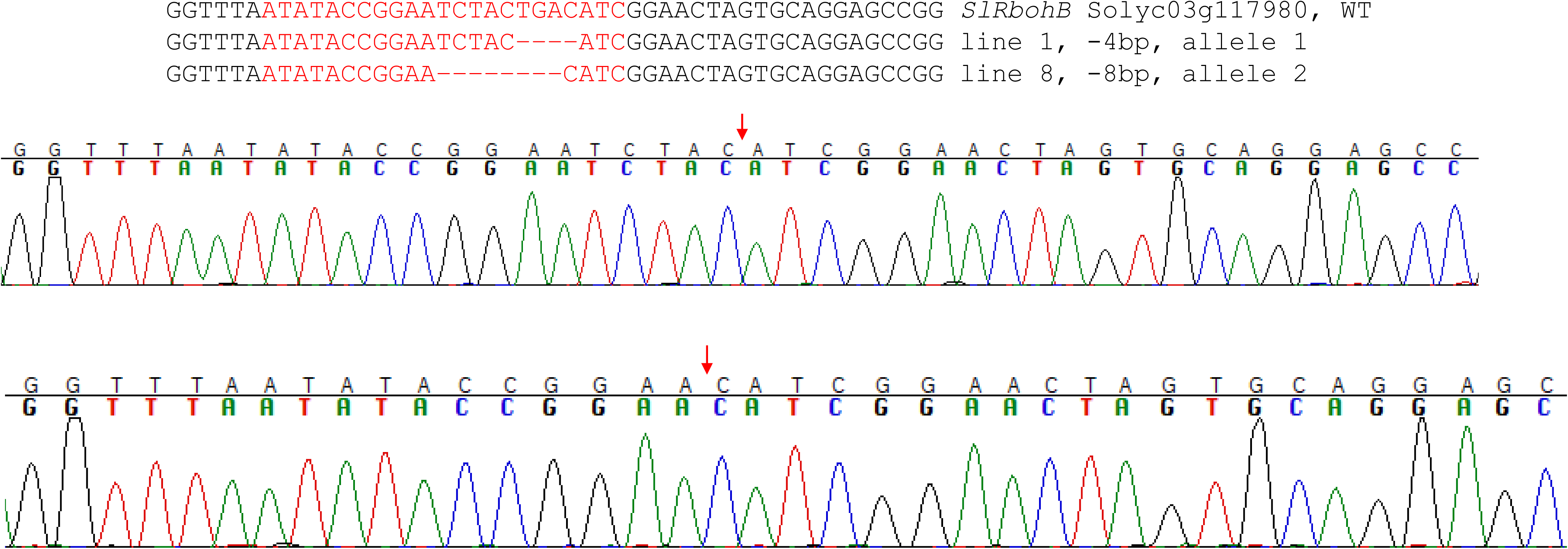
Genotype of *SlRbohB* mutant. The crRNA target site used for CRISPR/Cas12a-mediated gene editing is highlighted in red.

**Fig. S29:**
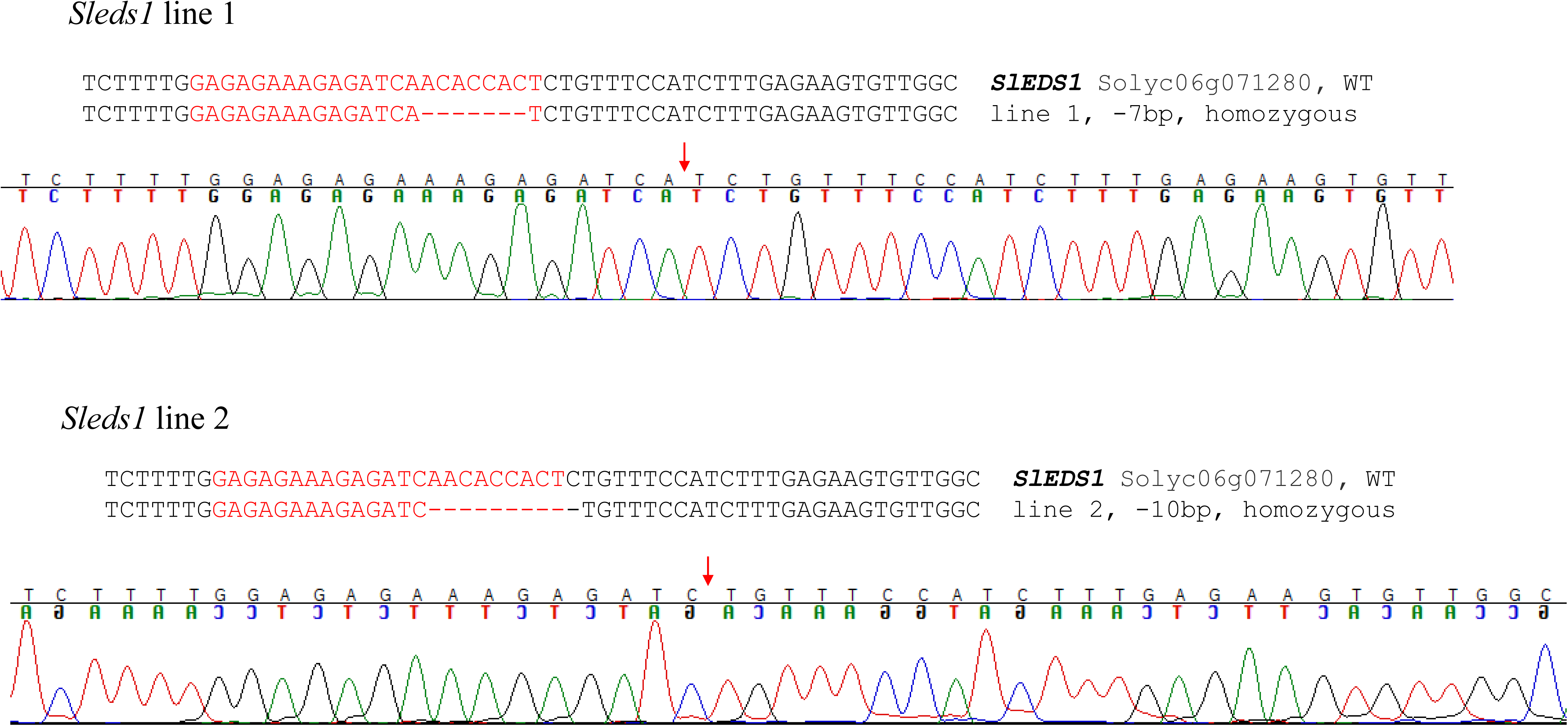
Genotypes of *sleds1* mutants. The crRNA target site used for CRISPR/Cas12a-mediated gene editing is highlighted in red.

**Fig. S30:**
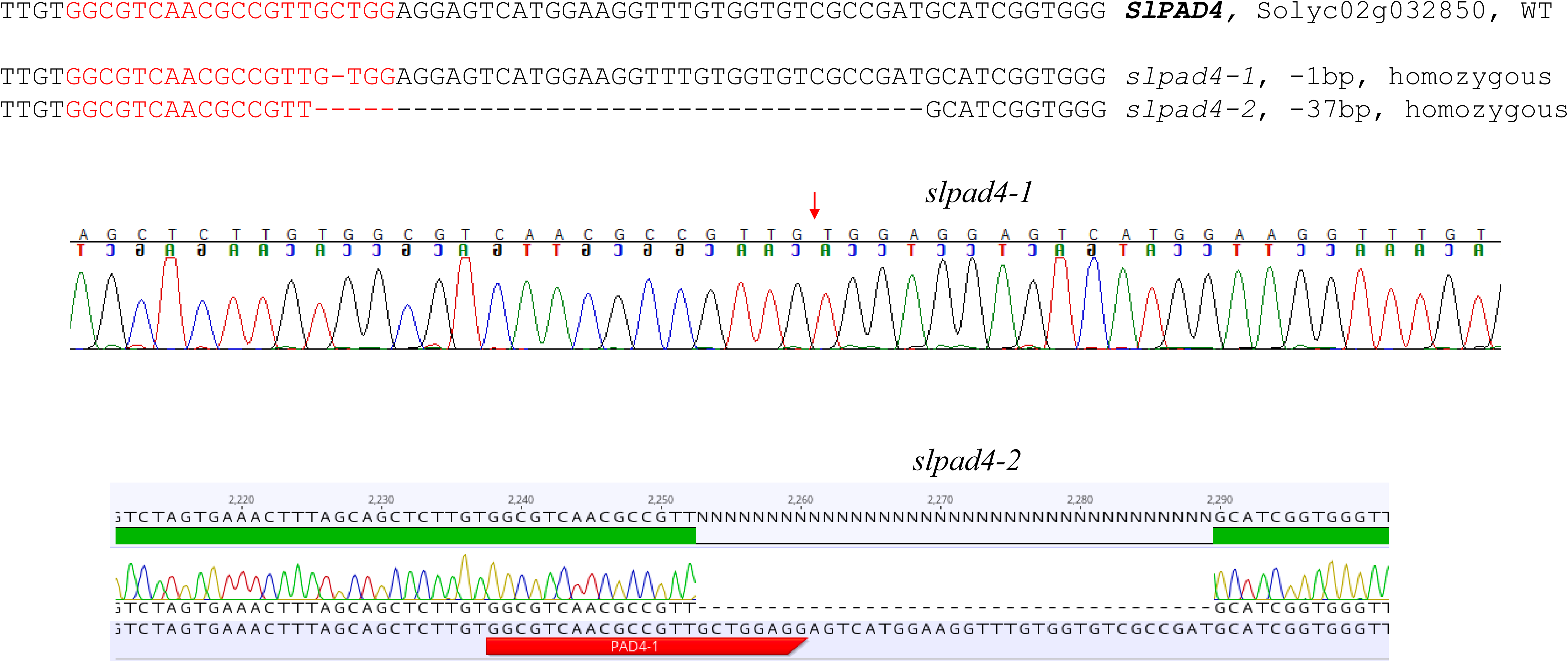
Genotypes of *slpad4* mutants. The gRNA target site used for CRISPR/Cas9-mediated gene editing is highlighted in red.

**Fig. S31.**
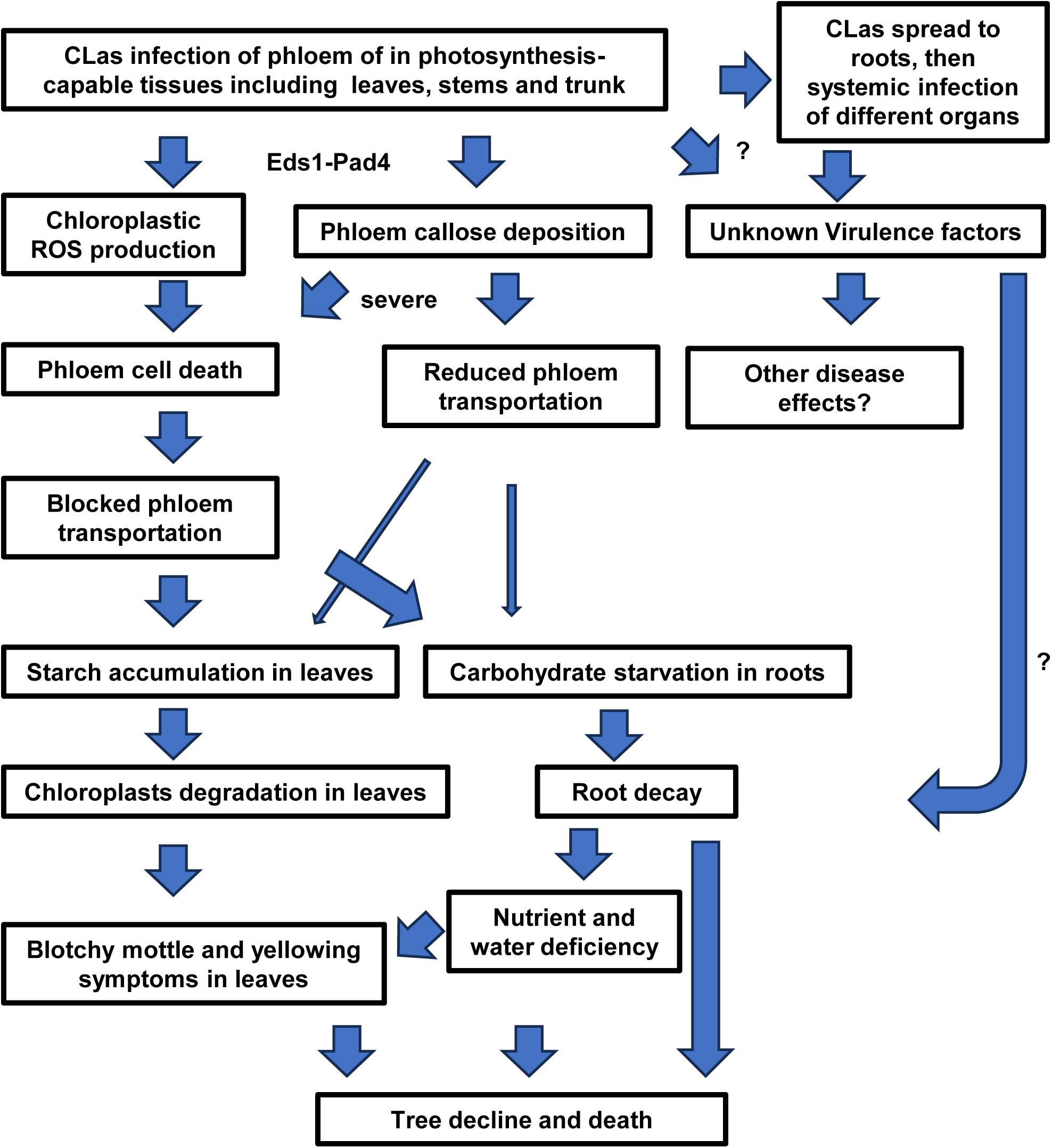
Model of the disease development of the citrus HLB. CLas: *Candidatus* Liberibacter asiaticus. ? indicates unknown.

